# A cell-free biosynthesis platform for modular construction of protein glycosylation pathways

**DOI:** 10.1101/833806

**Authors:** Weston Kightlinger, Katherine E. Duncker, Ashvita Ramesh, Ariel H. Thames, Aravind Natarajan, Allen Yang, Jessica C. Stark, Liang Lin, Milan Mrksich, Matthew P. DeLisa, Michael C. Jewett

## Abstract

Glycosylation plays important roles in cellular function and endows protein therapeutics with beneficial properties. However, constructing biosynthetic pathways to study and engineer protein glycosylation remains a bottleneck. To address this limitation, we describe a modular, versatile cell-free platform for <u>glyco</u>sylation <u>p</u>athway assembly by rapid *in vitro* <u>m</u>ixing and <u>e</u>xpression (GlycoPRIME). In GlycoPRIME, crude cell lysates are enriched with glycosyltransferases by cell-free protein synthesis and then glycosylation pathways are assembled in a mix-and-match fashion to elaborate a single glucose priming handle installed by an *N*-linked glycosyltransferase. We demonstrate GlycoPRIME by constructing 37 putative protein glycosylation pathways, creating 23 unique glycan motifs. We then use selected pathways to design a one-pot cell-free system to synthesize a vaccine protein with an α-galactose motif and engineered *Escherichia coli* strains to produce human antibody constant regions with minimal sialic acid motifs. We anticipate that our work will facilitate glycoscience and make possible new glycoengineering applications.

## Introduction

Glycosylation, the enzymatic process that attaches oligosaccharides to amino acid side chains in proteins, is one of the most abundant, complex, and important post-translational modifications found in nature^1, 2^. Glycosylation plays critical roles in human biology and disease^1^. It is also present in over 70% of approved or preclinical protein therapeutics^3^, having profound effects on stability^4, 5^, immunogenicity^6, 7^, and protein activity^8^. The importance of glycosylation and recent findings that the intentional manipulation of glycans can improve protein therapeutic properties^4, 6, 8^ have motivated many efforts to study and engineer protein glycosylation structures^9–12^.

Unfortunately, the study of glycan functions in natural systems and the engineering of protein glycosylation structures for desired properties remain constrained by multiple factors. One constraint is that glycosylation structures are difficult to control within complex cellular environments because glycan biosynthesis is not template driven^1, 13, 14^. Glycans are synthesized by the coordinated activities of many glycosyltransferases (GTs) across several subcellular compartments^1^, leading to heterogeneity and complicating engineering efforts^13–15^. Another challenge is the shortage of methods for modular assembly of biosynthetic pathways to rapidly access a diversity of glycan structures. New cell lines must be developed to test each new glycosylation pathway^11, 12^, and essential biosynthetic pathways in eukaryotic organisms constrain the ability to modularly build synthetic pathways towards any user-defined glycosylation structure^9, 16^. Furthermore, the efficient and controlled conjugation of glycans onto proteins outside of natural systems remains challenging^13, 15^. A key issue for biochemical approaches is that many of the most important components of protein glycosylation pathways are associated with cellular membranes^13^. Most notably, the engineering of asparagine (*N*-linked) glycosylation has generally relied on the use of oligosaccharyltransferases (OSTs) to transfer prebuilt sugars from lipid-linked oligosaccharides (LLOs) onto proteins as they are transported across membranes. OSTs are integral membrane proteins that often contain multiple subunits^17^ and LLOs are difficult to synthesize and manipulate *in vitro*^13^. Despite recent advances enabling the production, characterization, and use of OSTs in cell-free systems^18–20^, the complexity of these membrane-associated components still presents a major barrier for glycoengineering and the facile construction of multienzyme glycosylation pathways *in vitro*^13, 15^.

A recently discovered class of cytoplasmic, bacterial enzymes known as *N*-linked glycosyltransferases (NGTs), may overcome these limitations by enabling the construction of simplified, fully soluble glycosylation systems in prokaryotic hosts (*e.g.*, *Escherichia coli*)^9, 21–23^. NGTs efficiently install a glucose residue onto asparagine side-chains within proteins using a uracil-diphosphate-glucose (UDP-Glc) sugar donor^24^. Importantly, NGTs are soluble enzymes that can be easily expressed functionally in the *E. coli* cytoplasm^22, 25, 26^. Because glycosylation systems using NGT for glycan-protein conjugation do not require protein transport across membranes or lipid-associated components, they have elicited great interest from the glycoengineering community for the production of recombinant protein therapeutics and vaccines^9, 21, 22, 26–29^. Several recent advances set the stage for this vision. First, the acceptor specificity of NGTs has been extensively studied using peptide and protein substrates^25, 28^, glycoproteomic studies^25^, and the GlycoSCORES technique^26^. These studies revealed that NGTs modify N-X-S/T amino acid acceptor motifs resembling those in eukaryotic glycoproteins. Second, the NGT from *Actinobacillus pleuropneumoniae* (ApNGT) has been shown to modify both native and rationally designed glycosylation sites within eukaryotic proteins and peptides *in vitro* and in the *E. coli* cytoplasm^21, 22, 25, 26^. Third, the Aebi group recently reported a biosynthetic method in *E. coli* cells for elaborating the *N*-linked glucose installed by ApNGT to a polysialic acid motif, which may prolong the serum-half-life of small proteins^21^. Chemoenzymatic methods to transfer pre-built, oxazoline-functionalized oligosaccharides onto the ApNGT-installed glucose residue have also been reported^27, 29^. However, other biosynthetic pathways to build therapeutically relevant glycans using NGTs have not been explored^9^, leaving much of the vast theoretical utility of this enzyme class for biocatalysis inaccessible. Unfortunately, current techniques for building and testing glycosylation systems remain time-consuming because they require the synthesis of new genetic constructs, cellular transformation and expression, and recovery and analysis of glycoproteins from living cells to test each biosynthetic pathway^11, 12^.

One promising alternative to bypass current technical bottlenecks is the use of cell-free systems in which the production of proteins and metabolites occurs *in vitro* without using intact, living cells^30, 31^. Complementing efforts in cellular engineering, *E. coli*-based cell-free protein synthesis (CFPS) systems can achieve gram per liter titers of diverse proteins in hours^30, 32^ and have been shown to enable the rapid discovery, prototyping, and optimization of biosynthetic pathways without the need to reengineer an organism for each pathway iteration^31, 33, 34^. By shifting the design-build-test unit from a genetic construct to a lysate, this approach allows for the construction of glycosylation pathways in hours^18–20, 26^. We recently reported the development of a one-pot <u>c</u>ell-free <u>g</u>lyco<u>p</u>rotein <u>s</u>ynthesis (CFGpS) system to synthesize glycoproteins *in vitro* by overexpressing glycan biosynthesis pathways and OSTs in the *E. coli* chassis strain before lysis^18^. This CFGpS technique represents a significant simplification of cell-based methods and enables the production of glycoproteins with a range of relevant glycans at higher conversions than those reported *in vivo*^16, 18^. However, because our previous work required the construction and expression of glycan biosynthesis operons in living cells, it was not well suited for the rapid discovery and testing of multienzyme glycosylation pathways *in vitro*.

Here, we describe the development of a modular, cell-free method for <u>glyco</u>sylation <u>p</u>athway assembly by rapid *in vitro* <u>m</u>ixing and <u>e</u>xpression (GlycoPRIME) to construct new biosynthetic pathways for making glycoproteins. In this method, crude *E. coli* lysates are selectively enriched with individual GTs by CFPS and then combined in a mix-and-match fashion to construct and analyze multienzyme glycosylation pathways that can then be translated to biomanufacuring platforms (**Fig. 1**). A key feature of GlycoPRIME is the use of ApNGT to site-specifically install a single *N*-linked glucose primer onto proteins, which can then be elaborated to a diverse repertoire of glycans. To validate GlycoPRIME, we optimized the *in vitro* expression of 24 bacterial and eukaryotic GTs using CFPS and characterized their activity on a model glycoprotein substrate containing an acceptor sequence which can be efficiently glycosylated by ApNGT. We then combined these GTs *in vitro* to create 37 biosynthetic pathways, yielding 23 unique glycan structures composed of between 1-5 core saccharide motifs and longer repeating structures. These glycosylation pathways provide new biosynthetic routes to several useful structures such as an α1-3-linked galactose (αGal) epitope, sialylated glycans, as well as fucosylated and sialylated forms of lactose or poly-*N*-acetyllactosamine (LacNAc) without the need for cellular membranes or membrane-bound components. We demonstrated the utility of GlycoPRIME to inform the design of biosynthetic pathways by synthesizing a protein vaccine candidate with an αGal glycan known to have immunostimulatory properties^6, 7, 35^ in a one-pot CFGpS reaction and the constant region (Fc) of the human immunoglobulin (IgG1) antibody with minimal sialic acid glycan motifs known to modulate protein therapeutic properties^5, 36^ in the *E. coli* cytoplasm. We anticipate that the specific pathways discovered in this study as well as the GlycoPRIME method will provide new opportunities to produce diverse glycoproteins for basic research and biotechnological applications.

**Figure 1:**
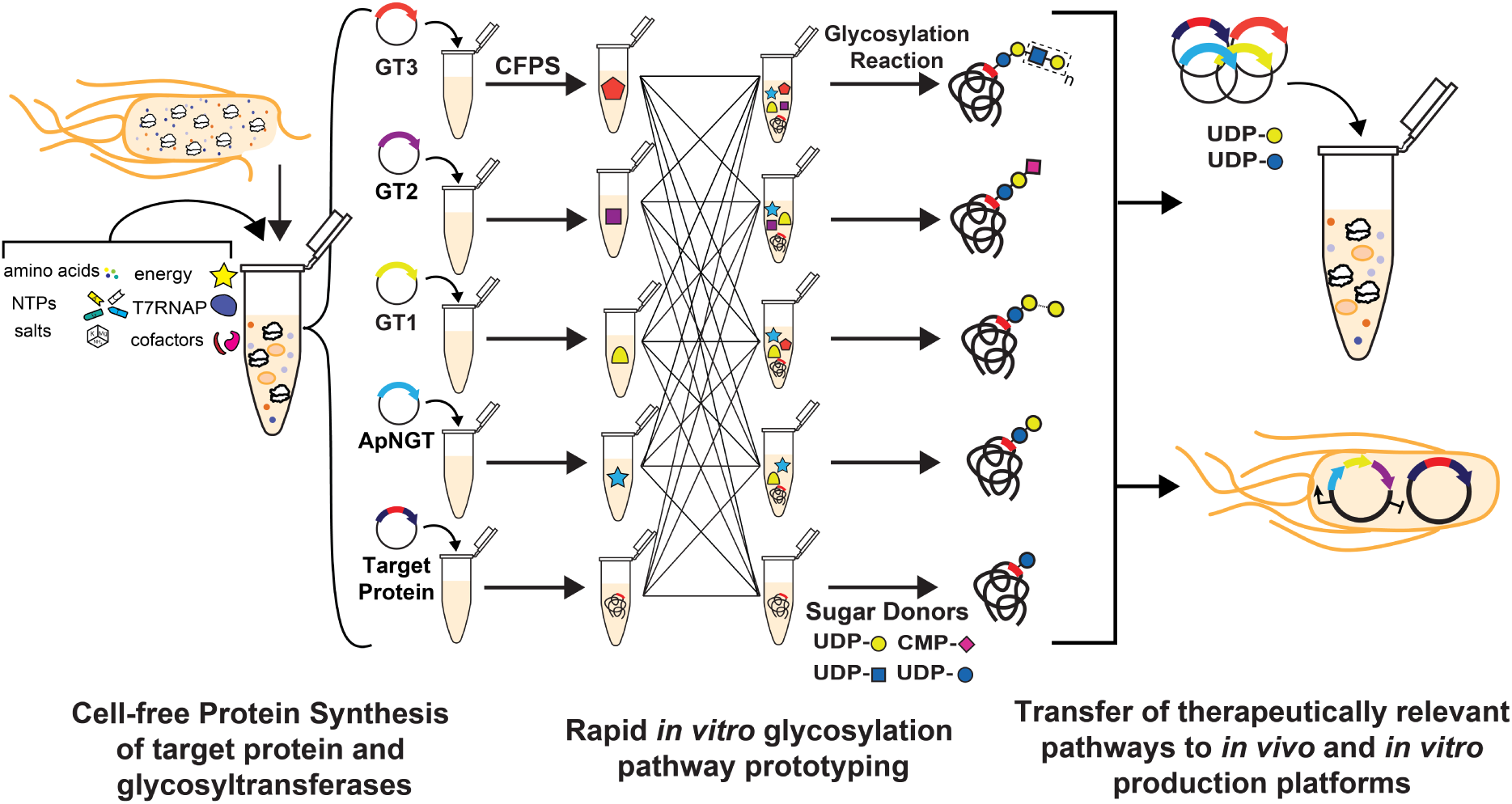
A method for <u>glyco</u>sylation <u>p</u>athway assembly by rapid *in vitro* <u>m</u>ixing and <u>e</u>xpression (GlycoPRIME). GlycoPRIME was established to identify biosynthetic pathways for the construction of diverse *N*-linked glycans. Crude *E. coli* lysates enriched with a target protein or individual glycosyltransferases (GTs) by cell-free protein synthesis (CFPS) were mixed in various combinations to identify biosynthetic pathways for the construction of diverse *N*-linked glycans. A model acceptor protein (Im7-6), the *N*-linked glycosyltranferase from *A. pleuropneumoniae* (ApNGT), and 24 elaborating GTs were produced in CFPS and then assembled with activated sugar donors to identify 23 biosynthetic pathways producing unique glycosylation structures, several with therapeutic relevance. Pathways discovered *in vitro* were transferred to a one-pot glycoprotein synthesis (CFGpS) system or living *E. coli* to produce therapeutically relevant glycoproteins.

## Results

### Establishing an *in vitro* glycosylation pathway engineering platform

To develop GlycoPRIME, we selected ApNGT to install a single *N*-linked glucose priming residue onto a model target protein. The glucose primer could then be elaborated using a modular, multiplexed framework of *in vitro* glycosylation (IVG) reactions to build a diversity of glycosylation motifs (**Fig. 1**). Our vision was to show the versatility of this platform by developing biosynthetic routes to distinct glycoprotein structures including sialylated and fucosylated lactose and LacNAc glycans as well as an αGal epitope.

For proof of concept, we set out to glycosylate a model protein with ApNGT in a setting that would enable our GlycoPRIME workflow. Namely, we needed to identify CFPS conditions that provided high GT expression titers so that the minimum volume of GT-enriched lysate required for complete conversion could be added to each IVG reaction, leaving sufficient reaction volume and generating the substrate for further elaboration by assembly of cell-free lysates. Based on our previous optimization of the ApNGT acceptor sequence^26^, we selected an engineered version of the *E. coli* immunity protein Im7 (Im7-6) bearing a single, optimized glycosylation acceptor sequence of GGNWTT at an internal loop as our model target protein (**Supplementary Table 1** and **Supplementary Note 1**). We used [^14^C]-leucine incorporation to measure and optimize the CFPS reaction temperature for our engineered Im7-6 target and ApNGT (**Supplementary Table 2** and **Fig. 2a**). We found that 23°C provided the optimum amount of soluble product in the trade-off between greater overall protein production at higher temperatures and higher solubility at lower temperatures. We synthesized Im7-6 and ApNGT in CFPS reactions for 20 h and then mixed those protein-enriched crude lysate products together along with 2.5 mM UDP-Glc in a 32-µl IVG reaction and incubated for 20 h at 30°C. We then purified the Im7-6 substrate by affinity purification with Ni-NTA functionalized magnetic beads and performed intact glycoprotein liquid chromatography mass spectrometry (LC-MS) analysis (see **Methods**). We observed the nearly complete conversion of 10 µM of Im7-6 substrate (11 µl) with just 0.4 µM ApNGT (1 µl) (**Fig. 2c**), as indicated by a mass shift of 162 Da (the mass of a glucose residue) in the deconvoluted protein mass spectra (theoretical masses shown in **Supplementary Table 3**). This finding demonstrates that crude lysates enriched by CFPS can be used to assemble IVG reactions without purification and with enough remaining reaction volume to allow for the addition of elaborating GTs.

**Figure 2:**
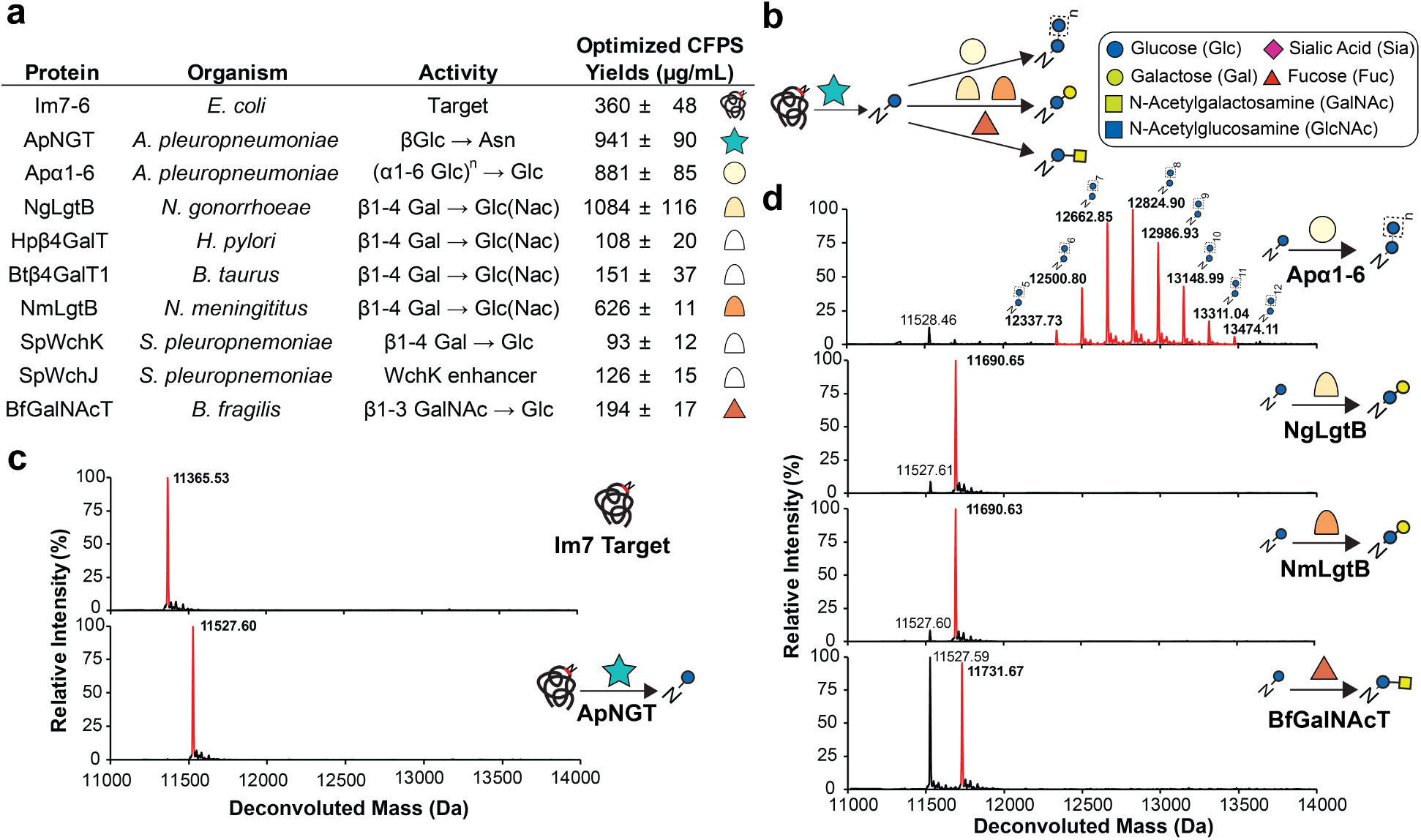
*In vitro* synthesis and assembly of one- and two-enzyme glycosylation pathways. **(a)** Protein name, species, previously characterized activity and optimized soluble CFPS yields for Im7-6 target protein, ApNGT, and GTs selected for glycan elaboration. References for previously characterized activities in **Supplementary Table 4**. CFPS yields and errors indicate mean and standard deviation (S.D.) from n=3 CFPS reactions quantified by [^14^C]-leucine incorporation. Full CFPS expression data in **Supplementary Table 2**. **(b)** Monosaccharide symbol key and *in vitro* glycosylation (IVG) reaction scheme for *N*-linked glucose installation on Im7-6 by ApNGT and elaboration by selected GTs. All glycan structures in this article use Symbol Nomenclature for Glycans (SNFG) and Oxford System conventions for linkages. All mentions of sialic acid refer to *N*-acetylneuraminic acid. **(c)** Deconvoluted mass spectrometry spectra from Im7-6 protein purified from *in vitro* glycosylation (IVG) reactions assembled from CFPS reaction products with and without 0.4 µM ApNGT as well as 2.5 UDP-Glc. Full conversion to *N*-linked glucose was observed after IVG incubation for 24 h at 30°C. **(d)** Intact deconvoluted MS spectra from Im7 protein purified from IVG reactions assembled from CFPS reaction products with 10 µM Im7-6, 0.4 µM ApNGT, and 7.8 µM NmLgtB, 13.9 µM NgLgtB, 3.1 µM BfGalNAcT, or 9.4 µM Apα1-6. IVG reactions were supplemented with 2.5 mM UDP-Glc as well as 2.5 mM UDP-Gal or 5 mM UDP-GalNAc as appropriate for 24 h at 30°C. Observed mass shifts and MS/MS fragmentation spectra (**Supplementary Fig. 1**) are consistent with efficient modification of *N*-linked glucose with β1-4Gal; β1-4Gal; β1-3GalNAc; and α1-6 dextran polymer. Theoretical protein masses found in **Supplementary Table 3**. Spectra from Hpβ4GalT, Btβ4GalT1, and SpWchJ+K, which did not modify the *N*-linked glucose installed by ApNGT are shown in **Supplementary Fig. 2**. All IVG reactions contained 10 µM Im7 and were incubated for 20 h with 2.5 mM of each appropriate nucleotide-activated sugar donor as indicated above. All spectra were acquired from full elution peak areas of all detected glycosylated and aglycosylated Im7-6 species and are representative of n=3 independent IVGs. Spectra from m/z 100-2000 were deconvoluted into 11,000-14,000 Da using Compass Data Analysis maximum entropy method.

We next identified a set of 7 GTs with previously characterized activities that could be useful in elaborating the glucose primer installed by ApNGT to relevant glycans (**Fig. 2** and **Supplementary Table 4**). Previous works indicate that the glucose installed by ApNGT is modified in its native host by the polymerizing Apα1-6 glucosyltransferase to form *N*-linked dextran^24^ and that this structure could be useful as a vaccine antigen^22^. The β1-4 galactosyltransferase LgtB from *Niesseria meningitis* was also shown to form an *N*-linked lactose (Asn-Glcβ1-4Gal) in the previously described synthesis of polysialic acid^21^. Here, we tested whether these pathways could be recapitulated *in vitro* and selected 5 additional enzymes with potentially useful activities (**Fig. 2a**). We chose the *N*-acetylgalactosamine (GalNAc) transferase from *Bacteroides fragilis* (BfGalNAcT) because the GalNAc residue it installs^37^ which could serve as a potential elaboration point for *O*-linked glycan epitope mimics. We also chose a variety of β1-4 galactosyltransferases from *Streptococcus pneumoniae* (SpWchK), *Niesseria gonorrhoeae* (NgLgtB), *Helicobacter pylori* (Hpβ4GalT), and *Bos taurus* (Btβ4GalT1) with similar activities to NmLgtB in order to determine the best biosynthetic route to an *N*-linked lactose structure. This *N*-linked lactose is a critical reaction node for further diversification of the *N*-linked glycan because lactose is a known substrate of a wide variety of GTs that modify milk oligosaccharides and the termini of human *N*-linked glycans^1, 38–41^.

We optimized the *in vitro* expression conditions of these 7 GTs (**Fig. 2** and **Supplementary Table 2**), as well as SpWchJ from *S. pneumoniae*, which is known to enhance the activity of SpWchK. We then assembled IVG reactions by mixing CFPS reaction products containing these GTs with Im7-6 and ApNGT CFPS products as well as UDP-Glc and other appropriate sugar donors according to previously characterized activities (**Fig. 2**). We observed mass shifts of intact Im7-6 and tandem MS (MS/MS) fragmentation spectra of trypsinized glycopeptides consistent with the previously characterized activities of NmLgtB, NgLgtB, BfGalNAcT, and Apα1-6 which are known to install β1-4Gal, β1-4Gal, β1-3GalNac, and an α1-6 dextran polymer, respectively (**Fig. 2**, **Supplementary Fig. 1, and Supplementary Table 5**). We did not observe modification by Hpβ4GalT, SpWchK (even in the presence of SpWchJ), or Btβ4GalT1 (even in the presence α-lactalbumin and conditions conducive to disulfide bond formation) (**Supplementary Fig. 2**). Because NmLgtB and NgLgtB have identical activities, we monitored the resulting glycosylation profile with decreasing amounts of enzyme to find the preferred homolog for *N*-linked lactose synthesis. We found that NmLgtB had greater specific activity than NgLgtB and could achieve nearly complete conversion with 2 µM NmLgtB (**Supplementary Fig. 3**). These results show that multienzyme glycosylation pathways can be rapidly synthesized, assembled, and evaluated in a multiplexed fashion *in vitro*. Using this approach, we found that ApNGT and NmLgtB provide an efficient *in vitro* route to *N*-linked lactose and discovered that ApNGT and BfGalNAcT can site-specifically install a GalNAc-terminated glycan.

### Modular construction of diverse glycosylation pathways

To demonstrate the power of GlycoPRIME for modular pathway construction, we next selected 15 GTs with previously characterized activities that may be able to elaborate the *N*-linked lactose structure installed by ApNGT and NmLgtB into a diverse repertoire of 3-5 saccharide motifs and longer repeating structures (**Fig. 3 and Supplementary Table 4**). Specifically, we targeted structures terminated in sialic acid (Sia) and galactose residues as well as fucosylated and sialylated forms of lactose and LacNAc. We first describe our rationale for selecting these pathway classes and then present our experimental results.

**Figure 3:**
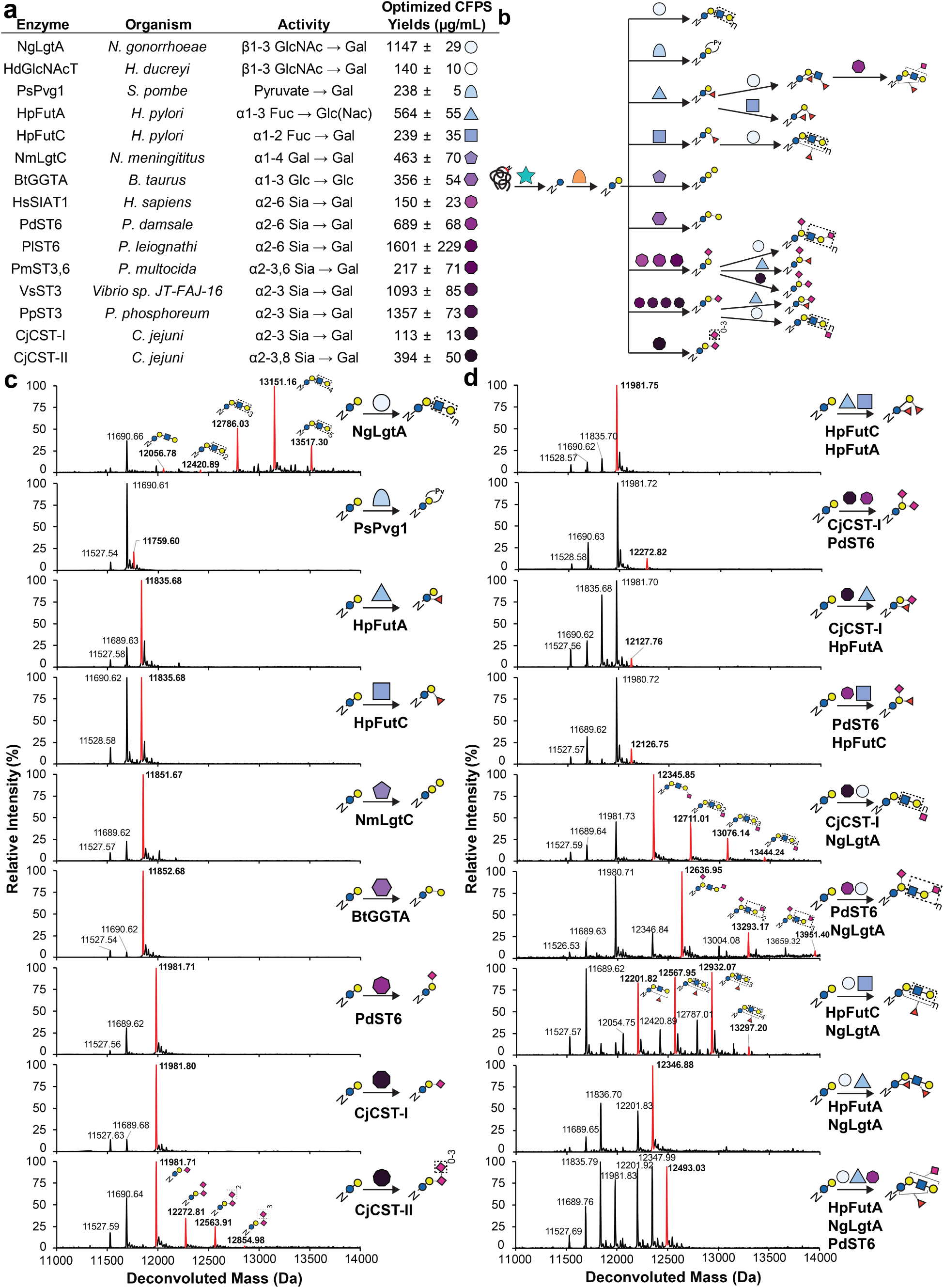
*In vitro* synthesis and assembly of complex glycosylation pathways. **(a)** Protein name, species, previously characterized activity (**Supplementary Table 4**) and optimized CFPS soluble yields (**Supplementary Table 2**) for enzymes tested for elaboration of *N*-linked lactose. CFPS yields and errors indicate mean and S.D. from n=3 CFPS reactions quantified by [^14^C]-leucine incorporation. CjCST-I and HsSIAT1 yields were measured under oxidizing conditions (see **Supplementary Fig. 7**). **(b)** Intact deconvoluted MS spectra from Im7-6 protein purified from IVG reactions with 10 µM Im7-6, 0.4 µM ApNGT, 2 µM NmLgtB, 2.5 mM appropriate sugar donors, and 4.0 µM BtGGTA, 5.3 µM NmLgtC, 4.9 µM HpFutA, 2.6 µM HpFutC, 4.9 µM PdST6, 5.0 µM CjCST-II, 1.3 µM CjCST-I, 11.5 µM NgLgtA, or 2.2 µM SpPvg1. Mass shifts of intact Im7-6, fragmentation spectra of trypsinized Im7-6 glycopeptides (**Supplementary Fig. 5**), and exoglycosidase digestions (**Supplementary Figs. 8 and 9**) are consistent with modification of *N*-linked lactose with α1-3Gal; α1-4Gal; α1-3 Fuc; α2-6 Sia; α2-3 Sia and α2-8 Sia; β1-3 GlcNAc, and pyruvylation according to known GT activities of BtGGTA, NmLgtC, HpFutA, HpFutC, PdST6, CjCST-II, CjCST-I, NgLgtA, or SpPvg1. **(d)** Deconvoluted intact Im7-6 spectra of fucosylated and sialylated LacNAc structures produced by four and five enzyme combinations. IVG reactions contained 10 µM Im7-6, 0.4 µM ApNGT, 2 µM NmLgtB, appropriate sugar donors and indicated GTs at half or one third the concentrations indicated in **b** for four and five enzyme pathways, respectively. Intact mass shifts and fragmentation spectra (**Supplementary Fig. 10**) are consistent with fucosylation and sialylation of LacNAc core according to known activities. Intact protein and glycopeptide fragmentation spectra from other screened GTs and GT combinations not shown here are found in **Supplementary Figs. 4-6 and 10-12.** To provide maximum conversion, IVG reactions were incubated for 24 h at 30°C, supplemented with an additional 2.5 mM sugar donors and incubated for 24 h at 30°C. Spectra were acquired from full elution areas of all detected glycosylated and aglycosylated Im7 species and are representative of at least n=2 IVGs. Spectra from m/z 100-2000 were deconvoluted into 11,000-14,000 Da using Compass Data Analysis maximum entropy method.

We aimed to construct glycans terminated in sialic acids because such structures are known to effectively modulate the trafficking, stability, and pharmacodynamics of glycoprotein therapeutics^5, 8, 21, 36, 42^. As the linkage details of terminal sialic acids are also important^5^, we selected enzymes to install Sia with α2-3, α2-6, and α2-8 linkages onto the *N*-linked lactose. To start, we chose to build a 3’-sialyllactose (Glcβ1-4Galα2-6Sia) structure. This structure could provide several useful properties including mimicry of the GM3 ganglioside (ceramide-Glcβ1-4Galα2-6Sia) for vaccines against cancer cells^43^. The 3’-sialyllactose structure may also mimic the recently reported GlycoDelete structure (GlcNAcβ1-4Galα2-3Sia) which has been shown to simplify the glycosylation patterns of granulocyte-macrophage colony-stimulating factor (GM-CSF) and IgG therapeutics while preserving their activities and *in vivo* circulation times^44^. To build 3’-sialyllactose, we chose four α2-3 sialyltransferases from *Pasteurella multocida* (PmST3,6), *Vibrio sp JT-FAJ-16* (VsST3), *Photobacterium phosphoreum* (PpST3), and *Campylobacter jejuni* (CjCST-I). Next, we aimed to produce 6’-sialyllactose (Glcβ1-4Galα2-6Sia) structure because *N*-glycans bearing terminal α2-6Sia are common in secreted human proteins^12^, have exhibited anti-inflammatory properties^8^, and would provide distinct siglec, lectin, and receptor binding profiles^5^. To produce 6’-sialyllactose we selected three α2-6 sialyltranferases from humans (HsSIAT1), *Photobacterium damselae* (PdST6), and *Photobacterium leiognathid* (PlST6). Finally, we investigated the *in vitro* production of glycans with α2-8Sia that may mimic the GD3 ganglioside (ceramide-Glcβ1-4Galα2-3Siaα2-8Sia), a possible vaccine epitope against human melanoma cells^45, 46^. Based on previous works that used CjCST-II to construct polysialic acids^21, 42^, we selected the CST-II bifunctional sialyltranferase from *C. jejuni* to install terminal α2-8Sia. In addition to these sialyltranferases, we also explored the synthesis of glycans terminated with pyruvate moieties because these structures display similar lectin-binding properties to sialic acids^47^. For this purpose, we selected a recently discovered pyruvyltransferase from *Schizosaccharomyces pombe* (SpPvg1)^47^.

Beyond structures terminated in sialic acids, we also explored pathways that could modify *N*-linked lactose with galactose, fucose (Fuc), and LacNAc motifs. For example, we aimed to engineer a first-of-its-kind bacterial system for complete biosynthesis of proteins modified with αGal (Glcβ1-4Galα1-3Gal) epitopes. αGal is an effective self:non-self discrimination epitope in humans and an estimated 1% of the human IgG pool reacts with this motif^6, 7, 35, 48^. As such, αGal has been shown to confer adjuvant properties when associated with various peptide, protein, whole-cell, and nanoparticle-based immunogens^6, 7, 35, 48–50^. To build the desired αGal pathway, we selected the α1,3 galactosyltransferase from *B. taurus* (BtGGTA). We also selected the globobiose structure (Glcβ1-4Galα1-4Gal), which mimics the Gb3 ganglioside (ceramide-Glcβ1-4Galα1-4Gal) used for the detection^51, 52^ and removal^53^ of *Shigella* toxins that are secreted by pathogenic bacteria and preferentially bind to Gb3^53^. We selected the galactosyltransferase LgtC from *N. meningitis* (NmLgtC) to synthesize this structure. An additional pathway class we sought to build were LacNAc polymers, which regulate cell-cell interactions in humans and are often found on aggressive cancer cells^54^. We chose to test two β1-3 *N*-acetylglucosamine (GlcNAc) transferases from *N. gonorrhoeae* (NgLgtA) and *Haemophilus ducreyi* (HdGlcNAcT) for their ability to make this structure. Finally, we aimed to build fucosylated lactose and LacNAc structures (which may find applications in mimicking properties of Lewis antigens and Helminth immunomodulatory glycans^55^) by testing α1,3 and α1,2 fucosyltransferases from *H. pylori* (HpFutA and HpFutC, respectively). In total, we sought to build 9 oligosaccharide structures by elaborating the *N*-linked lactose installed by ApNGT and NmLgtB.

Following our pathway design and GT selection, we used GlycoPRIME to synthesize and assemble three enzyme biosynthetic pathways containing ApNGT, NmLgtB, and each of the 15 GTs described above. We optimized the expression of each GT in CFPS (**Figure 3a and Supplementary Table 2**) and assembled IVG reactions by combining crude lysates containing each GT with lysates containing Im7-6, ApNGT, and NmLgtB as well as appropriate sugar donors according to their previously established activities. When reaction products were purified by Ni-NTA magnetic beads and analyzed by LC-MS(/MS), we observed mass shifts of intact Im7-6 (**Fig. 3 and Supplementary Fig. 4**) and fragmentation spectra of trypsinized glycopeptides (**Supplementary Fig. 5**) consistent with the modification of the *N*-linked lactose installed by ApNGT and NmLgtB according to the hypothesized activities of all α2-3 sialyltranferases (CjCST-I, and PpST3, VsST3, PmST3,6); all α2-6 sialyltranferases (PdST6, HsSIAT1, and PlST6); the bifunctional α2-3,8 sialyltranferase (CjCST-II); the α1-3 galactosyltransferase (BtGGTA); the α1-galactosyltransferase (NmLgtC); the α1-3 fucosyltransferase (HpFutA); the α1-2 fucosyltransferase (HpFutC), the pyruvyltransferase (SpPvg1), and one β1-3 *N*-acetylglucosaminyltransferase (NgLgtA). We did not observe activity from HdGlcNAcT (**Supplementary Fig. 6**). While we did detect low activity from all sialyltranferases either by intact protein or glycopeptide analysis, we found that CjCST-I and PdST6 provided the highest conversion of all α2-3 and α2-6 sialyltranferases, respectively (**Supplementary Fig. 4**). This optimization demonstrates the ability of GlycoPRIME to quickly compare several biosynthetic pathways to determine the enzyme combinations and conditions that yield the highest conversion to desired products. We also found that we could significantly increase the activity of CjCST-I and HsSIAT1 by conducting CFPS in oxidizing conditions (**Supplementary Fig. 7**), showing the utility of using the open reaction environment of CFPS reactions to improve enzyme synthesis conditions, including the synthesis of a human enzyme with disulfide bonds (HsSIAT1). Notably, we also found that NgLgtA not only installed a β1-3 *N*-acetylglucosamine, but also worked in turn with NmLgtB to form a polymeric LacNAc structure of up to 6 repeat units (**Fig. 3**). We performed digestions of Im7-6 modified by ApNGT, NmLgtB, and PdST6, HsSIAT1, CjCST-I, HpFutA, HpFutC, NgLgtA, and BtGGTA using commercially available exoglycosidases (**Supplementary Figs. 8 and 9**). Our findings support the previously established linkage specificities of these enzymes (**Figs. 2 and 3 and Supplementary Table 4**). Under these conditions, we found that PmST3,6 exhibited primarily α2-3 activity, which is consistent with previous reports^56^.

Having demonstrated the activity of diverse GTs using three enzyme pathways, we pushed the GlycoPRIME system further to evaluate biosynthetic pathways containing four and five enzymes. We aimed to produce sialylated and fucosylated lactose and LacNAc structures using combinations of HpFutA, HpFutC, CjCST-I, PdST6, and NgLgtA. While some combinations of these GTs have been used to create free oligosaccharides or glycolipids^38–41, 57–59^, the potential products of different enzyme combinations resulting from their interacting specificities have not been systematically studied in the context of a protein substrate. We tested all pairwise combinations of these 5 GTs using the GlycoPRIME system by expressing each of them in separate CFPS reactions and then mixing two of those crude lysates in equal volumes with CFPS reactions containing 10 µM Im7-6, 0.4 µM ApNGT, and 2 µM NmLgtB. Upon analysis of these IVG products, we found intact protein (**Fig. 3d**) and glycopeptide fragmentation products (**Supplementary Fig. 10**) indicating the synthesis of several interesting structures including difucosylated lactose, disialylated lactose, lactose variants with combinations of sialylation and fucosylation linkages, sialylated LacNAc structures with branching or only terminal sialic acids, and fucosylated LacNAc structures. This analysis also revealed some possible specificity conflicts between the enzymes. For example, the combinations of CjCST-I with HpFutA and PdST6 with HpFutC yielded products which were both sialyllated and fucosylated, but PdST6 with HpFutC and CjCST-I with HpFutC did not (**Supplementary Fig. 11**). Furthermore, we observed that in the combination of HpFutC with NgLgtA, only one fucose is added to the LacNac backbone regardless of its length (**Fig. 3d** and **Supplementary Fig. 10**). In contrast, when HpFutA and NgLgtA are combined, our observations suggest that both available Glc(NAc) residues may be modified; however, the shorted polymer length suggests that fucosylation with HpFutA may prohibit the continued growth of the LacNAc chain by NgLgtA (**Fig. 3**). While we focused here on the testing of single-pot reactions to inform applications in biomanufacuring platforms, sequential glycosylation reactions *in vitro* using a similar workflow could be used to further characterize these specificity conflicts. In a final test of the number of biosynthetic nodes the GlycoPRIME system can support, we constructed several five enzyme biosynthetic pathways using NgLgtA, one fucosyltransferase (HpFutA or HpFutC), and one sialyltransferase (CjCST-I or PdST6). While the complexity of these glycans did not allow us to unambiguously assign their structures, their intact protein mass shifts (**Supplementary Fig. 11**) and fragmentation spectra (**Supplementary Fig. 10**) did indicate the construction of LacNAc structures glycans in pathways containing NgLgtA, PdST6, and either HpFutA or HpFutC which were both fucosylated, sialylated (**Figure 3d** and **Supplementary Figs. 10 and 12**). Many of the glycosylation structures synthesized by these four and five enzyme combinations have not been found in nature or described in detail and further study will be required to understand the properties they might provide.

### Translating GlycoPRIME pathways to portable cell-free and bacterial production platforms

Having demonstrated the ability to construct new biosynthetic pathways using our modular, cell-free approach, we set out to transfer therapeutically relevant GlycoPRIME pathways to *in vitro* and *in vivo* production platforms (**Fig. 4**). Our goal was to determine if our newly discovered pathways could be used in different contexts and on different target proteins.

**Figure 4:**
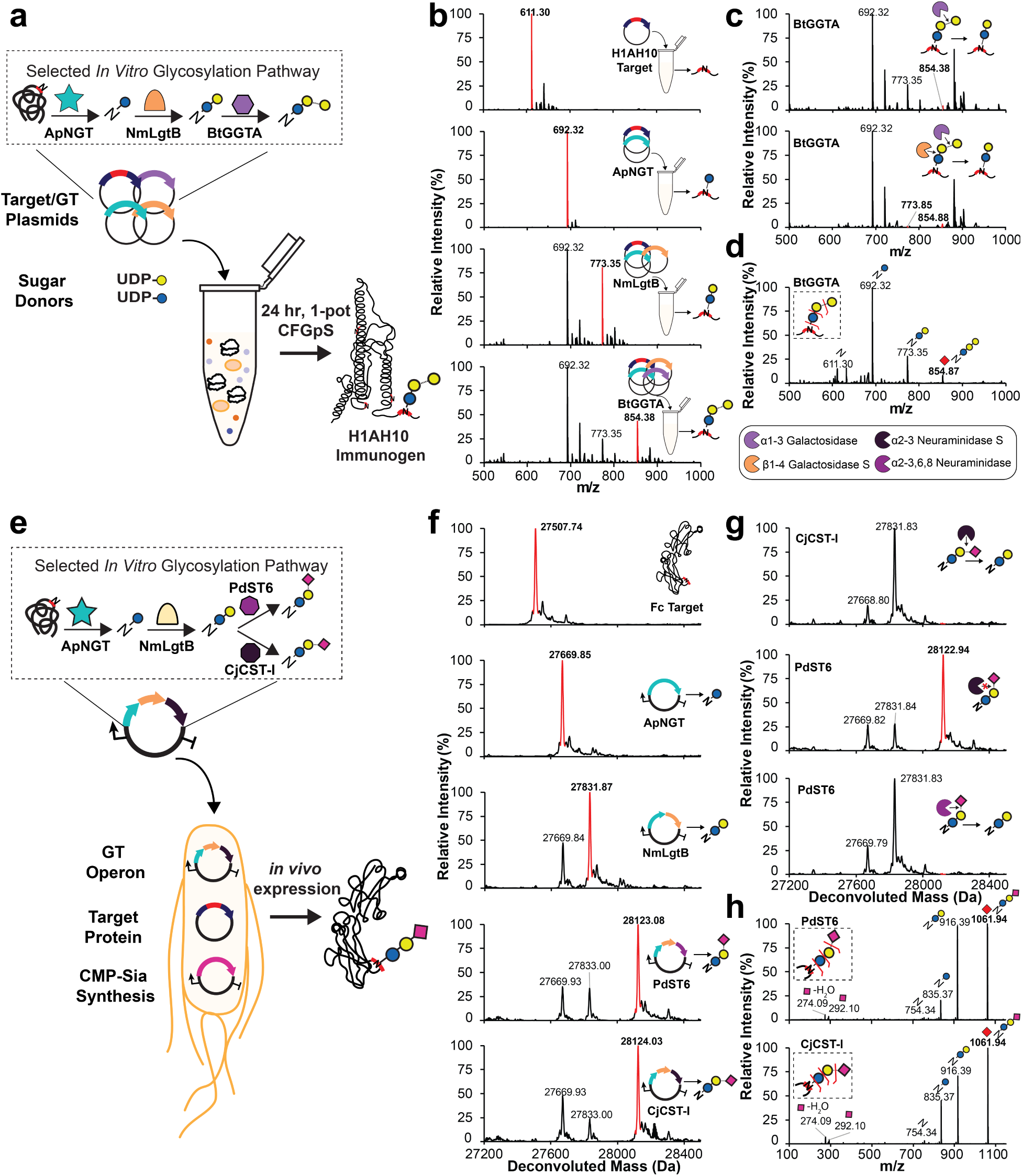
Design of biosynthetic pathways for cell-free and bacterial production platforms. (**a**) One-pot CFGpS for synthesis of H1HA10 protein vaccine modified with αGal glycan. Plasmids encoding the target protein and biosynthetic pathway GTs discovered by GlycoPRIME screening were combined with activated sugar donors in a CFGpS reaction. (**b**) Trypsinized glycopeptide MS spectra, (**c**) exoglycosidase digestions of glycopeptide, and (**d**) MS/MS glycopeptide fragmentation spectra from H1HA10 purified from IVG reactions containing equimolar amounts of each indicated plasmid encoding H1HA10, ApNGT, NmLgtB, and BtGGTA and 2.5 mM of UDP-Glc and UDP-Gal (see see **Methods**). All reactions contained 10 nM total plasmid concentration and were incubated for 24 h at 30°C. The glycopeptide contains one engineered acceptor sequence located at the *N*-terminus of H1HA10. Observed masses and mass shifts in **b-d** spectra are consistent with modification of the H1HA10 peptide with *N*-linked Glc by ApNGT, lactose (Glcβ1-4Gal) by ApNGT and NmLgtB, or αGal epitope (Glcβ1-4Galα1-3Gal) by ApNGT, NmLgtB, and BtGGTA. (**e**) Design of cytoplasmic glycosylation systems to produce sialylated IgG Fc in *E. coli*. Three plasmids containing NmNeuA (CMP-Sia synthesis), IgG Fc engineered with an optimized acceptor sequence (target protein), and biosynthetic pathways discovered using GlycoPRIME (GT operon). (**f**) Deconvoluted intact glycoprotein MS spectra, (**g**) exoglycosidase digestions of intact glycoprotein, and (**h**) MS/MS glycopeptide fragmentation spectra from Fc-6 purified from *E. coli* cultures supplemented with sialic acid, IPTG, and arabinose and incubated at 25°C overnight (see **Methods**). The last GT in all glycosylation pathways is indicated. MS spectra were acquired from full elution areas of all detected glycosylated and aglycosylated protein or peptide species and are representative of n=3 CFGpS or *E. coli* cultures. MS/MS spectra acquired by pseudo Multiple Reaction Monitoring (MRM) fragmentation at theoretical glycopeptide masses (red diamonds) corresponding to detected intact glycopeptide or protein MS peaks using 30 eV collisional energy. Deconvoluted spectra collected from m/z 100-2000 into 27,000-29,000 Da using Compass Data Analysis maximum entropy method. See **Supplementary Tables 5-7** for theoretical masses.

First, we aimed to produce a protein vaccine candidate modified with an αGal epitope in a one-pot CFGpS platform. We selected cell-free biosynthesis as a production platform because this approach has recently generated significant interest for portable, on-demand biomanufacturing using freeze-dried components at the point-of-need^60–63^. In contrast to the cell-free lysate mixing approach used in GlycoPRIME for engineering and discovering new pathways, here we focused on designing a one-pot production platform. In this CFGpS system, the glycosylation target protein is co-expressed with GTs in the presence of sugar donors to simultaneously synthesize and glycosylate the glycoprotein of interest. This method complements our previously reported one-pot CFGpS platform^18^ by synthesizing the glycosylation pathway enzymes *in vitro* rather than within the chassis strain before lysis. We also used commercially available activated sugar donors rather than LLOs, allowing for the testing of multienzyme pathways without the need to produce new chassis strains for each enzyme combination. We validated our one-pot approach by mixing the Im7-6 target protein plasmid, sets of up to three GT plasmids based on 12 successful biosynthetic pathways developed in our two-pot GlycoPRIME screening, and appropriate activated sugar donors in one-pot CFGpS reactions. In all reactions, we observed intact protein mass shifts consistent with the modification of Im7-6 with the same glycans observed in our two-pot system, albeit at lower efficiency (**Supplementary Fig. 13**). After validating our one-pot approach, we set-out to synthesize and glycosylate the recently reported influenza vaccine candidate, H1HA10^64^, with αGal using the biosynthetic pathway discovered by GlycoPRIME screening (**Fig. 4**). We selected H1HA10 as a model protein to demonstrate the αGal pathway because it is an effective immunogen that can be expressed in *E. coli*, and the chemoenzymatic installation of αGal has been shown to act as an intramolecular adjuvant for other influenzae vaccine proteins^7, 65^. When we combined UDP-Glc, UDP-Gal, and plasmids encoding the H1HA10 protein ApNGT, NmLgtB, and BtGGTA in a one-pot CFGpS reaction, we observed the installation of αGal on a tryptic peptide containing an engineered acceptor sequence at the *N*-terminus of the H1HA10 protein (**Fig. 4b**). We further confirmed the linkages of the αGal glycan at this site by digestion with commercially available exoglycosidases and MS/MS (**Fig. 4c-d and Supplementary Table 6**).

The advent of bacterial glycoengineering has presented new opportunities for protein vaccine and therapeutic production^9^. While polysialylated glycoproteins have been produced in engineered *E. coli*^21^, different terminal sialic acid linkages and simplified, more homogeneous glycosylation structures are desirable for some applications of glycoprotein therapeutics^14, 44^. To demonstrate the utility of GlycoPRIME for discovering pathways to manufacture glycoproteins in cells, we designed glycosylation systems to install *N*-linked 3’-sialyllactose and 6’-siallylactose in living *E. coli* (**Fig. 4**). Specifically, we sought to produce the Fc region of human IgG1 modified with terminal sialic acids in α2-3 and α2-6 linkages, which are known to modulate protein stability and therapeutic trafficking^5, 8, 36^. To achieve this goal, we constructed a three-plasmid system composed of a constitutively expressed cytidine-5’-monophospho-*N*-acetylneuraminic acid (CMP-Sia) synthesis plasmid containing the CMP-Sia synthase from *N. meningititus* (ConNeuA), an IPTG-inducible target protein plasmid, and a GT operon plasmid containing ApNGT, NmLgtB, and either CjCST-I or PdST6. The CMP-Sia synthase plasmid is necessary because *E. coli* strains do not endogenously produce CMP-Sia. Following previously reported strategies for production of CMP-Sia in *E. coli*^21, 41, 66^, we selected a K-12 derived *E. coli* strain carrying the NanT sialic acid transporter gene for intake of sialic acid supplemented in the media and knocked out the NanA CMP-Sia aldolase gene to prevent digestion of intracellular sialic acid, yielding CLM24Δ*nanA*. As with CFGpS, we validated the synthesis of our target glycans on the Im7-6 model protein. When we transformed our three-plasmid system into CLM24Δ*nanA* and induced both target protein and GT operon expression, we observed intact protein spectra consistent with the modification of Im7-6 with *N*-linked Glc by ApNGT, the elaboration to lactose by NmLgtB, and the elaboration to 3’-sialyllactose or 6’-siallylactose by CjCST-I or PdST6, respectively (**Supplementary Fig. 14**). To synthesize Fc modified with these glycans, we replaced the Im7-6 target plasmid with a plasmid encoding the human IgG Fc region containing an engineered acceptor site at the conserved Asn297 glycosylation site (Fc-6)^26^. In this system, we observed intact protein MS, MS/MS peptide fragmentation, and exoglycosidase digestions consistent with the expected installation of Glc, lactose, and either 3’-sialyllactose or 6’-sialyllactose onto Fc-6 according to the GT operon supplied (**Fig. 4f-h, Supplementary Fig. 15, and Supplementary Table 7**. Further investigations will be required to assess the efficacy of the αGal epitope as an adjuvant for H1HA10 and the therapeutic effects of minimal sialic acid motifs on Fc. However, our findings clearly demonstrate how the high level of control and modularity afforded by the GlycoPRIME workflow can facilitate the design of biosynthetic pathways to produce diverse glycoproteins in cell-free platforms and the bacterial cytoplasm.

## Discussion

In this work, we established and demonstrated the utility of the GlycoPRIME platform, a cell-free workflow for the modular synthesis and assembly of multienzyme glycosylation pathways. By moving GT production and biosynthetic pathway construction outside of the cell using CFPS and the multiplexed mixing of GT-enriched crude lysates, we were able to rapidly explore 37 putative protein glycosylation pathways, 23 of which successfully produced unique glycosylation motifs. Key to the modular assembly of these pathways was the use of ApNGT, a soluble enzyme easily expressed in bacterial systems, to efficiently install a priming *N*-linked glucose onto glycoproteins. By elaborating this glucose residue, we were able to create diverse glycosylation motifs *in vitro* from the bottom-up that otherwise would have required the engineering of many living cell lines. Of the 23 unique glycosylation motifs produced in this work, many have been synthesized as free^38–41, 57, 58^ or lipid-linked^38, 39^ oligosaccharides or by remodeling existing glycoproteins^6, 27, 42^; however, to our knowledge, only glucose^21, 22, 26^, dextran^22^, lactose^21^, LacNAc^59^, and polysialyllactose^21^ have been previously produced as glycoprotein conjugates in bacterial systems. Therefore, the pathways described in this paper represent a major addition to the repertoire of *N*-linked glycans that can be produced in bacterial glycoprotein engineering platforms. To our knowledge, we achieved the first bacterial biosynthesis of proteins bearing *N*-linked 3’-siallylactose, 6’-siallylactose, the αGal epitope, pyruvylated lactose, 2’-fucosyllactose (Glcβ1-4Galα1-2Fuc), 3-fucosyllactose (Glcβ1-4[α1-3Fuc]Gal), as well as many mono- or di-fucosylated and sialylated forms of lactose or LacNAc. Many of these protein-linked glycans may provide new antigens or desired properties for protein vaccines and therapeutics. For example, αGal is known to be immunostimulatory^6, 7^ and terminal sialic acids can increase protein therapeutic stability and modulate protein trafficking^5, 8, 36^.

An important feature of GlycoPRIME screening is that discovered pathways can be implemented in new contexts and on new proteins for biomanufacuring platforms *in vitro* and in the *E. coli* cytoplasm. Specifically, we demonstrated the production of a candidate vaccine protein, H1AH10, modified with an αGal motif in a one-pot CFGpS reaction and the production of IgG1 Fc region modified with 3’-siallylactose and 6’-siallylactose in living *E. coli* (**Fig. 4**). The use of ApNGT for protein conjugation in the glycosylation pathways described in this work makes them attractive for the production of glycoproteins in bacterial systems because they do not require transport across cellular membranes or membrane-associated components.

Looking forward, GlycoPRIME provides a new way to discover, study, and optimize natural and synthetic glycosylation pathways. For example, future applications (particularly those involving biosynthetic pathways with many GTs) could leverage GlycoPRIME to better understand GT specificities and identify optimal sets of enzymes and enzyme stoichiometry for biosynthesis. By enabling the identification and rapid assembly of enzymes that reliably produce desired glycoproteins, GlycoPRIME is also poised to expand the glycoengineering toolkit and help enable the production of glycoproteins on demand and by design. For example, recent reports showing successful production of ppGalNAcTs^26^ and OSTs^19^ *in vitro* as well as methods to supplement lipid-associated glycans into cell-free synthesis reactions^18–20, 67^ present new opportunities to discover biosynthetic pathways yielding diverse glycosylation structures (*N*- and *O*-linked) with small modifications to the GlycoPRIME workflow. Finally, the diverse glycans accessible by GlycoPRIME pathways could be useful for systematically determining the minimal requirements for desired glycoprotein functions and properties endowed by larger glycans structures.

In summary, we expect that the GlycoPRIME method and new biosynthetic pathways described in this work will quicken the pace of development towards improved glycoprotein therapeutics and make possible new biomanufacturing opportunities.

## Acknowledgements

The authors acknowledge T. Jaroentomeechai, A. Karim, J. Hershewe, and J. Kath for helpful critiques and sharing of reagents as well as S. Habibi, A. Ott, and S. Shafie for assistance with LC-MS instrumentation. This work made use of the IMSERC core facility at Northwestern University, which has received support from the Soft and Hybrid Nanotechnology Experimental (SHyNE) Resource (NSF ECCS-1542205), the State of Illinois, and the International Institute for Nanotechnology (IIN). This material is based upon work supported by the Defense Threat Reduction Agency (HDTRA1-15-10052/P00001), the David and Lucile Packard Foundation, the Dreyfus Teacher-Scholar program, the National Institutes of Health (NIH) and National Institutes of Environmental Health Sciences (NIEHS) through T32 ES007059, and the National Science Foundation through MCB-1413563 and the Graduate Research Fellowship program (DGE-1324585). Its contents are the sole responsibility of the authors and do not necessarily represent the official views the funding agencies above.

## Author contributions

W.K., K.E.D, A.R., and A.H.T designed, performed, and analyzed experiments. A.N. constructed plasmids and assisted with engineered *E. coli* strain design. L.L. and A.Y. conducted early glycosyltransferase activity screens. J.C.S. and M.P.D. assisted with CFGpS system development. M.P.D. and M.M. interpreted the data and helped guide the study. M.C.J. interpreted the data and directed the study. W.K. and M.C.J. conceived of the study and wrote the manuscript.

## Data availability

All data generated or analyzed during this study are included in this article (and its supplementary information) or are available from the corresponding authors on reasonable request.

## Methods

### Plasmid construction and molecular cloning

Details and sources of plasmids used in this study are shown in **Supplementary Table 1** with applicable database accession numbers. Full coding sequence regions with plasmid context are shown in **Supplementary Note 1**. Codon-optimized DNA sequences encoding glycosylation targets and GTs in CFPS were synthesized as gene fragments or intact plasmids by Twist Bioscience, Integrated DNA Technologies, or Life Technologies. Gene fragments were inserted between NdeI and SalI restriction sites in the pJL1^26^ *in vitro* expression vector using Gibson assembly and standard molecular biology techniques^68^. Some GTs were produced with an *N*-terminal CAT-Strep-Linker (CSL) fusion sequence that has been shown to increase *in vitro* expression^26^ (see **Supplementary Note 1**). Plasmids for expression of Im7-6 and Fc-6 glycosylation targets in the CLM24Δ*nanA E. coli* strain were generated by polymerase chain reaction (PCR) amplification of engineered forms of Im7 (Im7-6) and Fc (Fc-6) carrying optimized ApNGT glycosylation acceptor sequences and His-tags from pJL1.Im7-6 and pJL1.Fc-6^26^. These gene fragments were then placed into a pBR322 (ptrc99) backbone^69^ with Carbenicillin resistance and IPTG inducible expression between NcoI and HindIII restriction sites using Gibson assembly. Plasmids for expression of GT operons in *E. coli* were constructed by PCR amplification of ApNGT, NmLgtB, and CjCST-I or PdST6 from their pJL1 plasmid forms followed by Gibson assembly into a pMAF10 backbone^26^ with Trimethoprim resistance, a pBBR1 origin of replication, and arabinose inducible expression between NcoI and HindIII restriction sites. Strep-II tags, FLAG-tags, and ribosome binding sites designed using the RBS Calculator v2.0^70^ for maximum translation initiation rate were inserted into these plasmids as shown in **Supplementary Table 1** and **Supplementary Note 1**. The pCon.NeuA plasmid for production of CMP-Sia in *E. coli* was generated by PCR amplification of NeuA from pTF^71^ followed by Gibson assembly into a pConYCG backbone with Kanamycin resistance and modified with a P32100 promoter for constitutive expression between the NsiI and SalI restriction sites.

### Preparation of cell extracts for CFPS

CFPS of glycosylation enzymes and target proteins was performed using crude *E. coli* lysate from a recently described, high-yielding MG1655-derived *E. coli* strain C321.ΔA.759^32^ prepared using well-established methods^26, 32^. Briefly, 1-liter cultures of *E. coli* cells were grown from a starting OD600 = 0.08 in 2xYTPG media (yeast extract 10 g/l, tryptone 16 g/l, NaCl 5 g/l, K2HPO4 7 g/l, KH2PO4 3 g/l, and glucose 18 g/l, pH 7.2) in 2.5-liter Tunair flasks at 34°C with shaking at 250 r.p.m. Cells were harvested on ice at OD600 = 3.0 and pelleted by centrifugation at 5,000xg at 4°C for 15 min. Cell pellets were washed three times with cold S30 buffer (10 mM Tris-acetate pH 8.2, 14 mM magnesium acetate, 60 mM potassium acetate, 2 mM dithiothreitol [DTT]) before being frozen on liquid nitrogen and then stored at −80°C. Cell pellets were thawed on ice and resuspended in 0.8 ml of S30 buffer per gram of wet cell weight and lysed in 1.4 ml aliquots on ice using a Q125 Sonicator (Qsonica) using three pulses (50% amplitude, 45 s on and 59 s off). After sonication, 4 µl of 1 M DTT was added to each aliquot. Each aliquot was centrifuged at 12,000xg and 4°C for 10 min. The supernatant was incubated at 37°C at 250 r.p.m. for 1 h and centrifuged at 10,000xg at 4°C for 10 min. The clarified S12 lysate supernatant was then frozen on liquid nitrogen and stored at −80°C.

### Cell-free protein synthesis

CFPS of glycosylation targets and GTs was performed using a well-established PANOx-SP crude lysate system^72^. Briefly, CFPS reactions contained 0.85 mM each of GTP, UTP, and CTP; 1.2 mM ATP; 170 µg/ml of *E. coli* tRNA mixture; 34 µg/ml folinic acid; 16 µg/ml purified T7 RNA polymerase; 2 mM of each of the 20 standard amino acids; 0.27 mM coenzyme-A (CoA); 0.33 mM nicotinamide adenine dinucleotide (NAD); 1.5 mM spermidine; 1 mM putrescine; 4 mM sodium oxalate; 130 mM potassium glutamate; 12 mM magnesium glutamate; 10 mM ammonium glutamate; 57 mM HEPES at pH = 7.2; 33 mM phosphoenolpyruvate (PEP); 13.3 µg/ml DNA plasmid template encoding the desired protein in the pJL1 vector; and 27% v/v of *E. coli* crude lysate. *E. coli* total tRNA mixture (from strain MRE600) and phosphoenolpyruvate were purchased from Roche Applied Science. ATP, GTP, CTP, UTP, the 20 amino acids, and other materials were purchased from Sigma-Aldrich. Plasmid DNA for CFPS was purified from DH5-α *E. coli* strain (NEB) using ZymoPURE Midi Kit (Zymo Research). CFPS reactions under oxidizing conditions conducive to disulfide bond formation were performed similarly to standard CFPS reactions except for the use of a 30 minute preincubation of the lysate with 14.3 µM IAM and the addition of 4 mM oxidized L-glutathione GSSG, 1 mM reduced L-glutathione, and 3 µM of purified *E. coli* DsbC to the CFPS reaction^73^. All proteins were expressed in 15 µl batch CFPS reactions in 2.0 ml centrifuge tubes. For GlycoPRIME, CFPS reactions were incubated for 20 h at optimized temperatures for each protein (**Supplementary Table 2**).

### Cell-free glycoprotein synthesis

One-pot, CFGpS was performed similarly to CFPS, except that CFGpS reactions had a total volume of 50 µl and were supplemented with 2.5 mM of each appropriate activated sugar donor as well as multiple plasmid templates from the desired target protein and up to three GTs. CFGpS reactions contained a total plasmid concentration of 10 nM, divided equally between each of the unique plasmids in the reaction. CFGpS reactions were incubated for 24 h at 23°C before purification by Ni-NTA magnetic beads for glycopeptide or intact protein analysis by LC-MS.

### Quantification of CFPS yields

CFPS yields of glycosylation targets and GTs for GlycoPRIME were determined by supplementation of standard CFPS reactions with 10 µM [^14^C]-leucine using established protocols^26, 32^. Briefly, proteins produced in CFPS were precipitated and washed three times using 5% trichloroacetic acid (TCA) followed by quantification of incorporated radioactivity by a Microbeta2 liquid scintillation counter. Soluble yields were determined from fractions isolated after centrifugation at 12,000xg for 15 min at 4°C. Low levels of background radioactivity were measured in CFPS reactions containing no plasmid template and subtracted before calculation of protein yields.

### Assembly and purification from *in vitro* glycosylation reactions

IVG reactions for GlycoPRIME were assembled in standard 0.2 ml tubes from the supernatant of completed CFPS reactions containing the Im7-6 target protein and indicated GTs centrifuged at 12,000xg for 10 min at 4°C. Target and enzyme yields were quantified and optimized by [^14^C]-leucine incorporation (**Supplementary Table 2**). Standard IVG reactions contained 10 µM Im7-6 target, indicated amounts of up to five GTs forming a putative biosynthetic pathway, 10 mM MnCl2 (to provide the preferred metal cofactor for NmLgtB and other GTs), 23 mM HEPES buffer at pH = 7.5, and 2.5 mM of each required nucleotide-activated sugar donor (according to previously characterized activities shown in **Supplementary Table 4**). Each reaction contained a total volume of 32 µl with 25 µl of completed CFPS reactions (when necessary, the remaining CFPS reaction volume was filled by a completed CFPS reaction which had synthesized sfGFP). After assembly, IVG reactions containing up to two GTs were incubated for 24 h at 30°C. To increase conversion, IVG reactions containing more than two GTs were incubated for 24 h at 30°C, supplemented with an additional 2.5 mM of each activated sugar donor, and then incubated for an additional 24 h. When desired, both CFPS reactions and IVGs could be flash-frozen frozen after their respective incubation steps. After incubation, Im7-6 was purified from IVG reactions using magnetic His-tag Dynabeads (Thermo Fisher Scientific). The IVG reactions were diluted in 90 µl of Buffer 1 (50 mM NaH2PO4 and 300 mM NaCl, pH 8.0) and centrifuged at 12,000xg for 10 min at 4°C. This supernatant was incubated at room temperature for 10 minon a roller with 20 µl of beads which had been equilibrated with 120 µl of Buffer 1. The beads were then washed three times with 120 µl of Buffer 1 and then eluted using 70 µl of Buffer 1 with 500 mM imidazole. The samples were dialyzed against Buffer 2 (12.5 mM NaH2PO4 and 75 mM NaCl, pH 7.5) overnight using 3.5 kDa MWCO microdialysis cassettes (Pierce). Purification of one-pot CFGpS reactions was completed similarly to IVG reactions.

### Production and purification of glycoproteins from living *E. coli*

The *E. coli* strain CLM24Δ*nanA* (genotype W3110 *ΔwecA ΔnanA ΔwaaL*::kan) was constructed to enable the intake and survival of sialic acid in the cytoplasm for the production of sialylated glycoproteins *in vivo*. CLM24ΔnanA was generated from W3110 using P1 transduction of the *wecA*::kan, *nanA*::kan, and *waaL*::kan alleles in that order, derived from the Keio collection^74^. Between successive transductions, the kanamycin marker was removed using pE-FLP as described previously^75^. As indicated, CLM24Δ*nanA* was sequentially transformed with the CMP-Sia production plasmid pCon.NeuA; a target protein plasmid pBR322.Im7-6 or pBR322.Fc-6; and a GT operon plasmid pMAF10.NGT, pMAF10.ApNGT.NmLgtB, pMAF10.CjCST-I.NmLgtB.ApNGT, or pMAF10.PdST6.NmLgtB.ApNGT by isolating individual clones with appropriate antibotics at each step. The completed strain was then used to inoculate a 5 ml overnight culture in LB media containing appropriate antibiotics which was then subcultured at OD600 = 0.08 into 5 ml of fresh LB media supplemented with 5 mM *N*-Acetylneuraminic acid (sialic acid) purchased from Carbosynth and adjusted to pH = 6.0 using NaOH and HCl. This culture was then grown at 37°C with shaking at 250 r.p.m. GT operon expression was induced by supplementing the culture with 0.2% arabinose at OD600 = 0.4 and then target protein expression was induced at OD600 = 1.0 with 1 mM IPTG. After IPTG induction, the culture was grown overnight at 28°C and 250 r.p.m. The cells were pelleted by centrifugation at 4°C for 10 min at 4,000xg, frozen on liquid nitrogen, and stored at −80°C. Cell pellets were thawed and resuspended in 630 μl of Buffer 1 with 5 mM imidazole and supplemented with 70 μl of 10 mg/ml lysozyme (Sigma), 1 μl (250 U) Benzonase (Millipore), and 7 µl of 100X Halt protease inhibitor (Thermo Fisher Scientific). After 15 min of thawing and resuspension, the cells were incubated for 15-60 min on ice, sonicated for 45 s at 50% amplitude, and then centrifuged at 12,000xg for 15 min. The supernatant was then incubated on a roller for 10 min at RT with 50 µl of His-tag Dynabeads which had been pre-equilibrated with 5 mM imidazole in Buffer 1. The beads were then washed three times with 1 ml of Buffer 1 containing 5 mM imidazole and then eluted with 70 μl of Buffer 1 with 500 mM imidazole by a 10 min incubation on a roller at RT. Samples were then dialyzed with 3.5 kDa MWCO microdialysis cassettes overnight against Buffer 2 before glycopeptide or glycoprotein processing and analysis for LC-MS.

### LC–MS analysis of glycoprotein modification

Modification of intact glycoprotein targets was determined by LC-MS by injection of 5 µl (or about 5 pmol) of His-tag purified, dialyzed glycoprotein into a Bruker Elute UPLC equipped with an ACQUITY UPLC Peptide BEH C4 Column, 300Å, 1.7 µm, 2.1 mm X 50 mm (186004495 Waters Corp.) with a 10 mm guard column of identical packing (186004495 Waters Corp.) coupled to an Impact-II UHR TOF Mass Spectrometer (Bruker Daltonics, Inc.). Before injection, Fc samples were reduced with 50 mM DTT. Liquid chromatography was performed using 100% H2O and 0.1% formic acid as Solvent A and 100% acetonitrile and 0.1% formic acid as Solvent B at a flow rate of 0.5 mL/min and a 50°C column temperature. An initial condition of 20% B was held for 1 min before elution of the proteins of interest during a 4 min gradient from 20% to 50% B. The column was washed and equilibrated by 0.5 min at 71.4% B, 0.1 min gradient to 100% B, 2 min wash at 100% B, 0.1 min gradient to 20% B, and then a 2.2 min hold at 20% B, giving a total 10 min run time. An MS scan range of 100-3000 m/z with a spectral rate of 2 Hz was used. External calibration was performed prior to data collection.

### LC–MS analysis of glycopeptide modification

Glycopeptides for LC-MS(/MS) analysis were prepared by digesting His-tag purified, dialyzed glycosylation targets with 0.0044 µg/µl MS Grade Trypsin (Thermo Fisher Scientific) at 37°C overnight. Before injection, H1HA10 samples were reduced by incubation with 10 mM DTT for 2 h. LC-MS(/MS) was performed by injection of 2 µl (or about 2 pmol) of digested glycopeptides into a Bruker Elute UPLC equipped with an ACQUITY UPLC Peptide BEH C18 Column, 300Å, 1.7 µm, 2.1 mm X 100 mm (186003686 Waters Corp.) with a 10 mm guard column of identical packing (186004629 Waters Corp.) coupled to an Impact-II UHR TOF Mass Spectrometer. Liquid chromatography was performed using 100% H2O and 0.1% formic acid as Solvent A and 100% acetonitrile and 0.1% formic acid as Solvent B at a flow rate of 0.5 mL/min and a 40°C column temperature. An initial condition of 0% B was held for 1 min before elution of the peptides of interest during a 4 min gradient to 50% B. The column was washed and equilibrated by a 0.1 min gradient to 100% B, a 2 min wash at 100% B, a 0.1 min gradient to 0% B, and then a 1.8 min hold at 0% B, giving a total 9 min run time. LC-MS/MS of glycopeptides was performed to confirm that GT modifications were in accordance with previously characterized specificities. Pseudo multiple reaction monitoring (MRM) MS/MS fragmentation was targeted to theoretical glycopeptide masses corresponding to detected intact protein MS peaks. All glycopeptides were fragmented using a collisional energy of 30 eV with a window of ± 2 m/z from targeted m/z values. Theoretical protein, peptide, and sugar ion masses derived from expected glycosylation structures are shown in **Supplementary Tables 3 and 5-7**. For LC-MS and LC-MS/MS of glycopeptides, a scan range of 100-3000 m/z with a spectral rate of 8 Hz was used. External calibration was performed prior to data collection.

### Exoglycosidase digestions

When possible, sugar linkages installed by various GTs and biosynthetic pathways were confirmed by exoglycosidase digestion using commercially available enzymes from New England Biolabs with well-characterized activities. As indicated in figures and figure legends, glycoproteins or glycopeptides were incubated with exoglycosidases for at least 4 h at 37°C using buffers and digestion conditions suggested by the manufacturer. The exoglycosidases and associated product numbers used in this study are: β1-4 Galactosidase S (P0745S); α1-3,6 Galactosidase (P0731S); α1-3,4 Fucosidase (P0769S); and α1-2 Fucosidase (P0724S); α1-3,4,6 Galactosidase (P0747S); β-*N*-Acetylglucosaminidase S (P0744S); α2-3 Neuraminidase S (P0743S); and α2-3,6,8 Neuraminidase (P0720S).

### LC-MS(/MS) data analysis

LC-MS(/MS) data was analyzed using Bruker Compass Data Analysis version 4.1. Glycopeptide MS and intact glycoprotein MS spectra were averaged across the full elution times of the glycosylated and aglycosylated glycoforms (as determined by extracted ion chromatograms of theoretical glycopeptide and glycoprotein charge states). MS spectra for intact glycoproteins was then analyzed by Data Analysis maximum entropy deconvolution from the full m/z scan range of 100-2,000 into a mass range of 10,000-14,000 Da for Im7-6 samples or 27,000-29,000 Da for Fc-6 samples. Representative LC-MS/MS spectra from MRM fragmentation were selected and annotated manually. Observed glycopeptide m/z and intact protein deconvoluted masses are annotated in figures and theoretical values are shown in **Supplementary Tables 3 and 5-7**. LC-MS(/MS) data was exported from Compass Data Analysis and plotted in Microsoft Excel 365.

### Statistical Information

Figure legends indicate exact sample numbers for means, standard deviations (error bars), and representative data for each experiment. No tests for statistical significance or animal subjects were used in this study.

## Supplementary Tables

**Supplementary Table 1:**
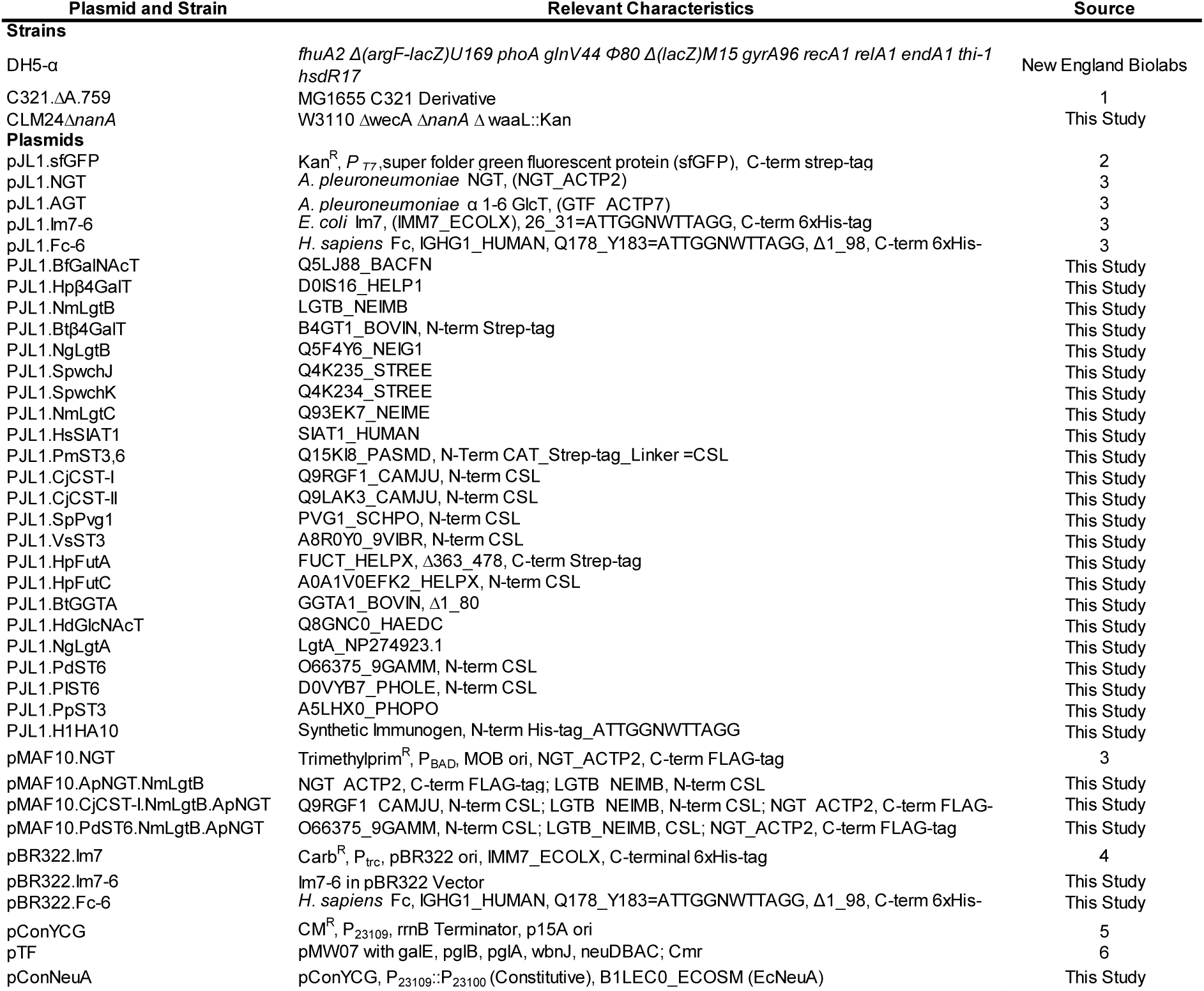
Strains and plasmids used in this study^1-6^.

**Supplementary Table 2:**
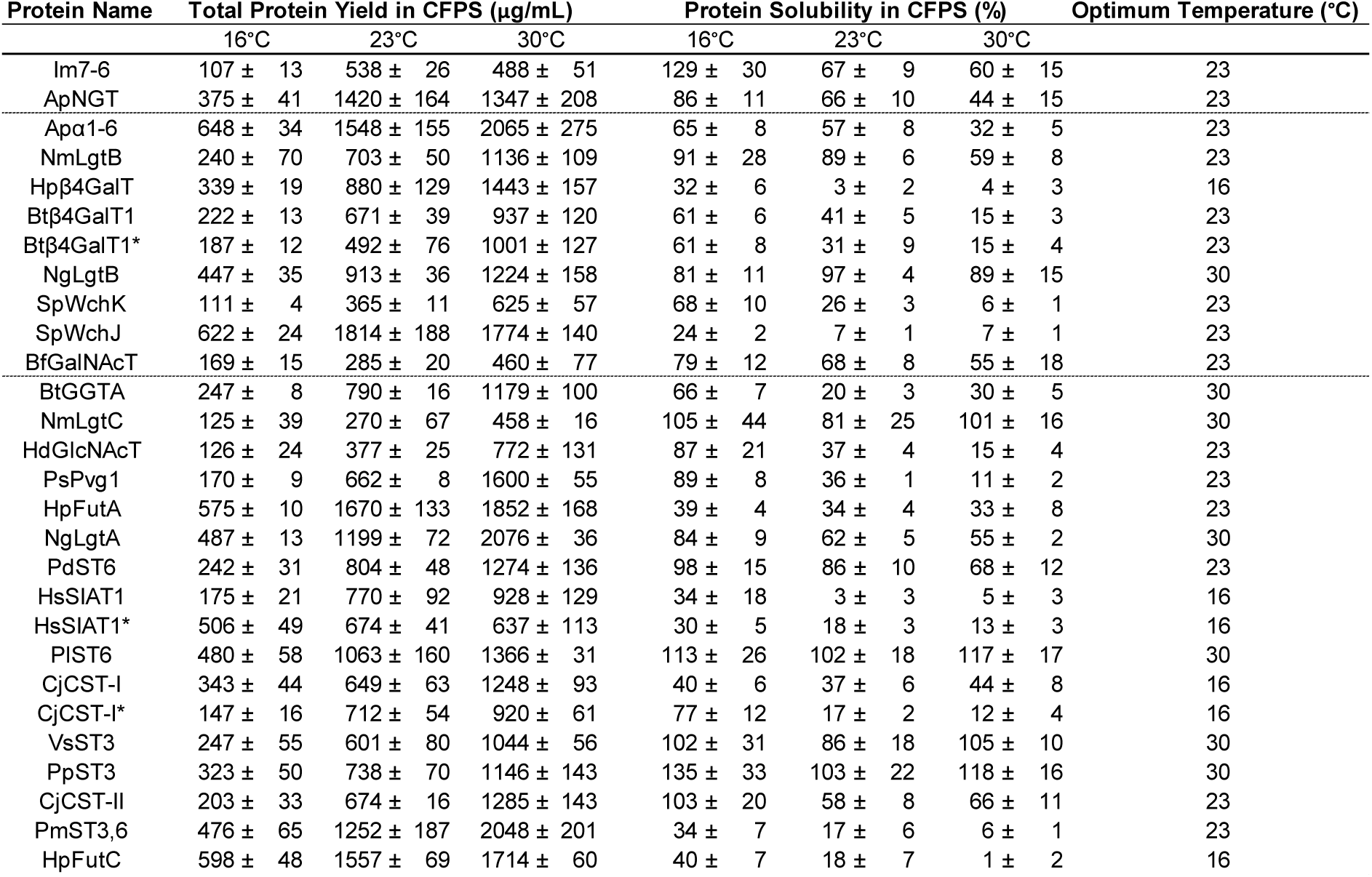
Optimization of cell-free protein synthesis of Im7 target and glycosylation enzymes. **(a)** CFPS yields of Im7-6 target and enzymes for *in vitro* glycosylation pathways tested by GlycoPRIME. CFPS yields and errors indicate mean and S.D. from n=3 CFPS reactions quantified by ^14^C-leucine incorporation. All CFPS reactions were incubated for 20 h at the indicated temperatures and conditions. Solubility was calculated from quantification of yields in fractions isolated after centrifugation at 12,000xg for 15 mins. Asterisk (*) indicates yields when CFPS was conducted under oxidizing conditions. Dotted lines indicate if enzymes were used in biosynthetic pathways with one, two, or more than three GTs. Yields under optimized conditions also shown in **Figs. 2 and 3**.

**Supplementary Table 3:**
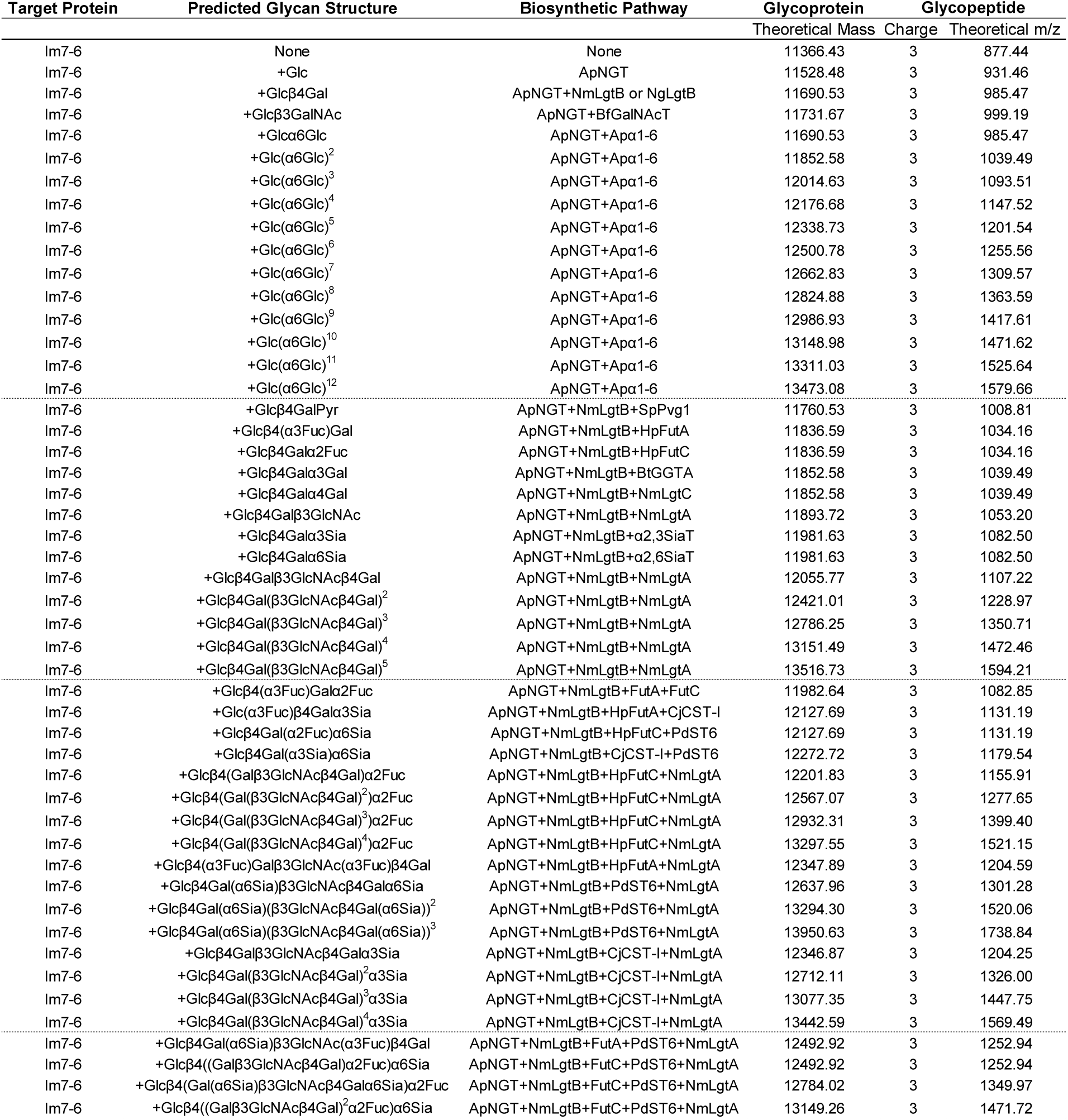
Theoretical glycoprotein and glycopeptide masses for Im7-6 glycoforms produced during GlycoPRIME biosynthetic pathway engineering. Predicted glycosylation structures are based on previously established GT activities shown in **Figs. 2 and 3 and Supplementary Table 4**. Theoretical, neutral, average masses of expected glycoprotein products and theoretical, triply charged, monoisotopic mass-to-charge ratios (m/z) of glycopeptides are shown below. Glycopeptide masses correspond to the only ApNGT glycosylation site within Im7-6 which is contained within the tryptic peptide EATTGGNWTTAGGDVLDVLLEHFVK. Experimentally observed masses are annotated in deconvoluted intact protein MS and glycopeptide MS/MS spectra.

**Supplementary Table 4:** Previously characterized activities of glycosyltransferases used this study^7-23^. GTs listed below were selected for testing in the GlycoPRIME system based on their previously established activities. Many have also been previously used for biosynthesis of glycolipids or free oligosaccharides, laying the foundation for their testing in the new context of elaborating the *N*-linked glucose installed by ApNGT in this study.

| Enzyme | Organism | Previously Characterized Activity | Reference |
| --- | --- | --- | --- |
| ApNGT | *A. pleuropneumoniae* | βGlc → Asn | 7 |
| Apα1-6 | *A. pleuropneumoniae* | (α1-6 Glc)<sup>n</sup> → Glc | 8 |
| NgLgtB | *N. gonorrhoeae* | β1-4 Gal → Glc(Nac) | 9 |
| Hpβ4GalT | *H. pylori* | β1-4 Gal → Glc(Nac) | 10 |
| Btβ4GalT1 | *B. taurus* | β1-4 Gal → Glc(Nac) | 11 |
| NmLgtB | *N. meningitidis* | β1-4 Gal → Glc(Nac) | 9 |
| SpWchK | *S. pleuropneumoniae* | β1-4 Gal → Glc | 12 |
| SpWchJ | *S. pleuropneumoniae* | WchK enhancer | 12 |
| BfGalNAcT | *B. fragilis* | β1-3 GalNAc → Glc | 13 |
| NgLgtA | *N. gonorrhoeae* | β1-3 GlcNAc → Gal | 14 |
| PsPvg1 | *S. pombe* | Pyruvate → Gal | 16 |
| HpFutA | *H. pylori* | α1-3 Fuc → Glc(Nac) | 15 |
| HpFutC | *H. pylori* | α1-2 Fuc → Gal | 17 |
| NmLgtC | *N. meningitidis* | α1-4 Gal → Gal | 17 |
| BtGGTA | *B. taurus* | α1-3 Glc → Glc | 18 |
| HsSIAT1 | *H. sapiens* | α2-6 Sia → Gal | 19 |
| PdST6 | *B. taurus* | α2-6 Sia → Gal | 20 |
| PIST6 | *P. leiognathi* | α2-6 Sia → Gal | 21 |
| PmST3,6 | *P. multocida* | α2-3,6 Sia → Gal | 21 |
| VsST3 | *V. sp. JT-FAJ-16* | α2-3 Sia → Gal | 21 |
| PpST3 | *P. phosphoreum* | α2-3 Sia → Gal | 21 |
| CjCST-I | *C. jejuni* | α2-3 Sia → Gal | 21 |
| CjCST-II | *C. jejuni* | α2-3,8 Sia → Gal | 22 |
| HdGlcNAcT | *H. ducreyi* | β1-3 GlcNAc → Gal | 23 |

**Supplementary Table 5:** Theoretical masses of sugar fragment ions detected in glycopeptide MS/MS spectra. During MS/MS fragmentation of glycopeptides, diagnostic sugar ions were detected. Theoretical mass to charge ratios of these sugar ions are shown below. All calculations of theoretical m/z assume singly charged ions. All mentions of sialic acid (Sia) in this article refer to *N*-Acetylneuraminic acid (NeuAc).

| **Sugar Structure** | **Theoretical m/z** |
| --- | --- |
| HexNAc | 204.20 |
| Sia-H<sub>2</sub>O | 274.09 |
| Sia | 292.10 |
| Hex<sup>2</sup> | 325.11 |
| HexNAc+Hex | 366.25 |
| Hex+Sia | 454.15 |
| Hex<sup>3</sup> | 487.16 |
| HexNAc+Hex+Fuc | 512.31 |
| Sia<sup>2</sup> | 583.20 |
| Hex<sup>4</sup> | 649.21 |
| HexNAc+Hex+Sia | 657.34 |
| (HexNAc+Hex)<sup>2</sup> | 731.49 |
| HexNAc+Hex+Fuc+Sia | 803.40 |
| (HexNAc+Hex)<sup>3</sup> | 1096.73 |
| (HexNAc+Hex)<sup>4</sup> | 1461.97 |

**Supplementary Table 6:** Theoretical glycopeptide masses for H1AH10 synthesized and glycosylated *in vitro*. Theoretical, doubly charged, monoisotopic mass-to-charge ratios (m/z) of the tryptic peptide containing the *N*-terminal, engineered glycosylation site within H1AH10 which was synthesized and glycosylated a one-pot *in vitro* reaction. Predicted glycosylation structures are based on previously established GT activities shown in **Figs. 2 and 3 and Supplementary Table 4**. Experimentally observed masses are annotated on deconvoluted MS and MS/MS spectra in **Fig. 4 and Supplementary Fig. 14**.

| Target Protein | Glycopeptide Sequence | Predicted Glycan Structure | Biosynthetic Pathway | Glycopeptide peptide |  |
| --- | --- | --- | --- | --- | --- |
|  |  |  |  | Charge | Theoretical m/z |
| H1HA10 | ATTGGNWTTAGGK | None | None | 2 | 611.29 |
| H1HA10 | ATTGGNWTTAGGK | +Glc | ApNGT | 2 | 692.32 |
| H1HA10 | ATTGGNWTTAGGK | +Glcβ4Gal | ApNGT + NmLgtB | 2 | 773.34 |
| H1HA10 | ATTGGNWTTAGGK | +Glcβ4Galα3Gal | ApNGT + NmLgtB + BtGGTA | 2 | 854.37 |

**Supplementary Table 7:**
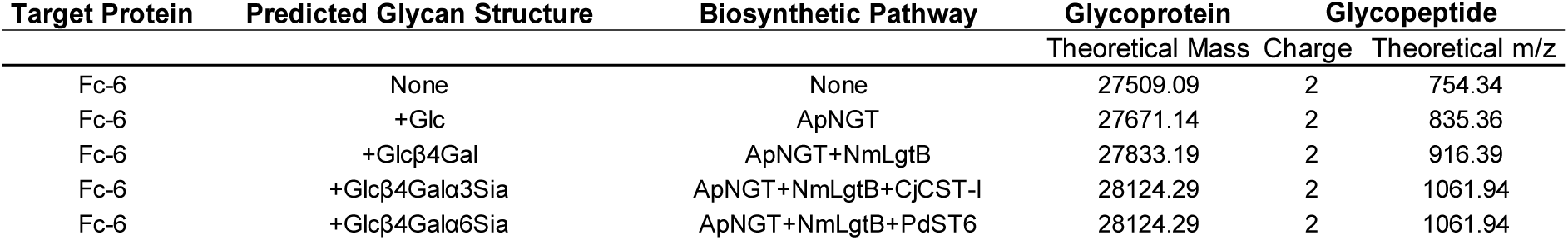
Theoretical glycoprotein and glycopeptide masses for Fc-6 synthesized and glycosylated in the *E. coli* cytoplasm. Predicted glycosylation structures are based on previously established GT activities shown in **Figs. 2 and 3 and Supplementary Table 4**. Theoretical, neutral, average masses of expected glycoprotein products and theoretical, triply charged, monoisotopic mass-to-charge ratios (m/z) of glycopeptides are shown below. Glycopeptide masses correspond to the only ApNGT glycosylation site within Fc-6 which is contained within the tryptic peptide EEATTGGNWTTAGGR. Experimentally observed masses are annotated on deconvoluted MS and MS/MS spectra in **Fig. 4 and Supplementary Fig. 15**.

| Target Protein | Predicted Glycan Structure | Biosynthetic Pathway | Glycoprotein |  | Glycopeptide |
| --- | --- | --- | --- | --- | --- |
| | | | Theoretical Mass | Charge | Theoretical $m/z$ |
| Fc-6 | None | None | 27509.09 | 2 | 754.34 |
| Fc-6 | +Glc | ApNGT | 27671.14 | 2 | 835.36 |
| Fc-6 | +Glcβ4Gal | ApNGT+NmLgtB | 27833.19 | 2 | 916.39 |
| Fc-6 | +Glcβ4Galα3Sia | ApNGT+NmLgtB+CjCST-I | 28124.29 | 2 | 1061.94 |
| Fc-6 | +Glcβ4Galα6Sia | ApNGT+NmLgtB+PdST6 | 28124.29 | 2 | 1061.94 |

## Supplementary Figures

**Supplementary Figure 1:**
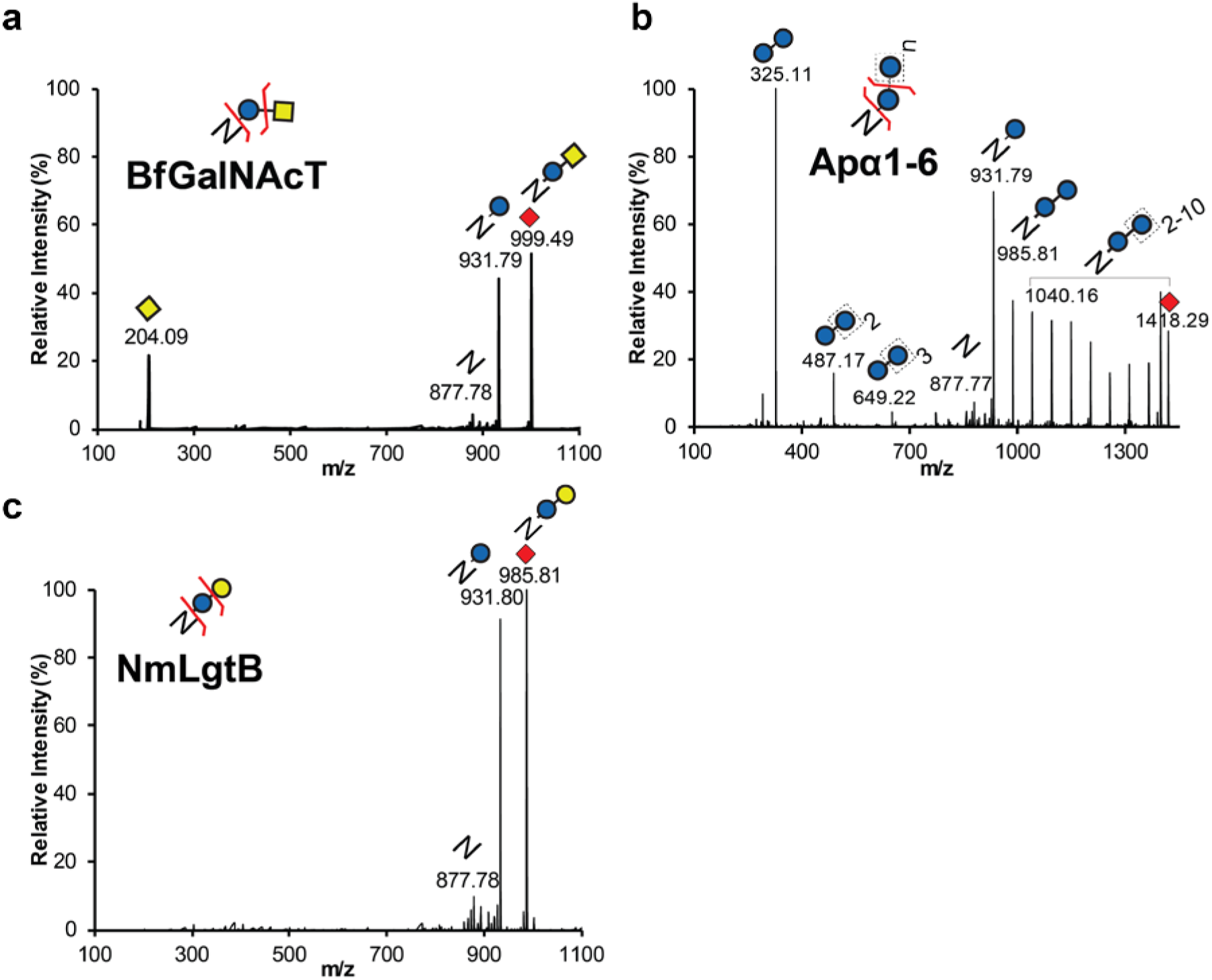
Glycopeptide MS/MS spectra of GlycoPRIME reaction products from two enzyme biosynthetic pathways elaborating *N*-linked glucose. Products from IVG reactions containing two enzyme pathways modifying Im7-6 shown in **Fig. 2** were purified, trypsinized, and analyzed by pseudo Multiple Reaction Monitoring (MRM) MS/MS fragmentation at theoretical glycopeptide masses (red diamonds) corresponding to detected protein MS peaks using a collisional energy of 30 eV (see **Methods**). Spectra representative of many MS/MS acquisitions from n=1 IVG reaction. Theoretical protein, peptide, and sugar ion masses derived from expected glycosylation structures are shown in **Supplementary Tables 3 and 5**. All indicated sugar ions are singly charged and glycopeptide fragmentation products are triply charged ions consistent with modification of Im7-6 tryptic peptide EATTGGNWTTAGGDVLDVLLEHFVK with indicated sugar structures. **(a)** MS/MS spectra of 999.49 ± 2 m/z corresponding to *N*-linked Glcβ1-3GalNAc installed by BfGalNAcT. **(b)** MS/MS spectra of 1418.29 ± 2 m/z corresponding to *N*-linked dextran polymer installed by Apα1-6. **(c)** MS/MS spectra of 985.81 ± 2 m/z corresponding with *N*-linked lactose installed by NmLgtB. All IVG reactions contained Im7-6, ApNGT, and appropriate sugar donors according to established enzyme activities (**Supplementary Table 4**).

**Supplementary Figure 2:**
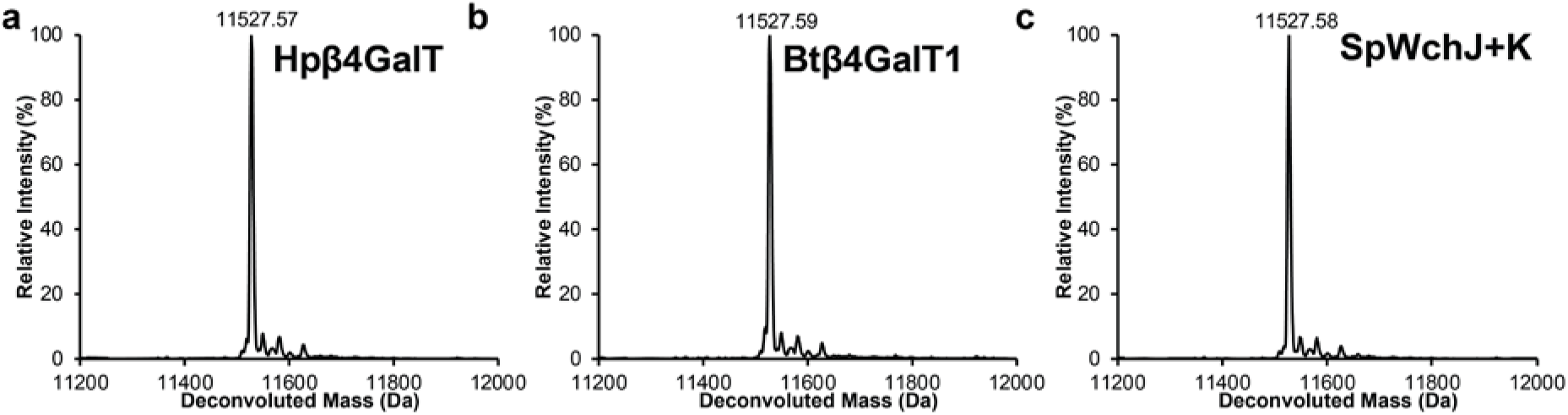
Deconvoluted intact protein MS spectra of IVG reaction products showing no modification of *N*-linked glucose installed by ApNGT. Products of IVG reactions containing 10 µM Im7-6, 0.4 µM ApNGT, 2.5 mM of appropriate sugar donors, and one elaborating GT were purified and analyzed by intact protein MS (see **Methods**). **(a)** Deconvoluted intact protein MS spectra of IVG containing 1.3 µM of Hpβ4GalT. **(b)** Deconvoluted intact protein MS spectra of IVG containing 1.4 µM of Btβ4GalT1 supplemented with 10 µM α-lactalbumin and performed under oxidizing conditions (see **Methods**). **(c)** Deconvoluted intact protein MS spectra of IVG containing 1.5 µM of SpWchJ and 1.0 µM of SpWchK. No peaks were detected that indicated the modification of Im7-6 with *N*-linked glucose installed by ApNGT (theoretical mass values shown in **Supplementary Table 3**). Spectra from m/z 100-2000 were deconvoluted into 11,000-14,000 Da using Compass Data Analysis maximum entropy method. Deconvoluted spectra shown here are representative of n=2 IVG reactions.

**Supplementary Figure 3:**
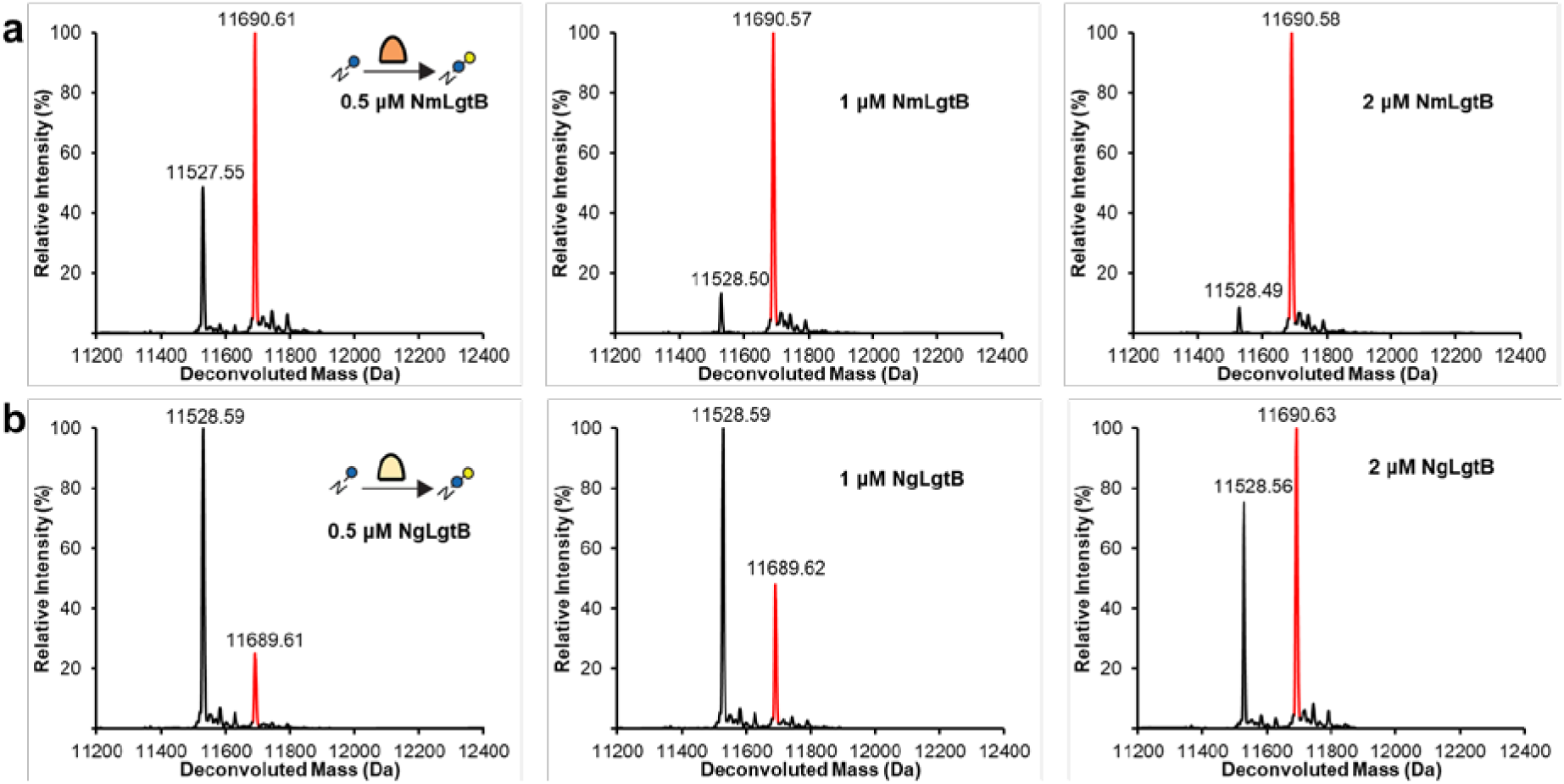
Optimization of LgtB homolog and concentration. Products of IVG reactions containing 10 µM Im7-6, 0.4 µM ApNGT, 2.5 mM of appropriate sugar donors, and indicated concentrations of NmLgtB or NgLgtB were purified and analyzed by intact protein MS (see **Methods**). **(a)** Deconvoluted intact protein MS spectra from IVG reactions containing indicated concentrations of NmLgtB. **(b)** Deconvoluted intact protein MS spectra from IVG reactions containing indicated concentrations of NgLgtB. Results representative of n=2 IVG reactions conducted for 24 h at 30°C indicate that NmLgtB produced in CFPS has greater specific activity and that nearly homogeneous *N*-linked lactose can be obtained with 2 µM NmLgtB. Theoretical mass values shown in **Supplementary Table 3**. All spectra were acquired from full elution peak areas of all detected glycosylated and aglycosylated Im7-6 species and were deconvoluted from m/z 100-2000 into 11,000-14,000 Da using Compass Data Analysis maximum entropy method.

**Supplementary Figure 4:**
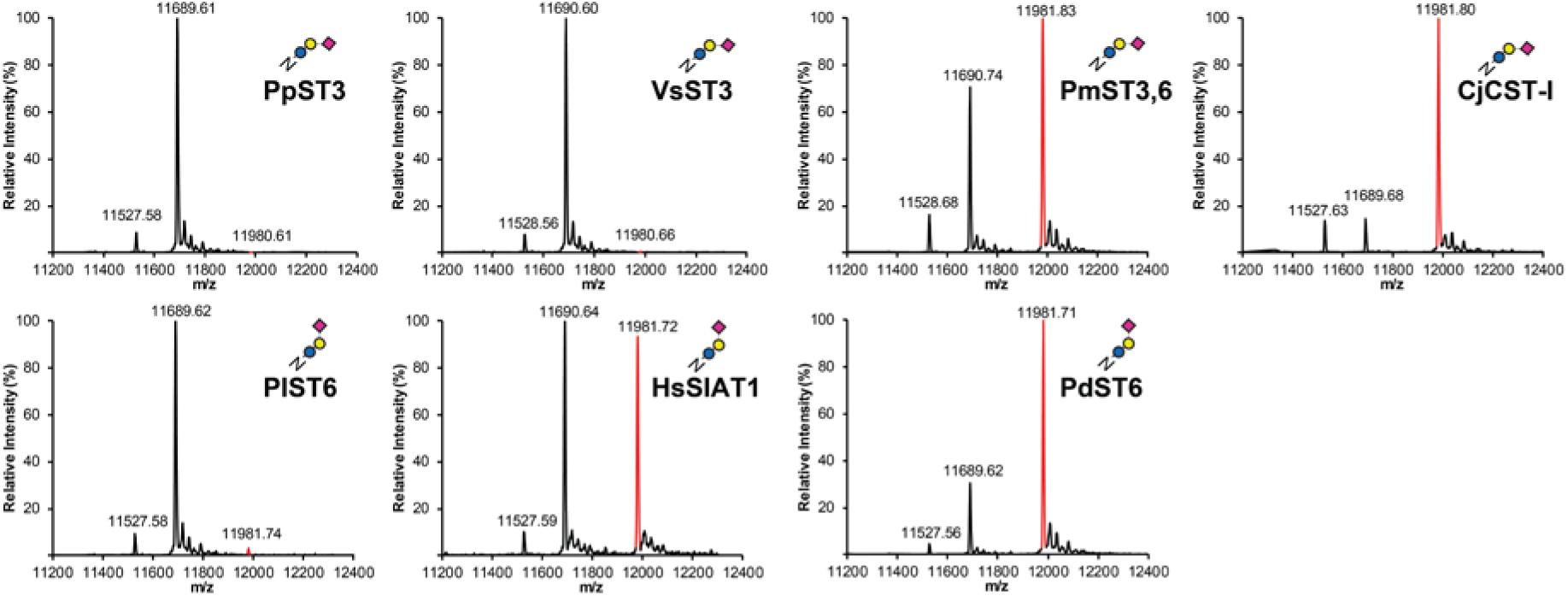
Optimization of sialyltranferase homologs. Deconvoluted intact protein MS spectra representative of n=2 IVG reactions containing 0.4 µM ApNGT, 2 µM NmLgtB, each sialyltranferase shown in **Fig. 3**, and 2.5 mM each of UDP-Glc, UDP-Gal, and CMP-Sia. Lysates enriched with sialyltransferases by CFPS were added with equal volumes to each IVG reaction such that each 32 µl-IVG reaction contained a total of 25 µl of CFPS lysates. These reactions contained 12.9 µM PpST3; 9.8 µM VsST3; 1.8 µM PmST3,6; 1.3 µM CjCST-I; 5.6 µM PlST6; 0.7 µM of HsSIAT1; and 4.9 µM PdST6, based on CFPS yields shown in **Supplementary Table 2**. CjCST-I and HsSIAT1 were synthesized in CFPS with oxidizing conditions because they were found to be more active when produced in this way (**Supplementary Fig. 7**). Under the conditions above, the reaction containing PdST6 provided the most efficient conversion to 6’-siallylactose and the reaction containing CjCST-I provided the most efficient conversion to 3’-siallylactose (exoglycosidase digestions to confirm linkages are shown in **Supplementary Fig. 8**). Although only traces amounts appear in PpST6 and VsST3, MS/MS detection and identification shows that these enzymes are functional (**Supplementary Fig. 5**). All spectra were acquired from full elution peak areas of all detected glycosylated and aglycosylated Im7-6 species and were deconvoluted from m/z 100-2000 into 11,000-14,000 Da using Compass Data Analysis maximum entropy method.

**Supplementary Figure 5:**
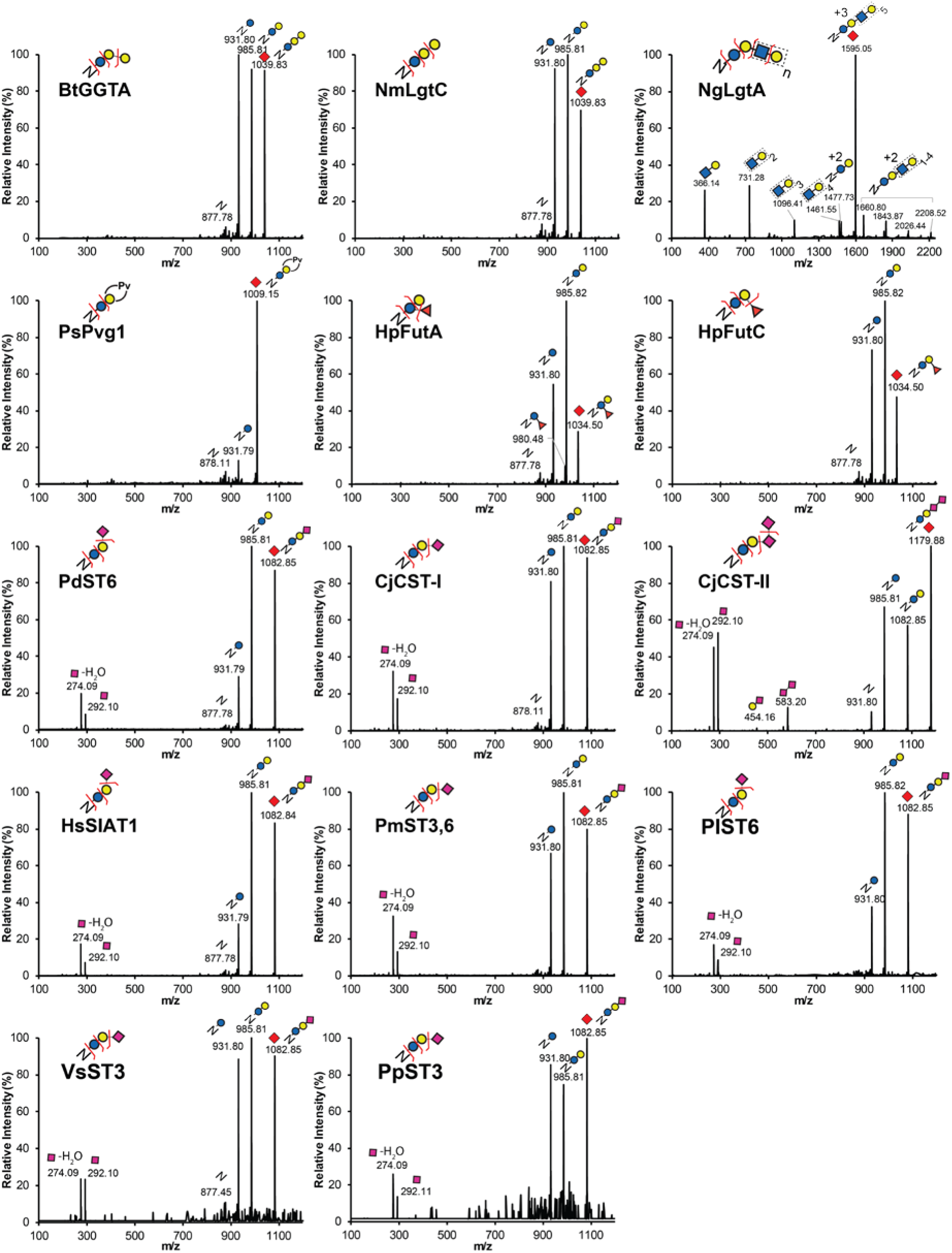
Glycopeptide MS/MS spectra of GlycoPRIME reaction products from three enzyme biosynthetic pathways elaborating *N*-linked lactose. Products from IVG reactions containing three enzyme pathways modifying Im7-6 shown in **Fig. 3** were purified, trypsinized, and analyzed by pseudo MRM MS/MS fragmentation at theoretical glycopeptide masses (indicated by red diamonds) corresponding to detected protein MS peaks in **Fig. 3 and Supplementary Fig. 4**. All glycopeptides were fragmented using a collisional energy of 30 eV with a window of ± 2 m/z from targeted m/z values (see **Methods**). Spectra are representative of many MS/MS acquisitions from n=1 IVG reaction. Theoretical protein, peptide, and sugar ion masses derived from expected glycosylation structures are shown in **Supplementary Tables 3 and 5**. All indicated sugar ions are singly charged and glycopeptide fragmentation products are triply charged ions consistent with modification of Im7-6 tryptic peptide EATTGGNWTTAGGDVLDVLLEHFVK with indicated sugar structures. Predicted sugar linkages based on previously established GT activities (**Supplementary Table 4**) and exoglycosidase sequencing (**Supplementary Figs. 8 and 9**). All IVG reactions contained Im7-6, ApNGT, NmLgtB, indicated GTs, and appropriate sugar donors according to established GT activities.

**Supplementary Figure 6:**
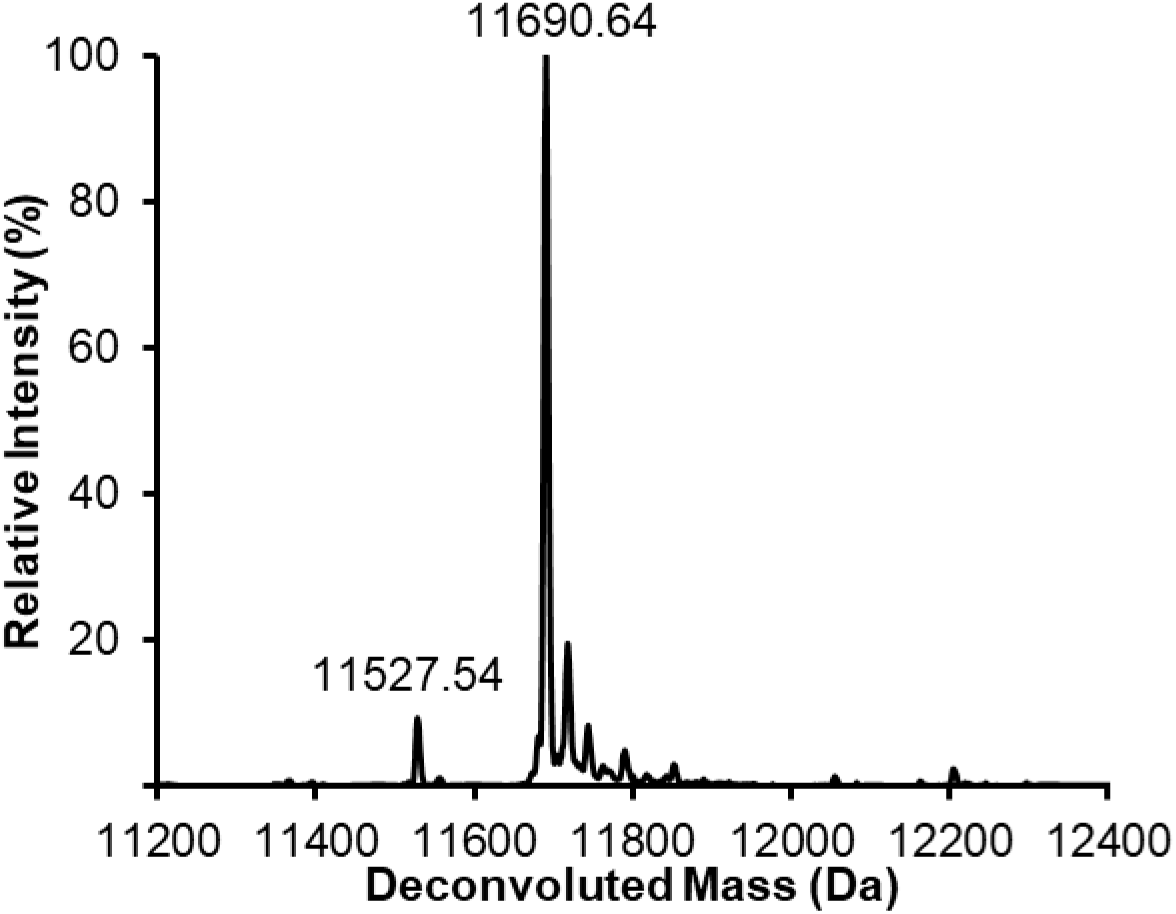
HdGlcNAcT does not modify the *N*-linked lactose substrate installed by ApNGT and NmLgtB. Deconvoluted intact protein MS spectra of IVG reaction product containing 10 µM Im7-6, 0.4 µM ApNGT, 2 µM NmLgtB, 1.5 µM HdGlcNAcT, and 2.5 mM of UDP-Glc, UDP-Gal, and UDP-GlcNAc. No peaks were detected that indicated the modification of Im7-6 with *N*-linked lactose installed by ApNGT and NmLgtB (see **Supplementary Table 3** for theoretical mass values). Deconvoluted spectra representative of n=2 IVG reactions.

**Supplementary Figure 7:**
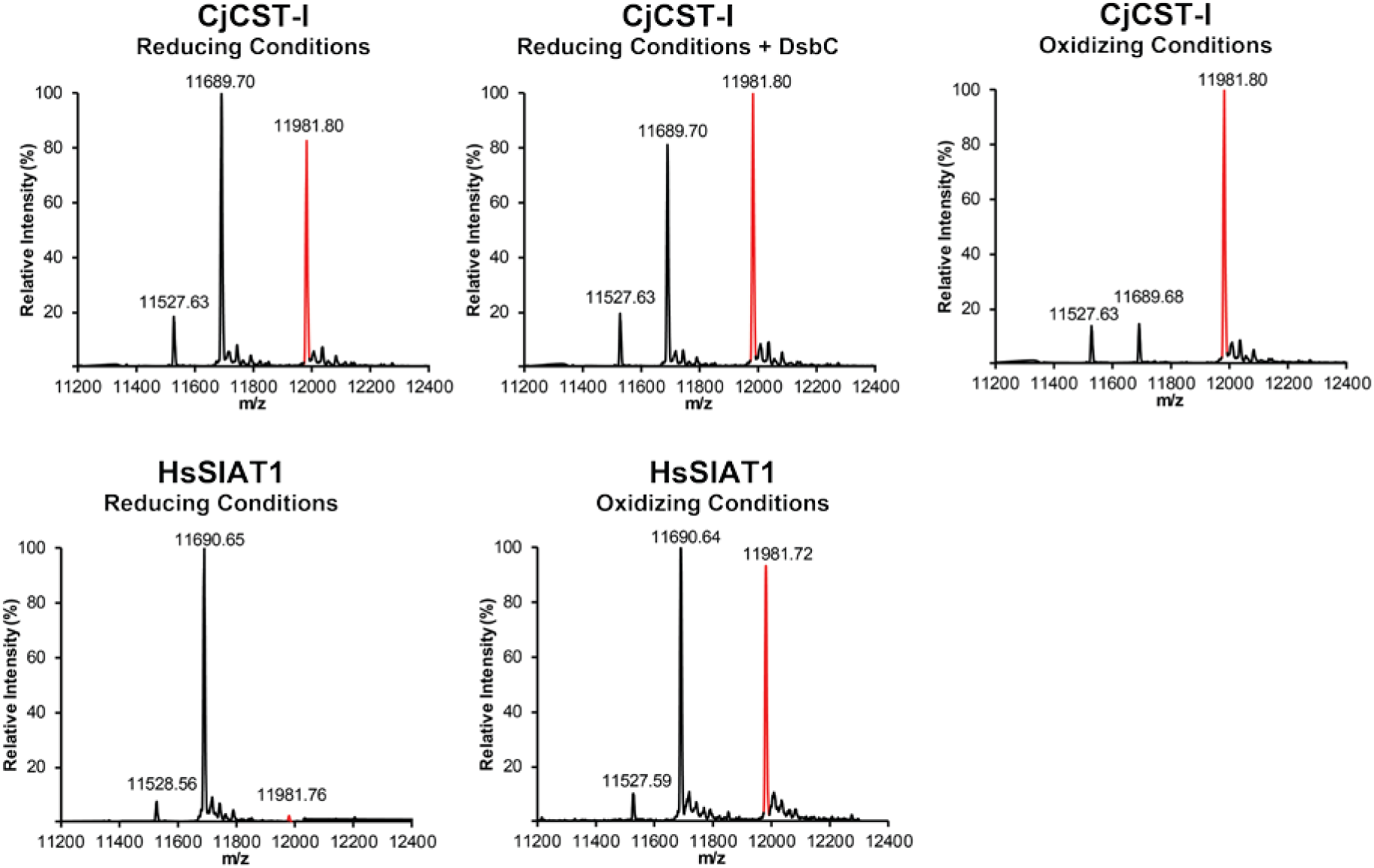
CjCST-I and HsSIAT1 exhibit greater activity when produced in oxidizing conditions. Deconvoluted intact protein MS spectra representative of of n=2 IVG reaction products containing 10 µM Im7-6, 0.4 µM ApNGT, 2 µM NmLgtB, 2.5 mM of UDP-Glc, UDP-Gal, and CMP-Sia as well as CjCST-I or HsSIAT1 made in CFPS conducted under oxidizing conditions, reducing conditions with supplemented the *E. coli* disulfide bond isomerase (DsbC), or standard reducing conditions (see **Methods**). CFPS conditions are known to create a protein synthesis environment conducive to disulfide bond formation as previously described^24^. Lysates enriched with sialyltranferases by CFPS were added in equal volumes. Therefore, reducing reaction conditions contained 1.9 µM of CjCST-I or 3.8 µM of HsSIAT1 while oxidizing reaction conditions reactions contained 1.3 µM of CjCST-I and 0.7 µM of HsSIAT1 (detailed CFPS yield information shown in **Supplementary Fig. 2**). Aside from CFPS synthesis conditions for the CjCST-I and HsSIAT1, IVG reactions were performed identically without ensuring an oxidizing environment for glycosylation. Im7-6, ApNGT, and NmLgtB were produced with standard CFPS reaction conditions. Relative glycosylation efficiencies indicate that the oxidizing CFPS environment of CFPS allows for greater enzyme activities per unit of CFPS reaction volume and per µM of enzyme. This observation makes sense for HsSIAT1 which is normally active in the oxidizing environment of the human golgi and is known to contain disulfide bonds. Interestingly, an oxidizing synthesis environment also seems to benefit the activity of CjCST-I which does not contain disulfide bonds. However, the increased activity of CjCST-I cannot be explained by the general chaperone activity of DsbC.

**Supplementary Figure 8:**
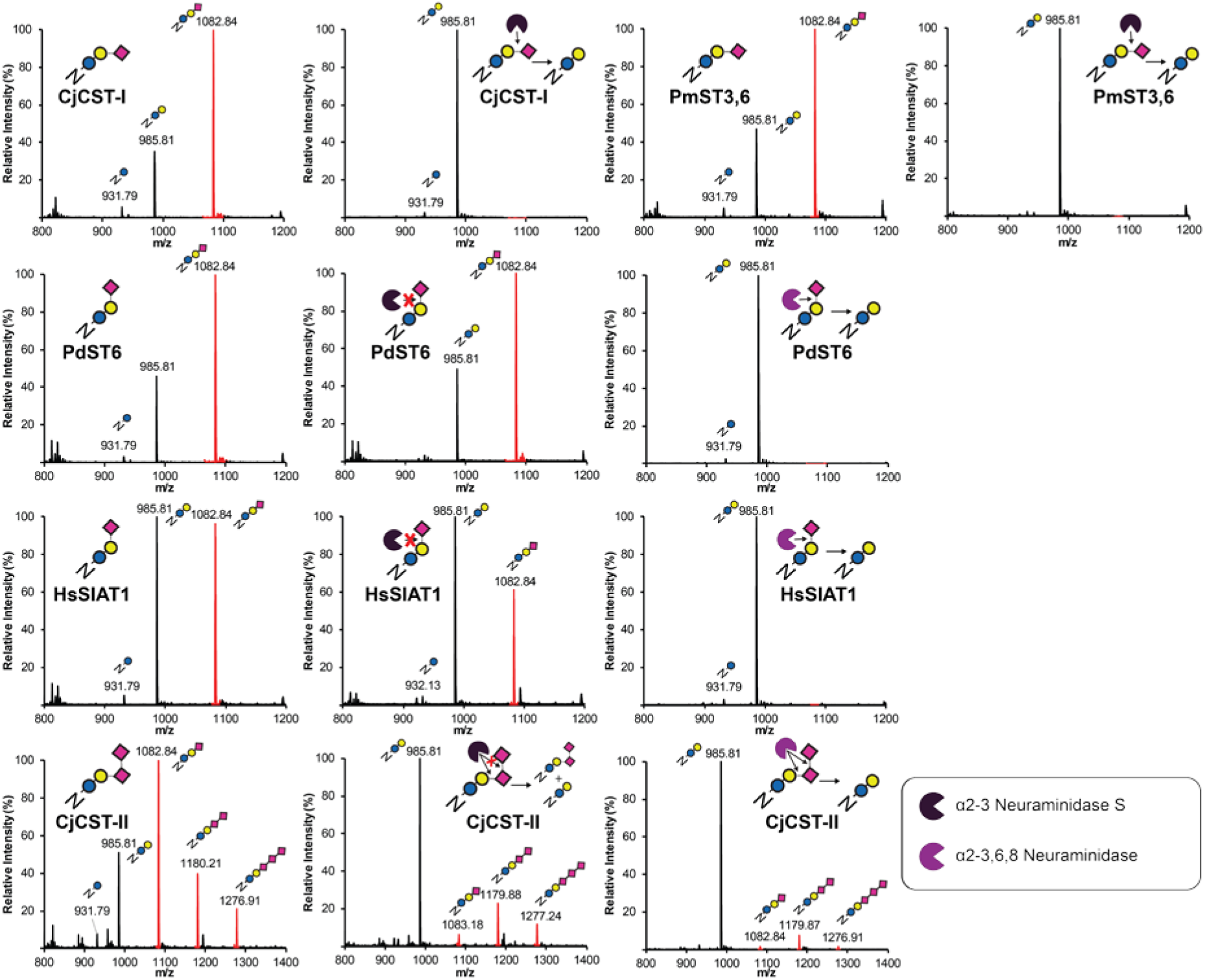
Exoglycosidase sequencing of Im7-6 modified by GlycoPRIME biosynthetic pathways containing sialic acids. Completed IVG reactions from the GlycoPRIME workflow where purified using Ni-NTA magnetic beads, incubated at 37°C for at least 4 h with and without indicated commercially available exoglycosidases, trypsinized overnight, and then analyzed by glycopeptide LC-MS. The α2-3 Neuraminidase S was able to remove the sialic acids installed by CjCST-I; PmST3,6; and the first sialic acid installed by CjCST-II, indicating that these enzymes were installed sialic acids with α2-3 linkages. Sialic acids installed by PdST6, HsSIAT1, as well as the second and third sialic acids installed by CjCST-II were resistant to digestion by α2-3 Neuraminidase S but were susceptible to cleavage by an α2-3,6,8 Neuraminidase which is consistent with the established α2-6 activity of PdST6 and HsSIAT1 and the α2,8 linkages installed by CjCST-II in subsequent sialic acid additions. See **Methods** section for exoglycosidase details. All spectra were acquired from full elution peak areas of all detected glycosylated and aglycosylated species of the Im7-6 tryptic peptide EATTGGNWTTAGGDVLDVLLEHFVK containing an ApNGT glycosylation acceptor sequence. All indicated glycopeptide products are triply charged ions consistent with this Im7-6 tryptic peptide modified with indicated sugar structures.

**Supplementary Figure 9:**
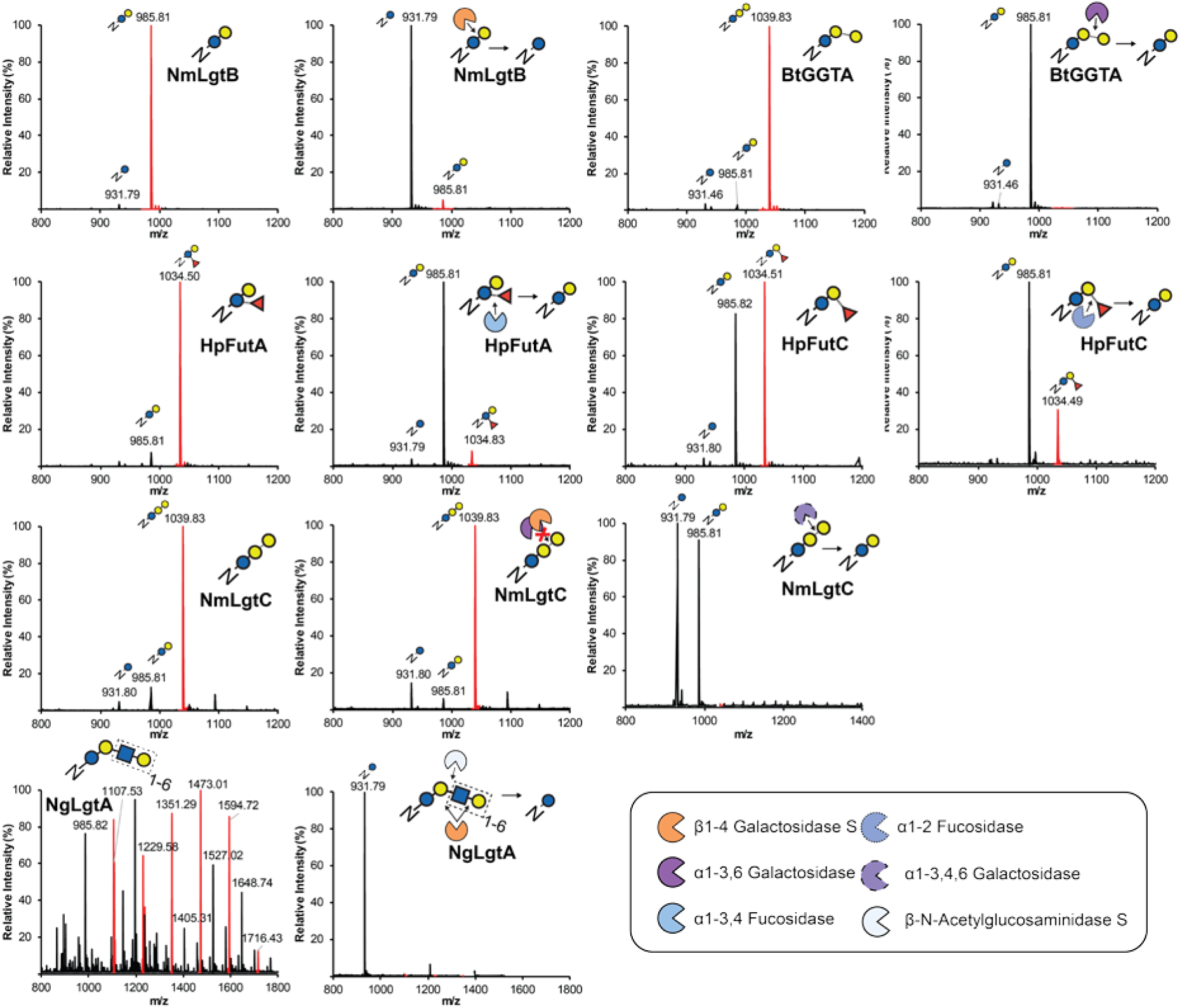
Exoglycosidase sequencing of Im7-6 modified by GlycoPRIME biosynthetic pathways not containing sialic acids. Completed IVG reactions from the GlycoPRIME workflow where purified using Ni-NTA magnetic beads, incubated at 37°C for at least 4 h with and without indicated commercially available exoglycosidases, trypsinized overnight, and then analyzed by glycopeptide LC-MS. The sugars installed by NmLgtB, BtGGTA, HpFutA, and HpFutC were susceptible to cleavage by commercially available β1-4 Galactosidase S; α1-3,6 Galactosidase; α1-3,4 Fucosidase; and α1-2 Fucosidase, respectfully. The galactose installed by NmLgtC was resistant to cleavage by β1-4 Galactosidase S and α1-3,6 Galactosidase, but susceptible to cleavage by α1-3,4,6 Galactosidase. The LacNAc polymer installed by alternating activities by NmLgtB and NgLgtA was susceptible to cleavage by a mixture of β1-4 Galactosidase S and the β-*N*-Acetylglucosaminidase S. All spectra were acquired from full elution peak areas of all detected glycosylated and aglycosylated species of the Im7-6 tryptic peptide EATTGGNWTTAGGDVLDVLLEHFVK containing an ApNGT glycosylation acceptor sequence. All indicated glycopeptide products are triply charged ions consistent with this Im7-6 tryptic peptide modified with indicated sugar structures. Cleavage observations are consistent with previously established GT activities (**Figs. 2-3 and Supplementary Table 4**). See **Methods** section for exoglycosidase details.

**Supplementary Figure 10:**
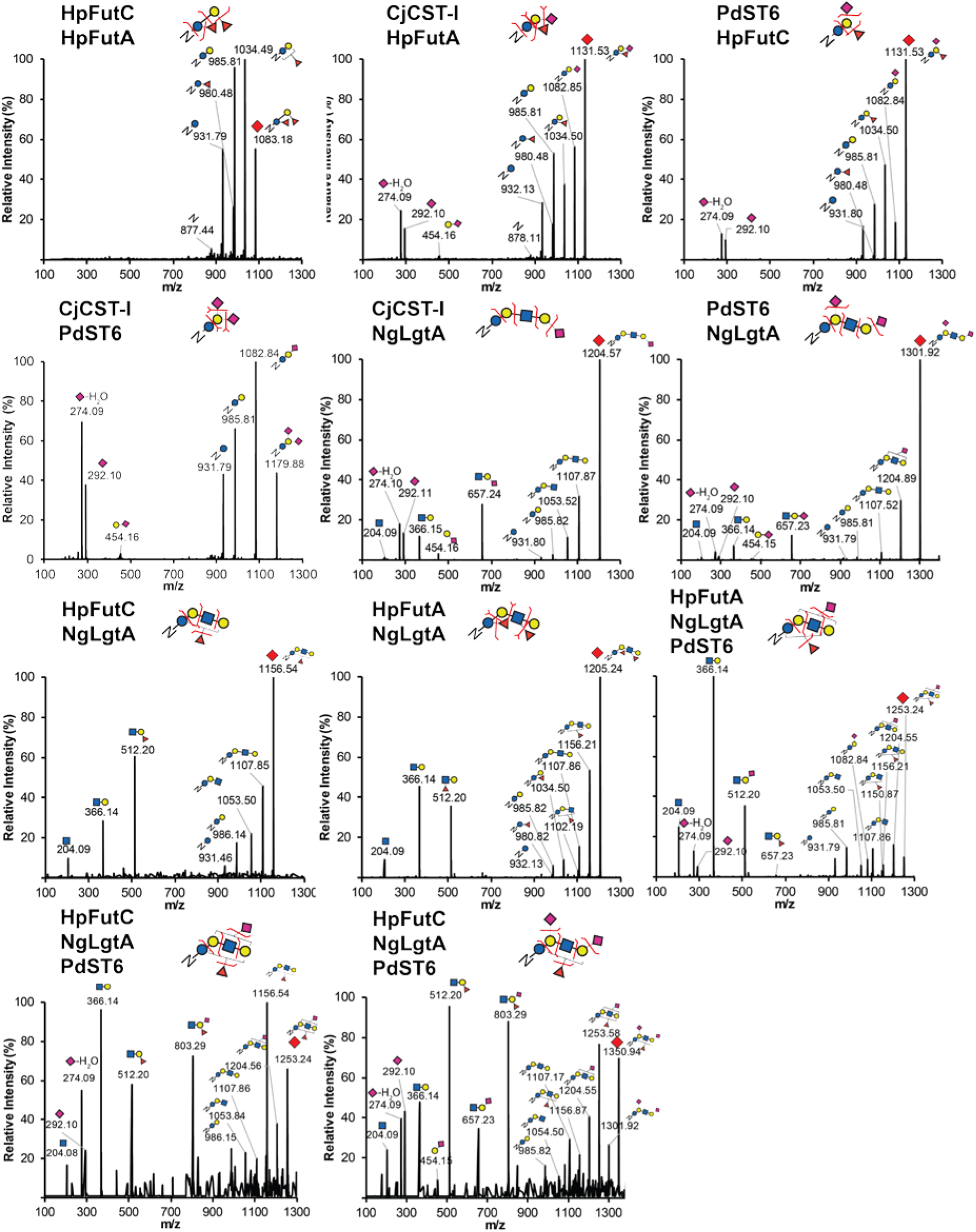
Glycopeptide MS/MS spectra of GlycoPRIME reaction products from four and five enzyme biosynthetic pathways elaborating *N*-linked lactose. Products from IVG reactions containing four and five enzyme pathways modifying Im7-6 shown in **Fig. 3d and Supplementary Fig. 12** were purified, trypsinized, and analyzed by pseudo MRM MS/MS fragmentation at theoretical glycopeptide masses (indicated by red diamonds) corresponding to detected protein MS peaks in **Fig. 3d and Supplementary Fig. 12**. All glycopeptides were fragmented using a collisional energy of 30 eV with a window of ± 2 m/z from targeted m/z values (see **Methods**). Spectra representative of many MS/MS acquisitions from n=1 IVG reaction. Theoretical protein, peptide, and sugar ion masses derived from expected glycosylation structures are shown in **Supplementary Tables 3 and 5**. All indicated sugar ions are singly charged and glycopeptide fragmentation products are triply charged ions consistent with modification of Im7-6 tryptic peptide EATTGGNWTTAGGDVLDVLLEHFVK with indicated sugar structures. Predicted sugar linkages based on previously established GT activities (**Supplementary Table 4**). Although products from five-enzyme biosynthetic pathway product could not be unambiguous defined, sugar and glycopeptide fragments do suggest modification with both fucose and sialic acids. All IVG reactions contained Im7-6, ApNGT, NmLgtB, indicated enzymes, and appropriate sugar donors according to established GT activities.

**Supplementary Figure 11:**
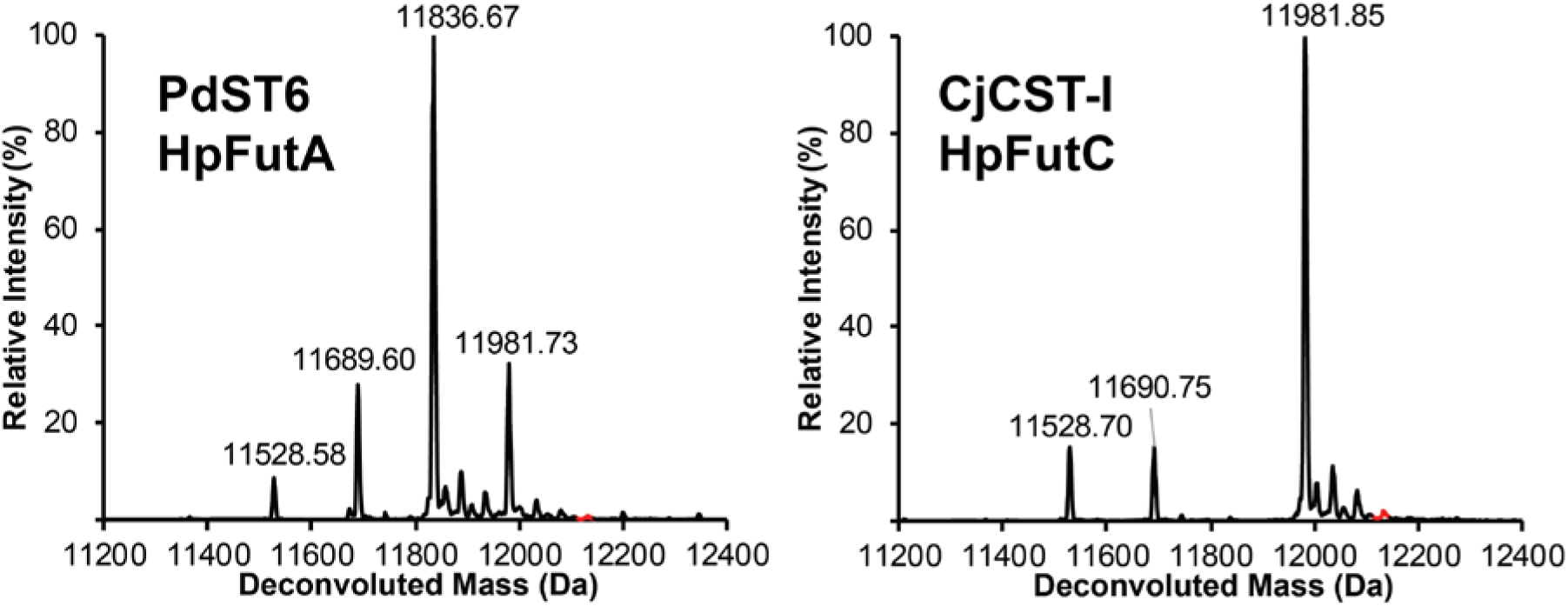
Deconvoluted intact protein MS spectra of IVG reaction products showing no production fucosylated and sialylated species. Products of IVG reactions containing 10 µM Im7-6, 0.4 µM ApNGT, 2 µM NmLgtB, indicated enzymes, and 2.5 mM of appropriate sugar donors (UDP-Glc, UDP-Gal, CMP-Sia, and GDP-Fuc) were purified and analyzed by intact protein MS. Reactions contained 2.4 µM HpFutA and 2.4 µM PdST6 or 1.3 µM HpFutC and 0.65 µM CjCST-I as indicated. Deconvoluted spectra representative of n=2 IVGs. No peaks were detected that indicated the presence of Im7-6 modified with both a sialic acid and a fucose (the region of the spectra annotated in red line shows expected range of sialylated and fucosylated species) (see **Supplementary Table 4** for theoretical mass values).

**Supplementary Figure 12:**
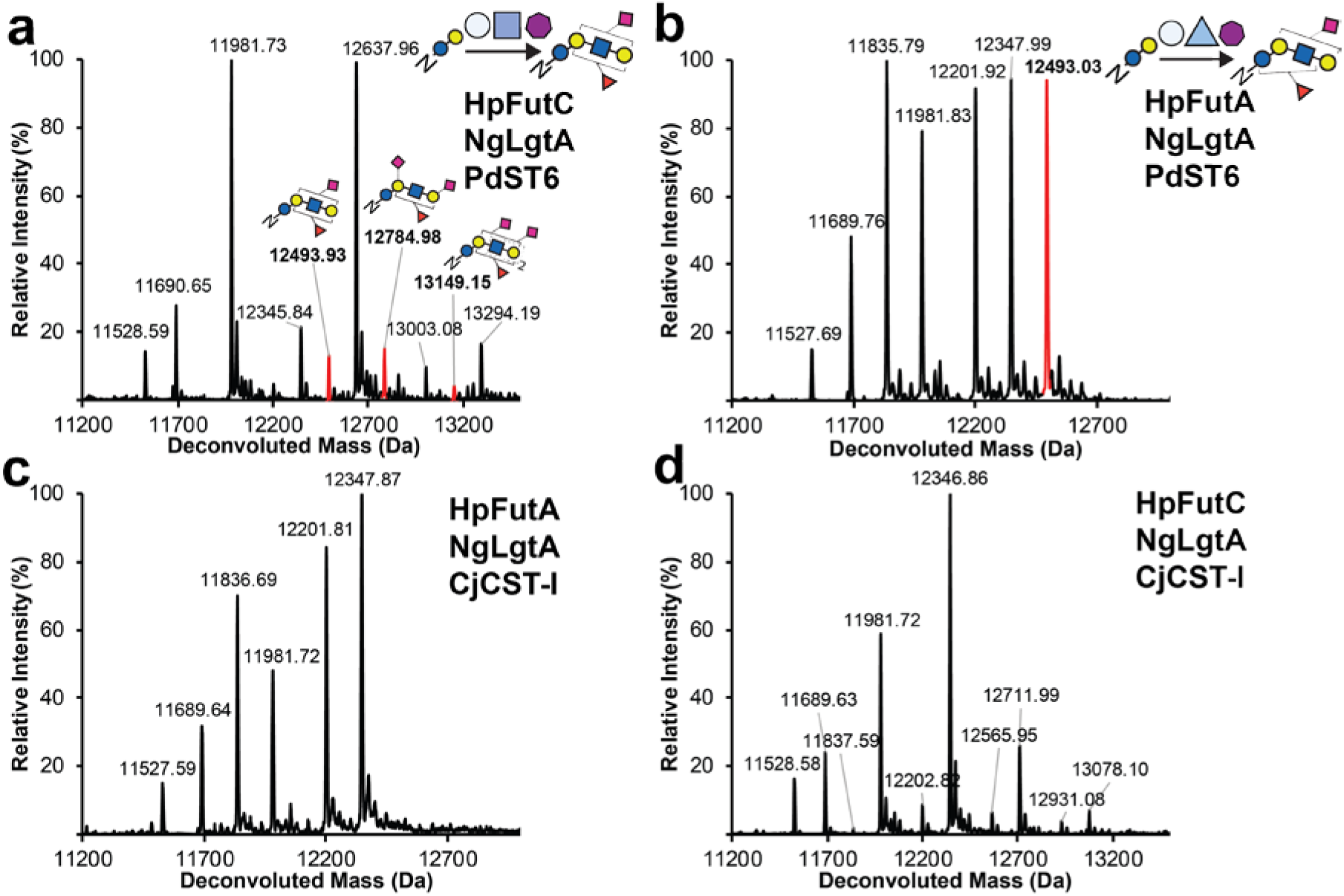
GlycoPRIME screening of biosynthetic pathways containing five enzymes. Products of IVG reactions containing 10 µM Im7-6, 0.4 µM ApNGT, 2 µM NmLgtB, indicated GTs, and 2.5 mM of appropriate sugar donors (UDP-Glc, UDP-Gal, CMP-Sia, and GDP-Fuc) were purified from and analyzed by intact protein MS. Deconvoluted spectra representative of n=2 IVGs. **(a)** Deconvoluted intact protein MS of IVG reactions containing 0.87 µM HpFutC, 3.83 µM NgLgtA, and 1.63 µM PdST6. **(b)** Deconvoluted intact protein MS of IVG reactions containing 1.63 µM HpFutA, 3.83 µM NgLgtA, and 1.63 µM PdST6 (also shown in **Fig. 3d**) **(c)** Deconvoluted intact protein MS of IVG reactions containing 1.63 µM HpFutA, 3.83 µM NgLgtA, and 0.43 µM CjCST-I. **(d)** Deconvoluted intact protein MS of IVG reactions containing 0.87 µM HpFutC, 3.83 µM NgLgtA, and 0.43 µM CjCST-I. Spectra in **a** and **b** as well as fragmentation spectra in **Supplementary Fig. 10** indicated three and one species, respectively, which contained both sialic acid and fucose. Predicted glycosylation structures based on previously established GT activities (**Supplementary Table 4**) and fragmentation spectra (**Supplementary Fig. 10**). Although structures cannot be unambiguously identified, the previously observed incompatibility of HpFutA and PdST6 as well as the presence of a 1083 m/z peak (Glcβ4Galα6Sia) and the absence of a 1034 m/z (Glc(α3Fuc)β4Gal) peak in fragmentation spectra suggests that in **b** the proximal galactose is modified with a sialic acid while the GlcNAc is modified with the fucose. No peaks in **c** or **d** were detected that indicated the presence of Im7-6 modified with both a sialic acid and a fucose (see **Supplementary Table 3** for theoretical mass values).

**Supplementary Figure 13:**
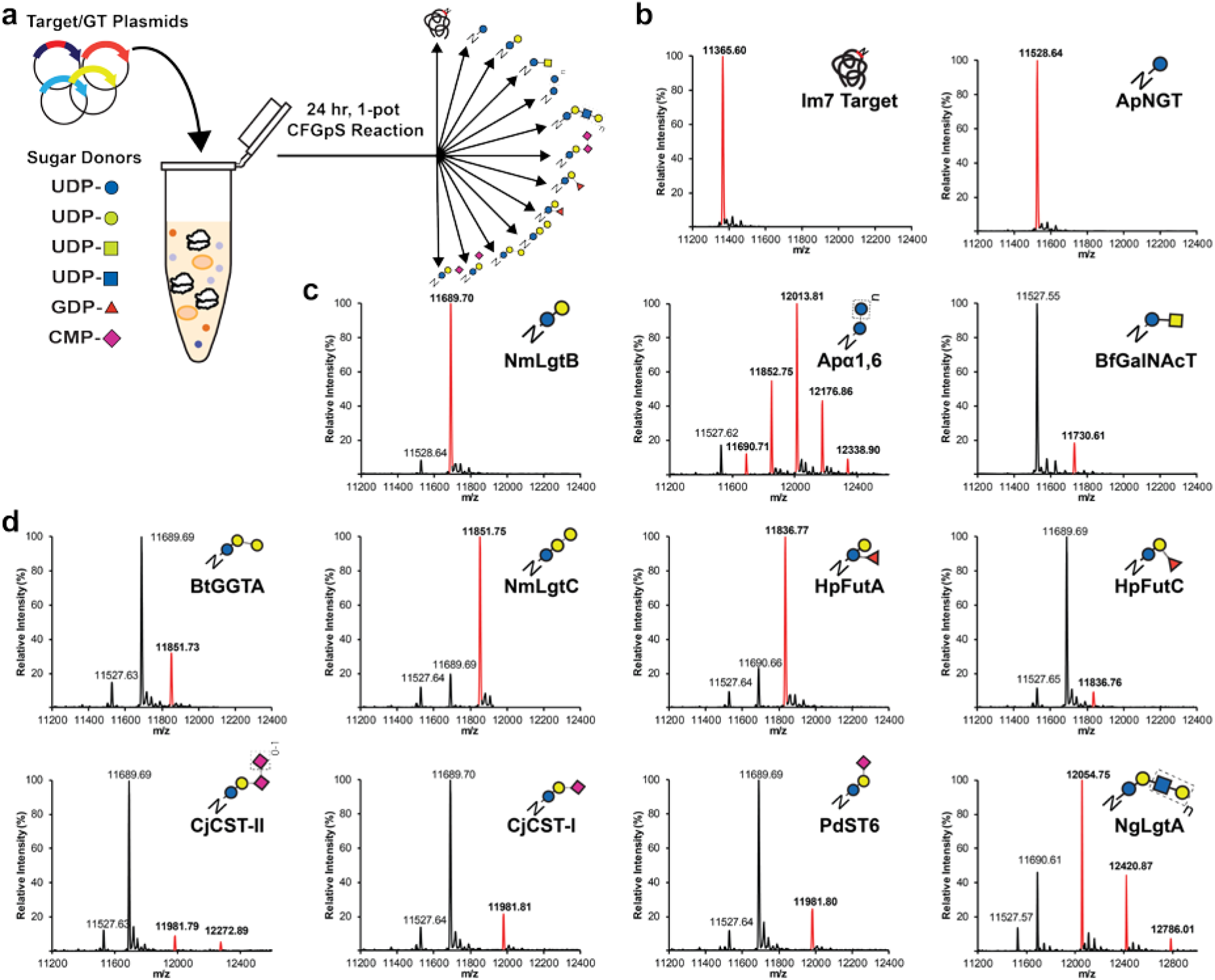
Intact protein MS spectra of Im7-6 synthesized and glycosylated by one-pot CFGpS reactions. (**a**) Plasmids encoding the Im7-6 target protein and sets of up to three GTs based on 12 successful biosynthetic pathways developed by two-pot GlycoPRIME screening were combined with appropriate sugar donors in a CFGpS reaction and incubated for 24 h at 30°C. (**b**) Deconvoluted intact protein spectra from Im7-6 synthesized and glycosylated in CFGpS reactions with and without ApNGT plasmid. (**c**) Deconvoluted intact protein spectra from Im7-6 synthesized and glycosylated in CFGpS reactions with ApNGT plasmid and indicated GT plasmids. (**d**) Deconvoluted intact protein spectra from Im7-6 synthesized and glycosylated in CFGpS reactions with ApNGT, NmLgtB, and indicated GT plasmids. All reactions contained equimolar amounts of each plasmid and a total plasmid concentration of 10 nM. All Im7-6 proteins were purified using Ni-NTA magnetic beads before intact protein analysis (see **Methods**). All reactions showed intact protein mass shifts consistent with the modification of Im7-6 with the same glycans observed in our two-pot system (**Figs. 2-3**), although at lower efficiency. MS spectra were acquired from full elution areas of all detected glycosylated and aglycosylated protein or peptide species and are representative of n=2 CFGpS reactions. Deconvoluted spectra collected from m/z 100-2000 into 11,000-14,000 Da using Compass Data Analysis maximum entropy method. See **Supplementary Fig. 3** for theoretical mass values.

**Supplementary Figure 14:**
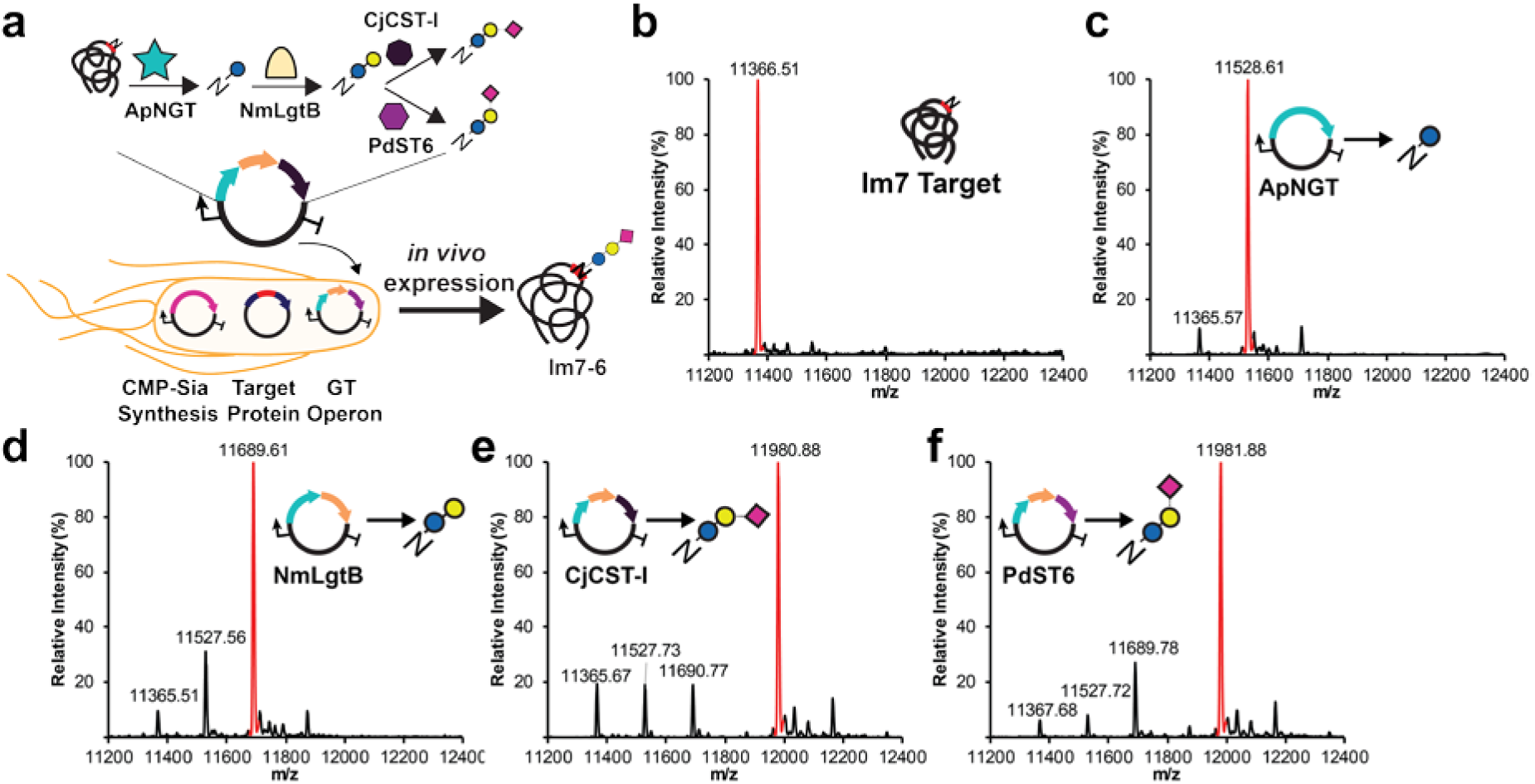
Production of sialylated Im7-6 in the *E. coli* cytoplasm. (**a**) Design of cytoplasmic glycosylation system to produce sialylated glycoproteins in *E. coli*. Three plasmids containing NmNeuA (CMP-Sia synthesis), target protein containing ApNGT glycosylation acceptor sequence, and biosynthetic pathways discovered using GlycoPRIME (GT operon). (**b-f**) Deconvoluted intact protein spectra from Im7-6 purified from CLM24*ΔnanA E. coli* strain containing CMP-Sia synthesis plasmid and Im7-6 target protein plasmid as well as no GT operon **b**; GT operon containing ApNGT **c**; GT operon containing ApNGT and LgtB **d**; GT operon containing ApNGT, NmLgtB, and CjCST-I **e**; or GT operon containing ApNGT, NmLgtB, and PdST6 **f**. The last GT in all glycosylation pathways is indicated. Mass shifts in intact protein spectra are consistent with established activities of each GT and the installation of *N*-linked Glc, lactose, 3’-sialyllactose, and 6’-sialyllactose onto Im7-6 in **b, c, d, e, and f**, respectively. All *E. coli* cultures were supplemented with 5 mM sialic acid and grown to OD600 = 0.6 at 37°C, induced with 1 mM IPTG and 0.2% arabinose, and then incubated overnight at 25°C. MS spectra were acquired from full elution areas of all detected glycosylated and aglycosylated protein species and were deconvoluted from m/z 100-2000 into 11,000-14,000 Da using Compass Data Analysis maximum entropy method. See **Supplementary Table 3** for theoretical masses. Spectra representative of n=2 bacterial cultures.

**Supplementary Figure 15:**
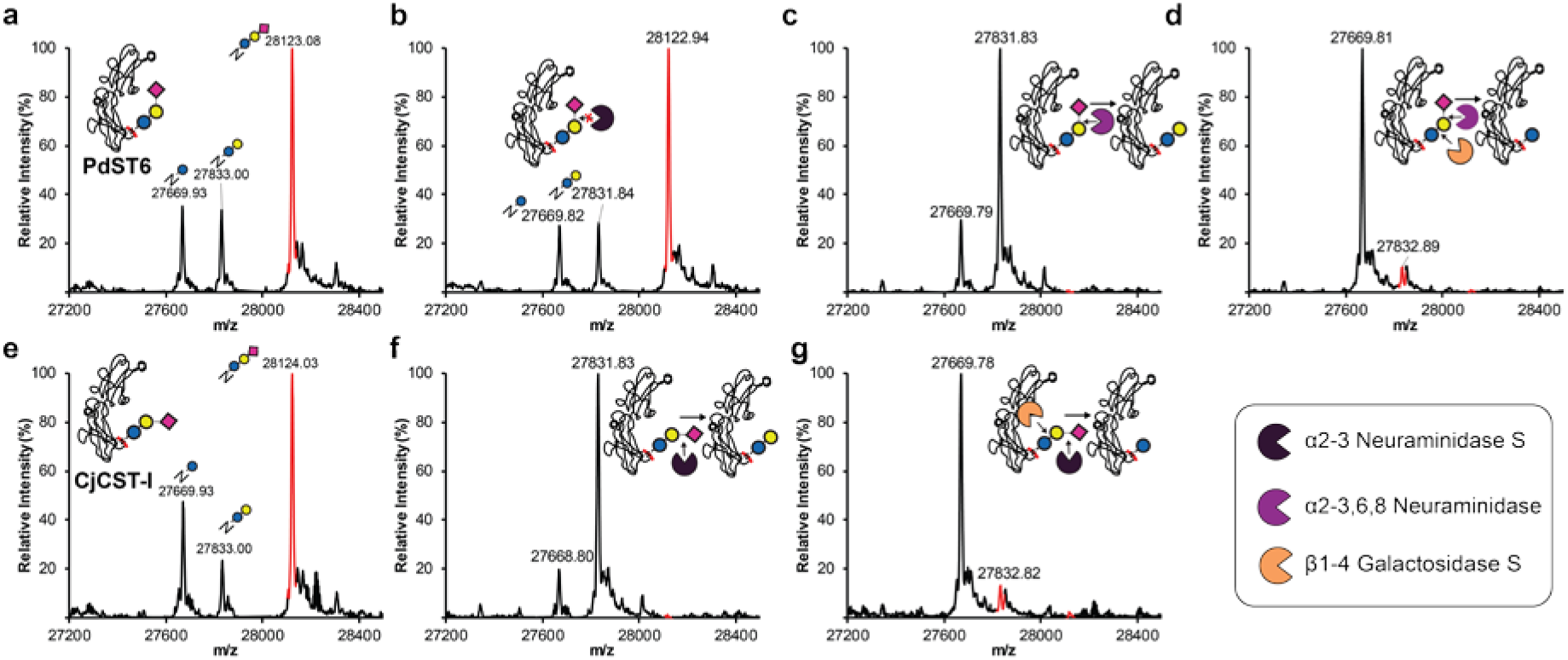
Exoglycosidase sequencing of Fc glycosylated in the *E. coli* cytoplasm. (**a**) Deconvoluted intact protein spectra from Fc-6 purified from CLM24Δ*nanA E. coli* strain containing CMP-Sia synthesis plasmid, Fc-6 target protein plasmid, and a GT operon plasmid containing ApNGT, NmLgtB, and PdST6. (**b-d**) Purified Fc-6 from **a** was incubated at 37°C for at least 4 h with commercially available α2-3 Neuraminidase S **b**, α2-3,6,8 Neuraminidase **c**, or β1-4 Galactosidase S and α2-3,6,8 Neuraminidase **d**. Resistance of terminal sialic acid to α2-3 Neuraminidase S and susceptibility to α2-3,6,8 Neuraminidase indicates an α2-6 linkage, which is consistent with previously established activity of PdST6 (**Supplementary Table 4**). (**e**) Deconvoluted intact protein spectra from Fc-6 purified from CLM24Δ*nanA E. coli* strain containing CMP-Sia synthesis plasmid, Fc-6 target protein plasmid, and a GT operon plasmid containing ApNGT, NmLgtB, and CjCST-I. (**f-g**) Purified Fc-6 from **e** was incubated at 37°C for at least 4 h with commercially available α2-3 Neuraminidase S **b**, or β1-4 Galactosidase S and α2-3 Neuraminidase S. Susceptibility of terminal sialic acid to α2-3 Neuraminidase confirms the previously established activity of CjCST-I (**Supplementary Table 4**). Removal of middle galactose with addition β1-4 Galactosidase S in **d** and **g** confirms the previously established activity of NmLgtB (**Supplementary Table 4**). **a-c** and **e-f** are also shown in **Fig. 4.** See **Methods** for exoglycosidase details and **Supplementary Table 7** for theoretical glycoprotein masses. All *E. coli* cultures were supplemented with 5 mM sialic acid and grown to OD600 = 0.6 at 37°C then induced with 1 mM IPTG and 0.2% arabinose then incubated overnight at 25°C. MS spectra were acquired from full elution areas of all detected glycosylated and aglycosylated protein species and were deconvoluted from m/z 100-2000 into 27,000-29,000 Da using Compass Data Analysis maximum entropy method.

---

**Supplementary Note 1: DNA sequences encoding engineered glycosylation targets, *in vitro* expressed glycosyltranferases, *in vivo* glycosyltranferases operons, and *in vivo* CMP-Sia production plasmid.**

## Key

TRANSLATED REGION

## Engineered glycosylation acceptor sequence

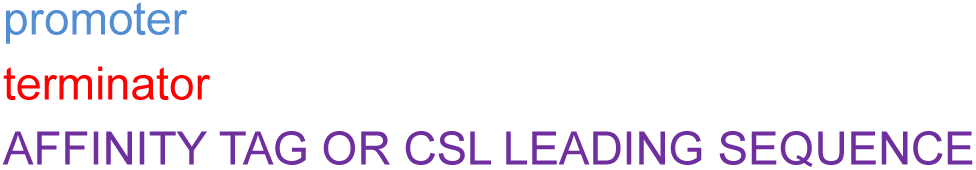

## DNA sequence for Im7-6 Variant in pJL1 plasmid context

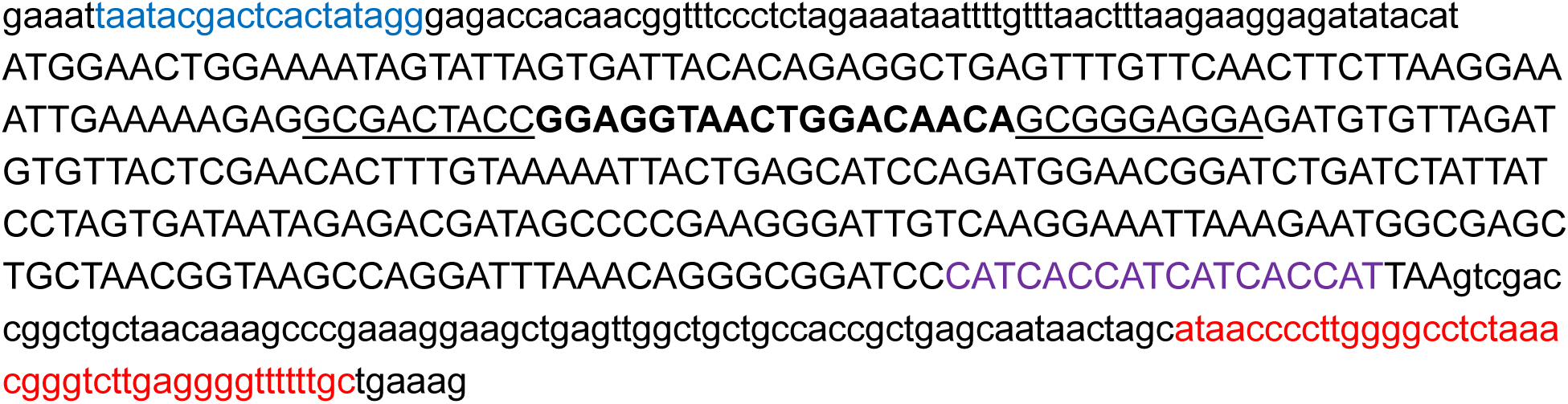

## DNA Sequence of ApNGT in pJL1 Context

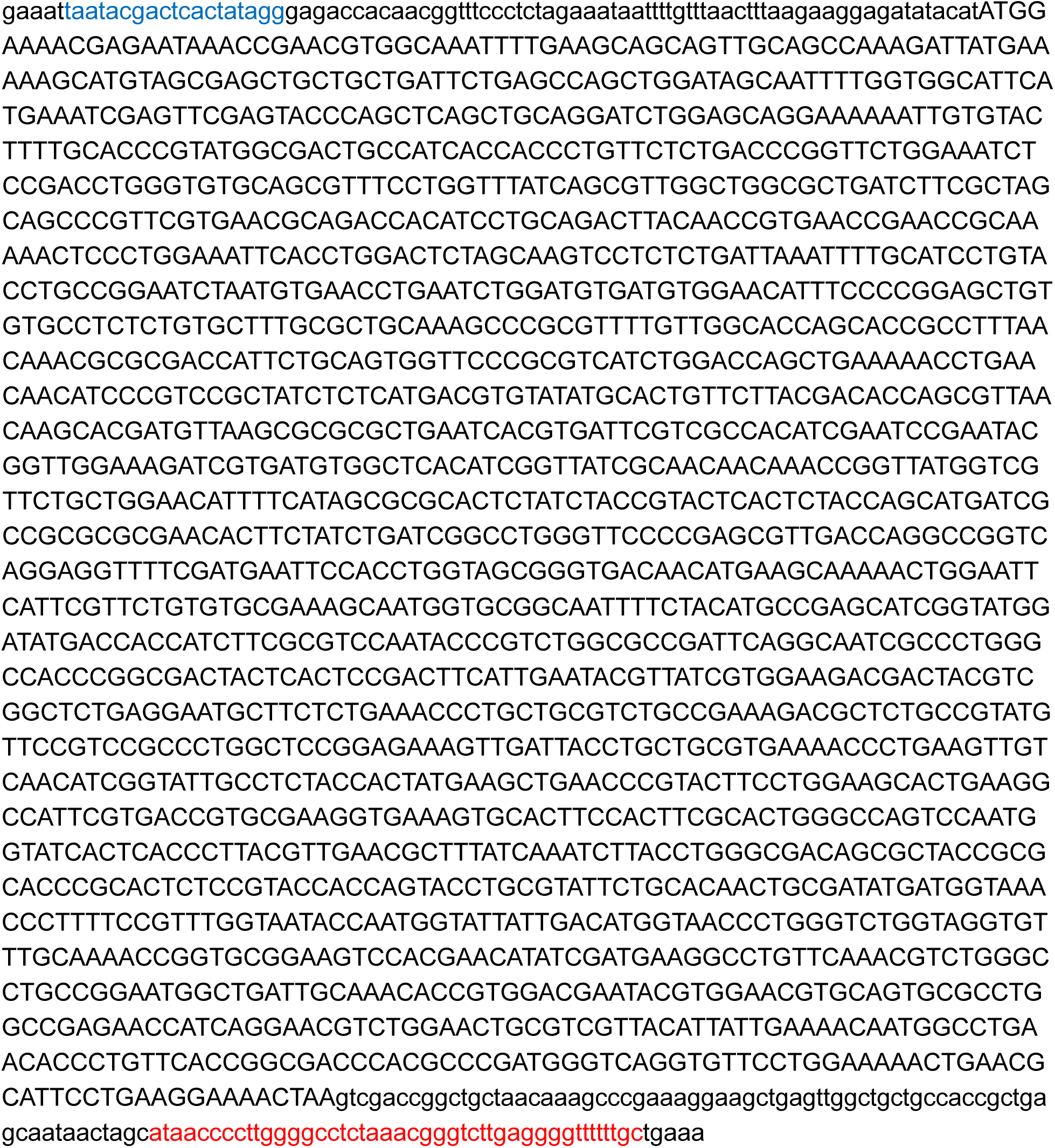

## DNA Sequence of Apα1-6 in pJL1 Context

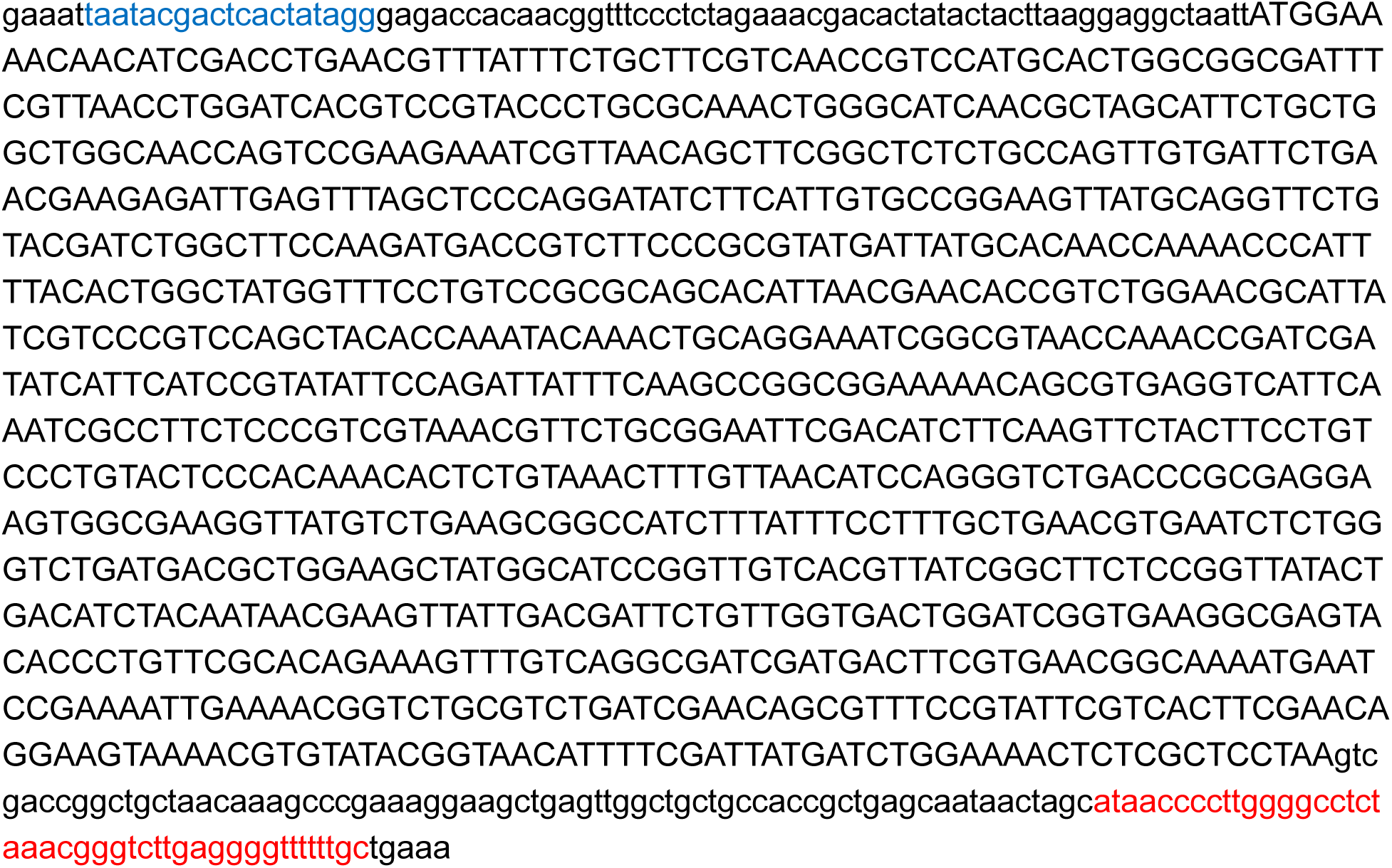

## DNA Sequence of BfGalNAcT1 in pJL1 Context

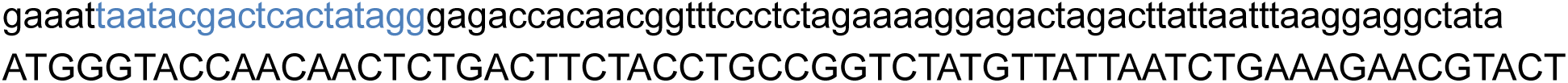

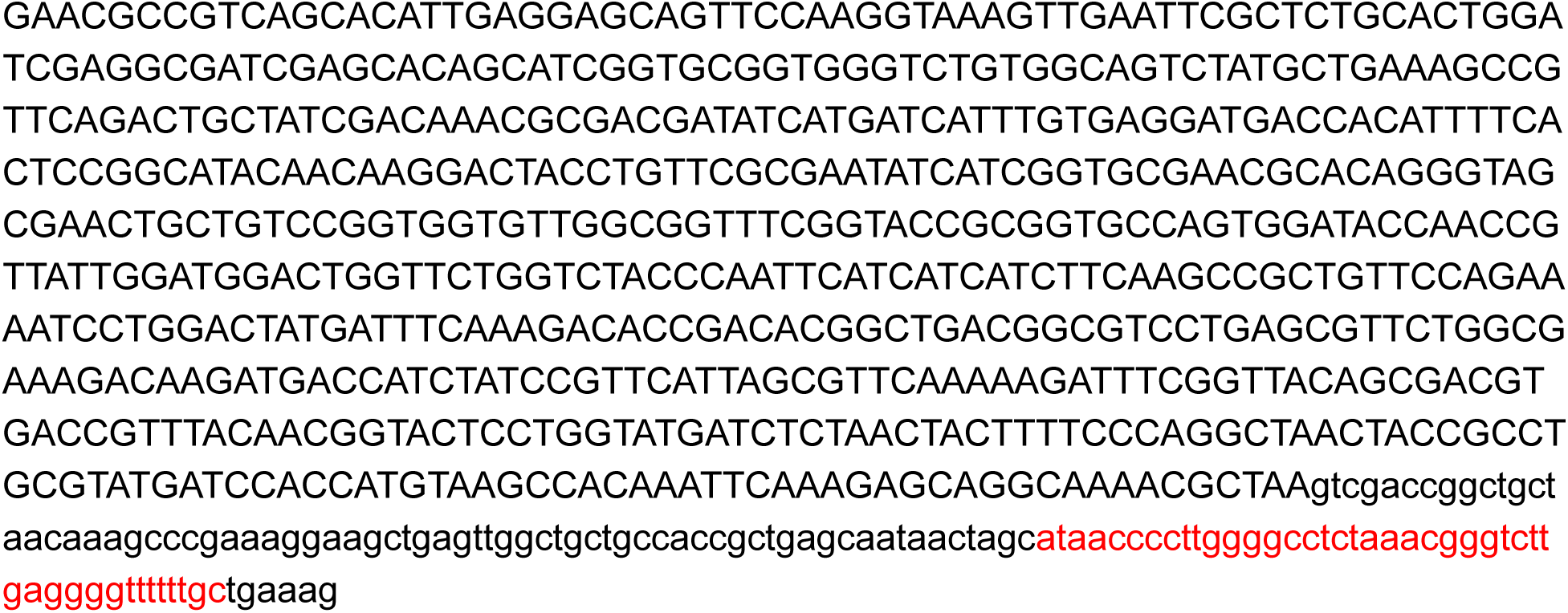

## DNA Sequence of Hpβ4GalT in pJL1 Context

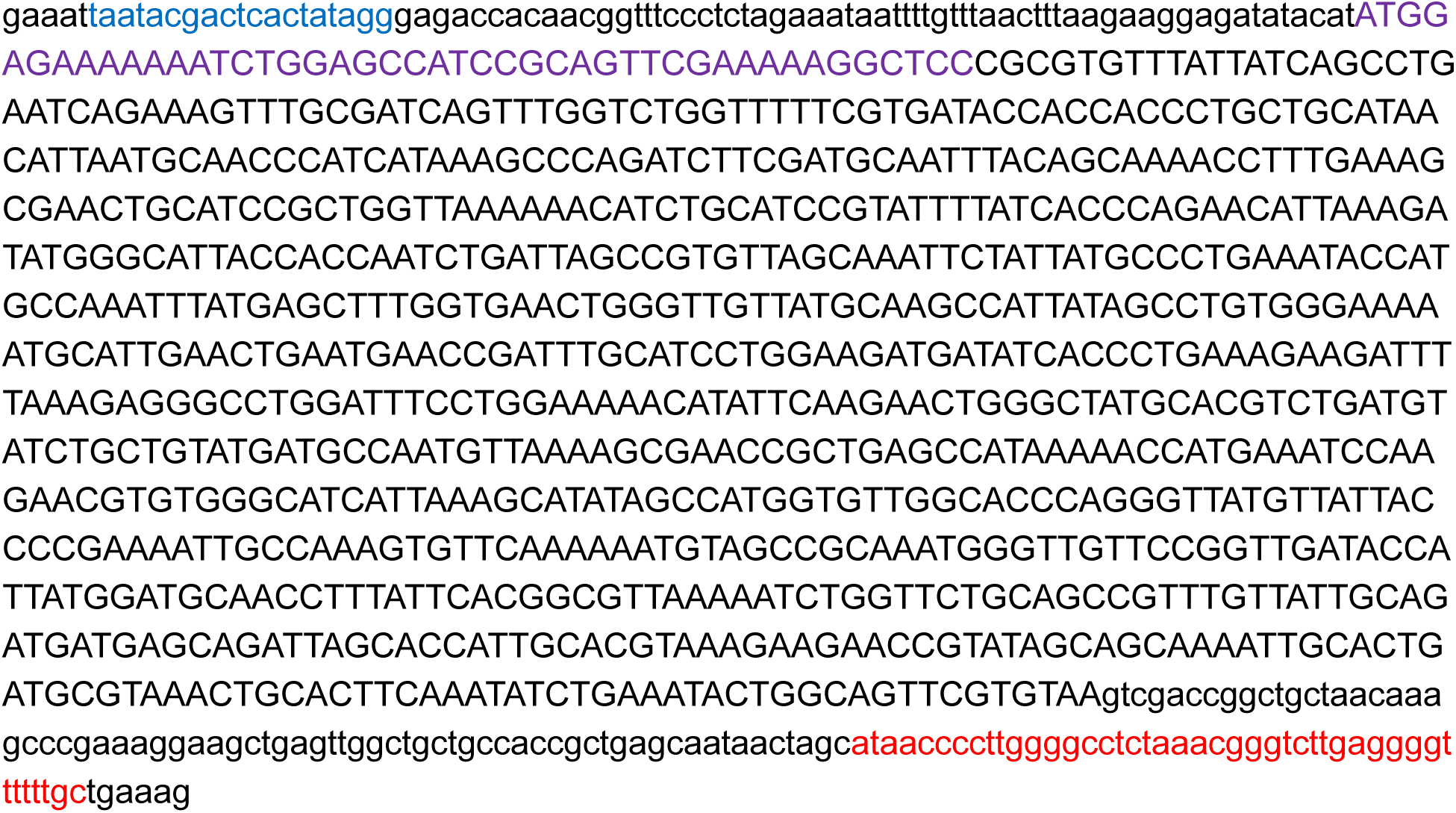

## DNA Sequence of NmLgtB in pJL1 Context

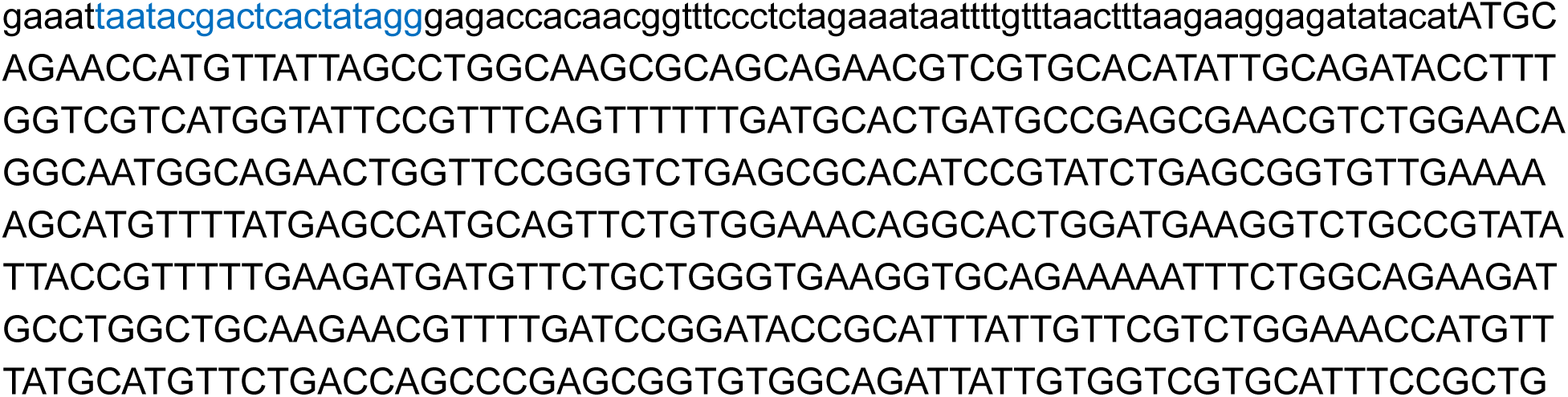

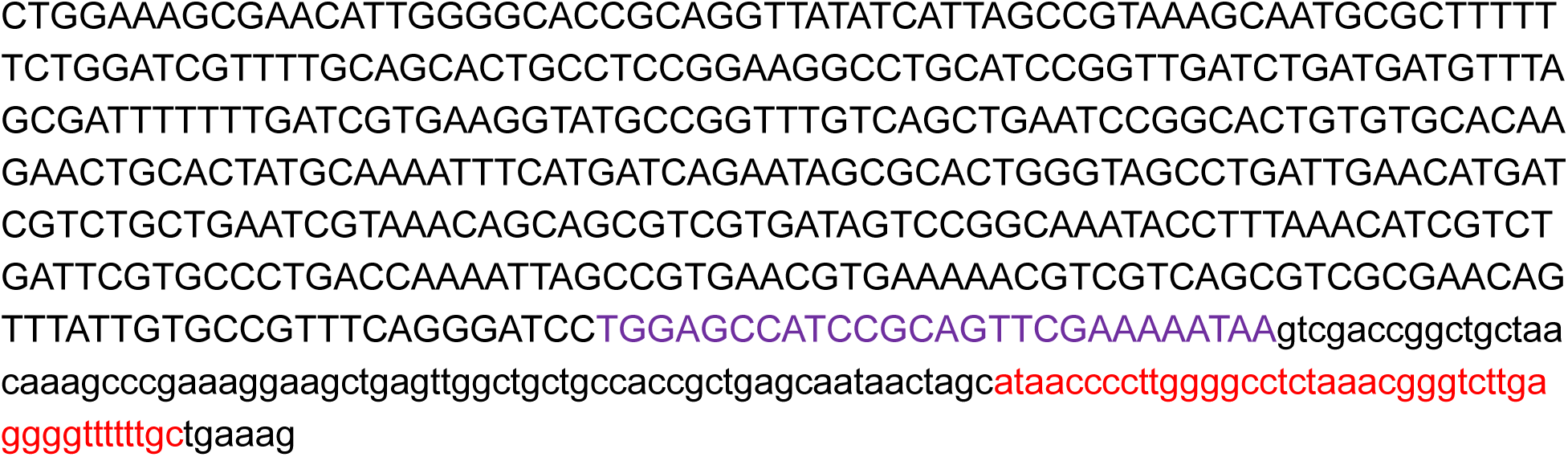

## DNA Sequence of Btβ4GalT1 in pJL1 Context

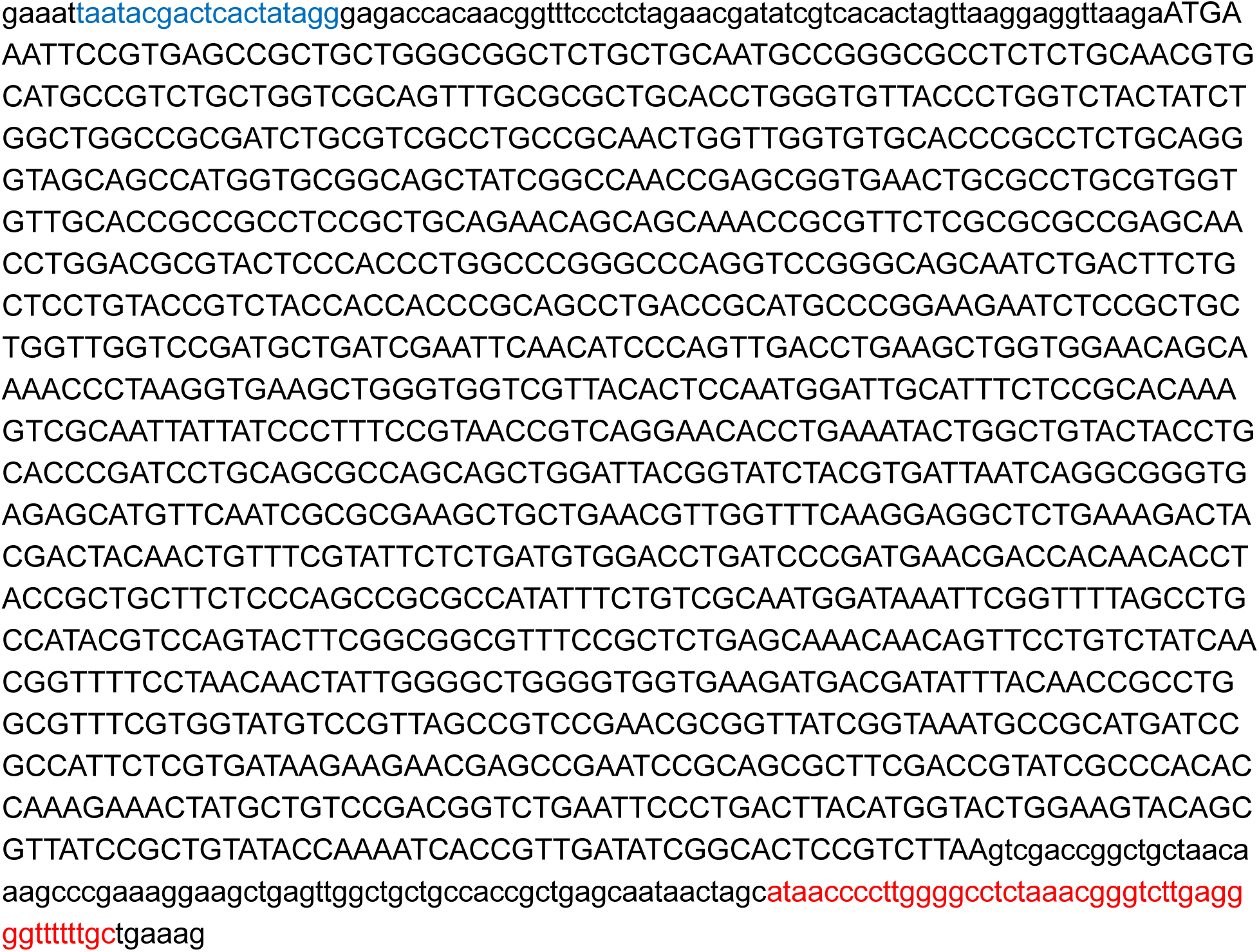

## DNA Sequence of NgLgtB in pJL1 Context

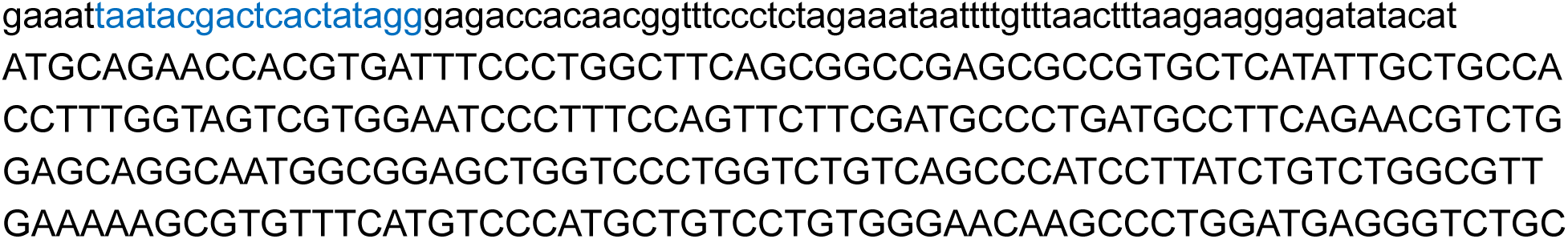

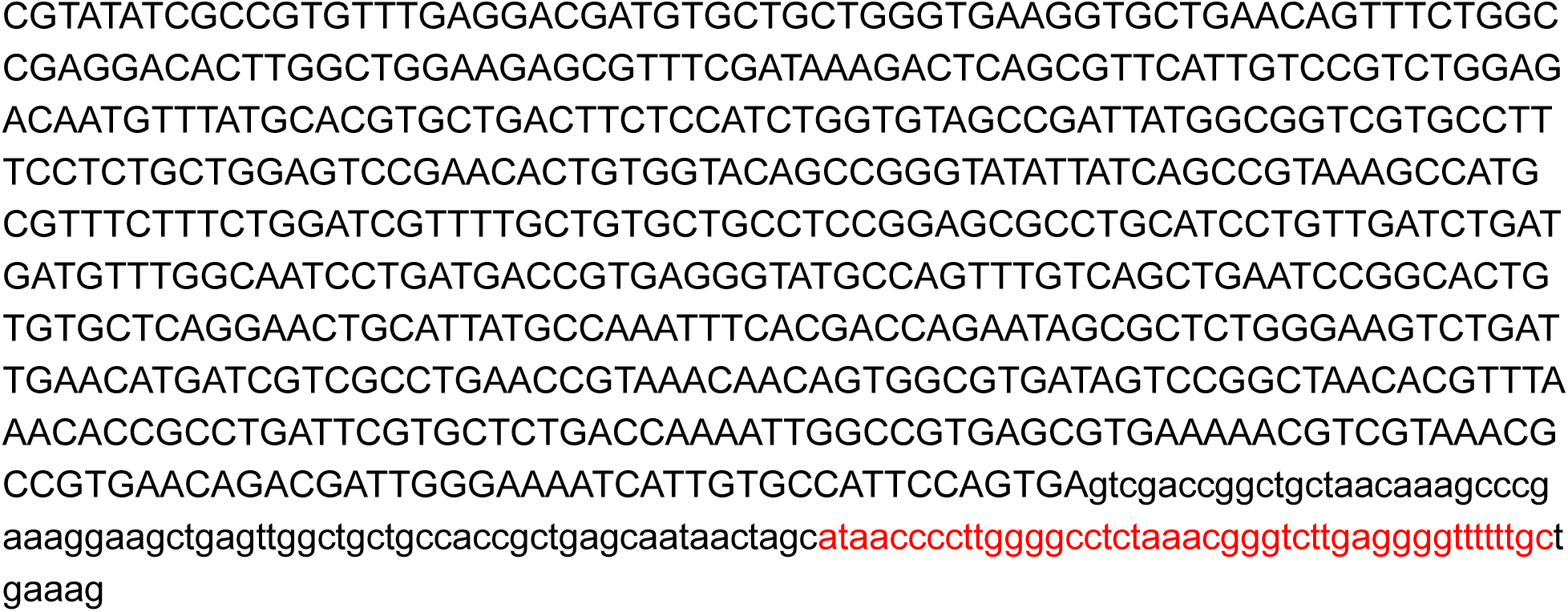

## DNA Sequence of SpWchJ in pJL1 Context

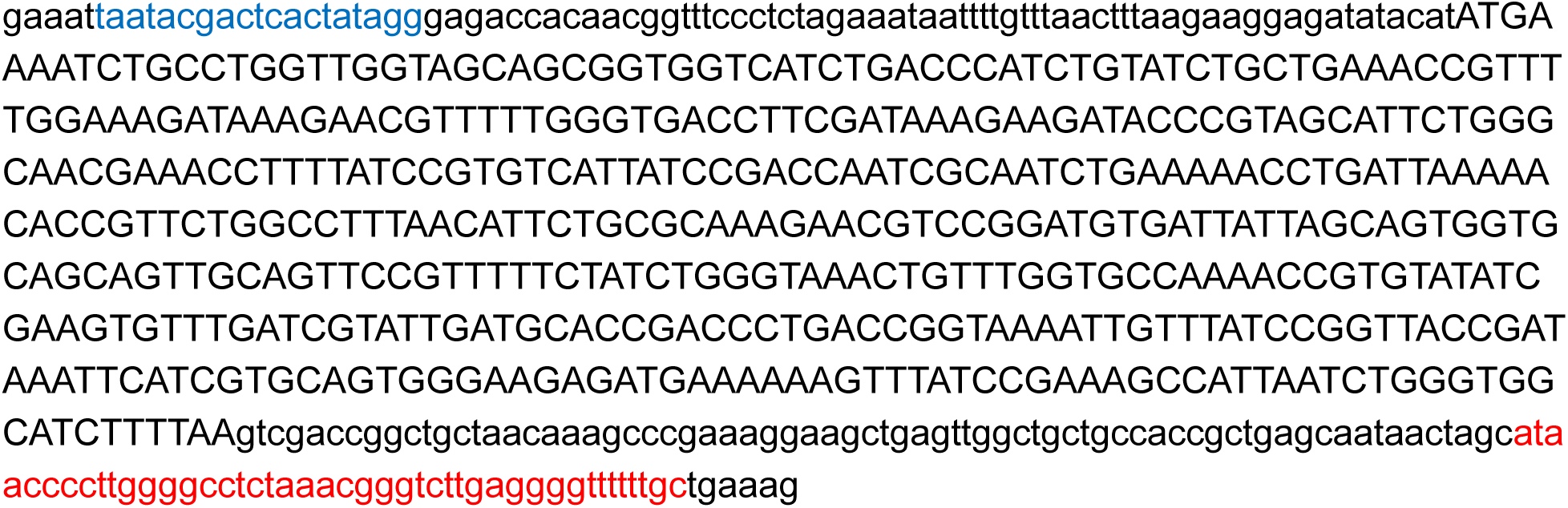

## DNA Sequence of SpWchK in pJL1 Context

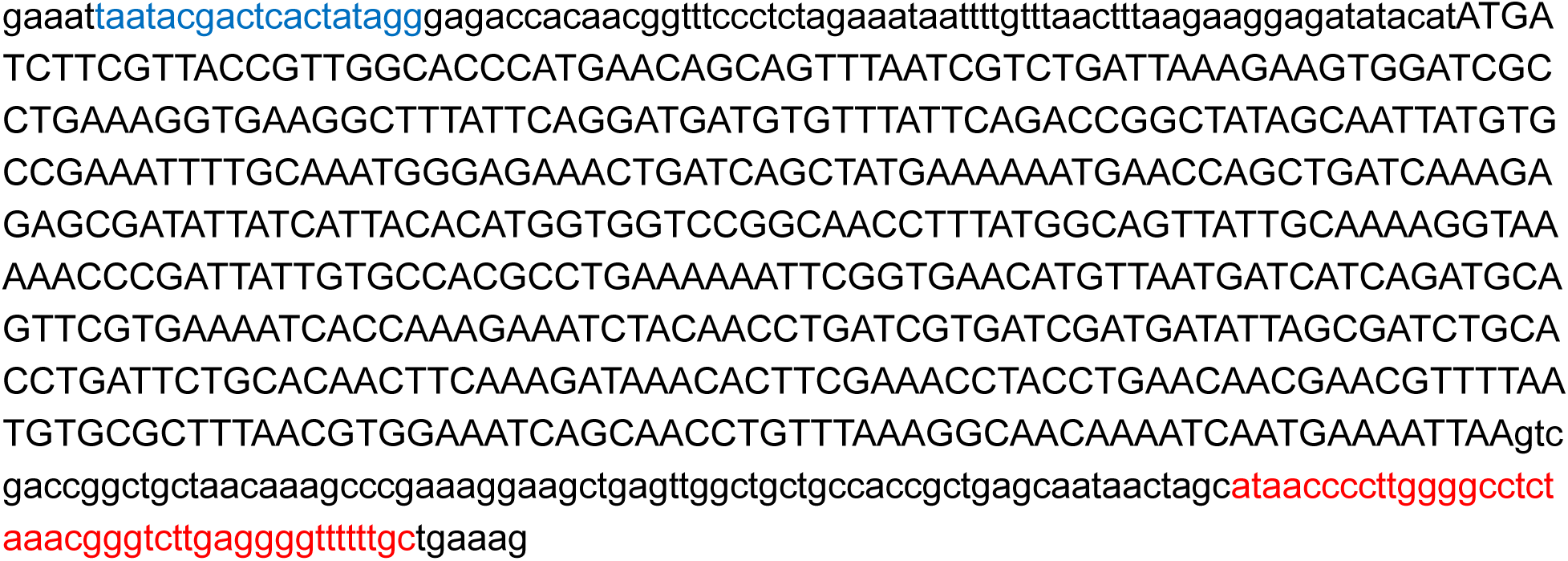

## DNA Sequence of NmLgtC in pJL1 Context

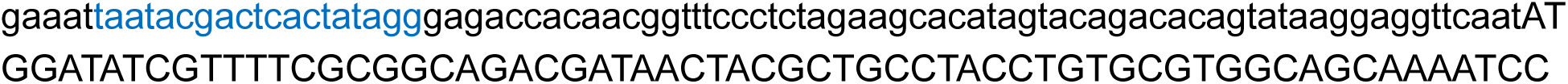

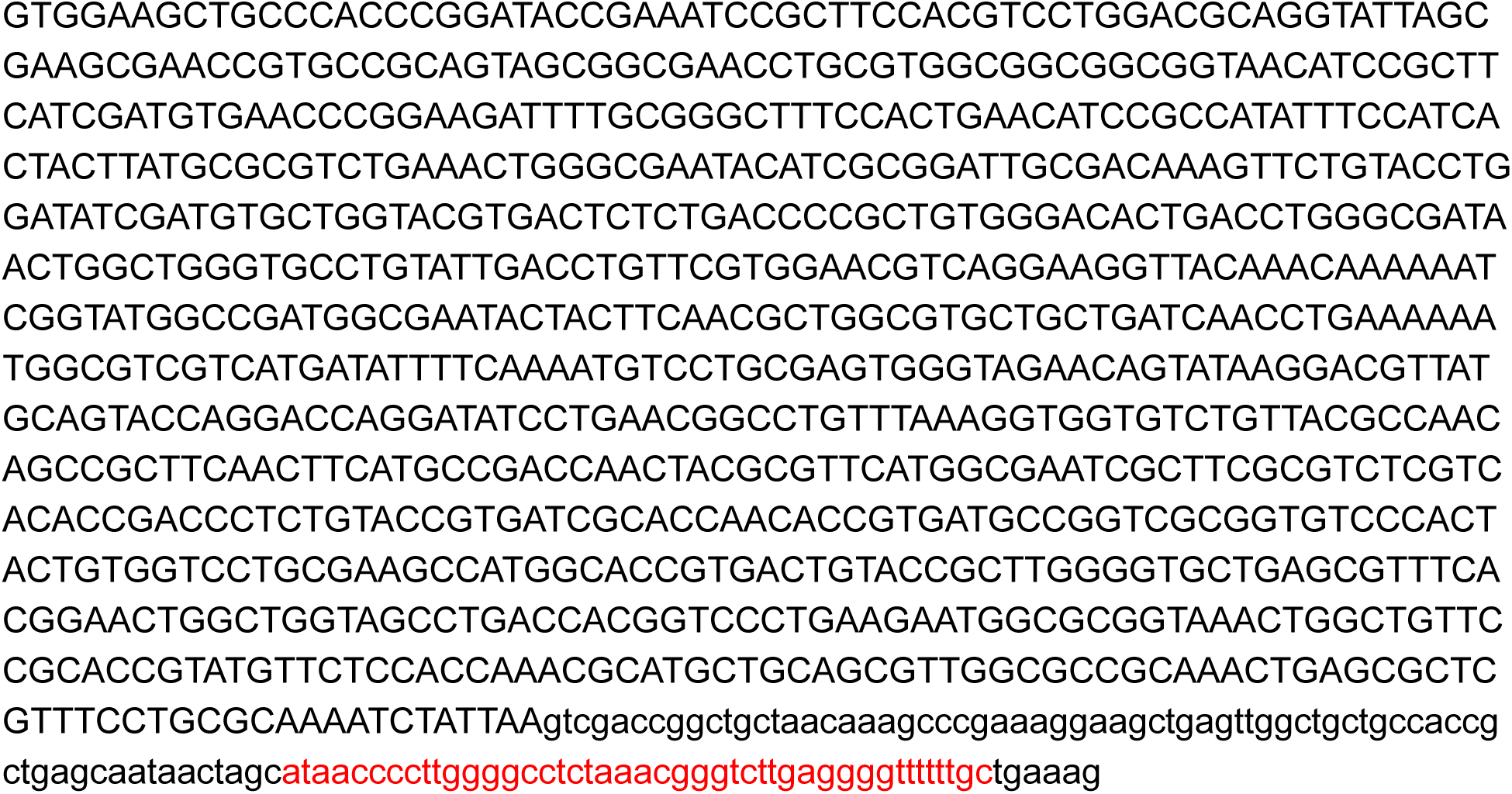

## DNA Sequence of HsSIAT1 in pJL1 Context

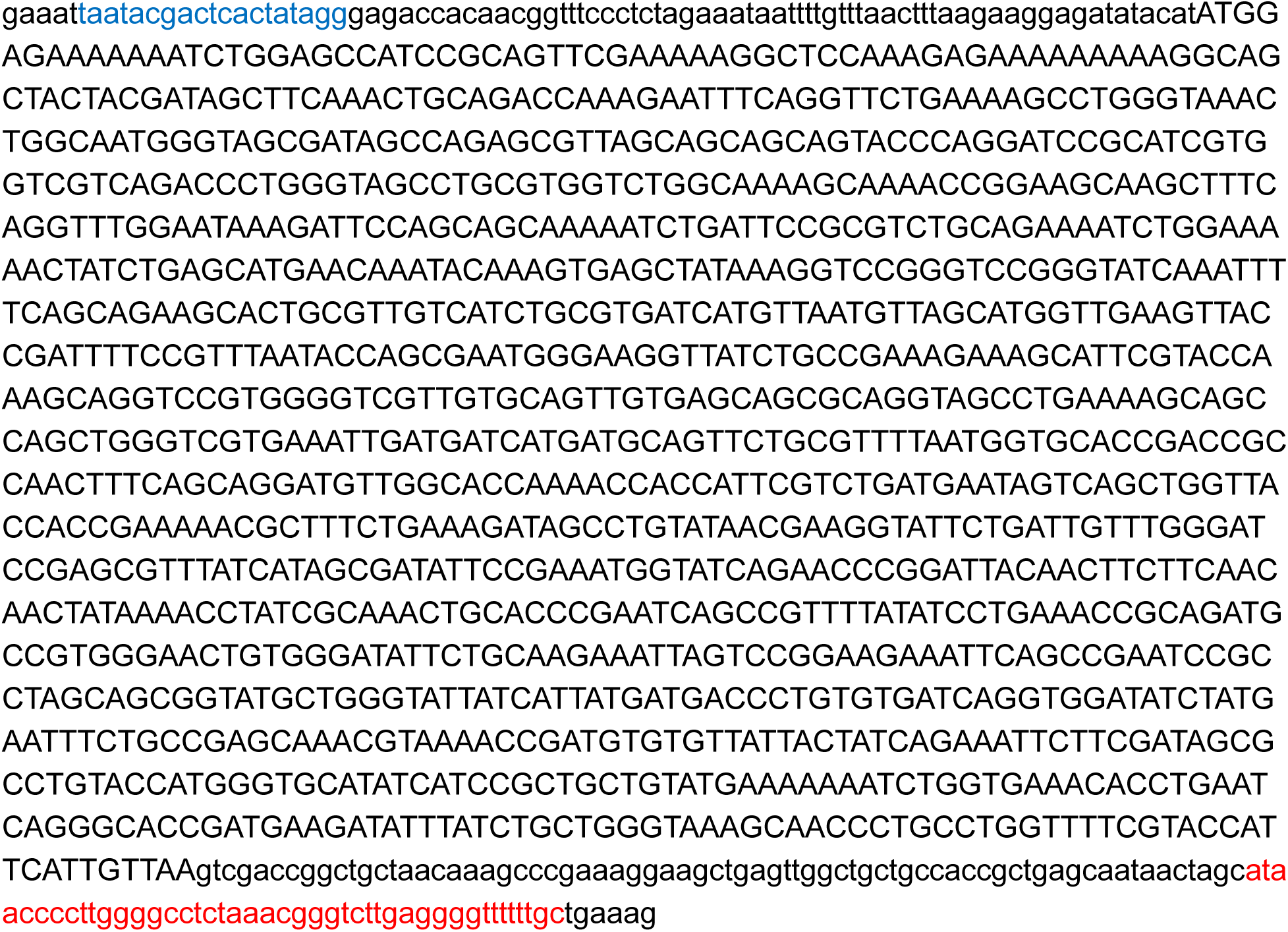

## DNA Sequence of PmST3,6 in pJL1 Context

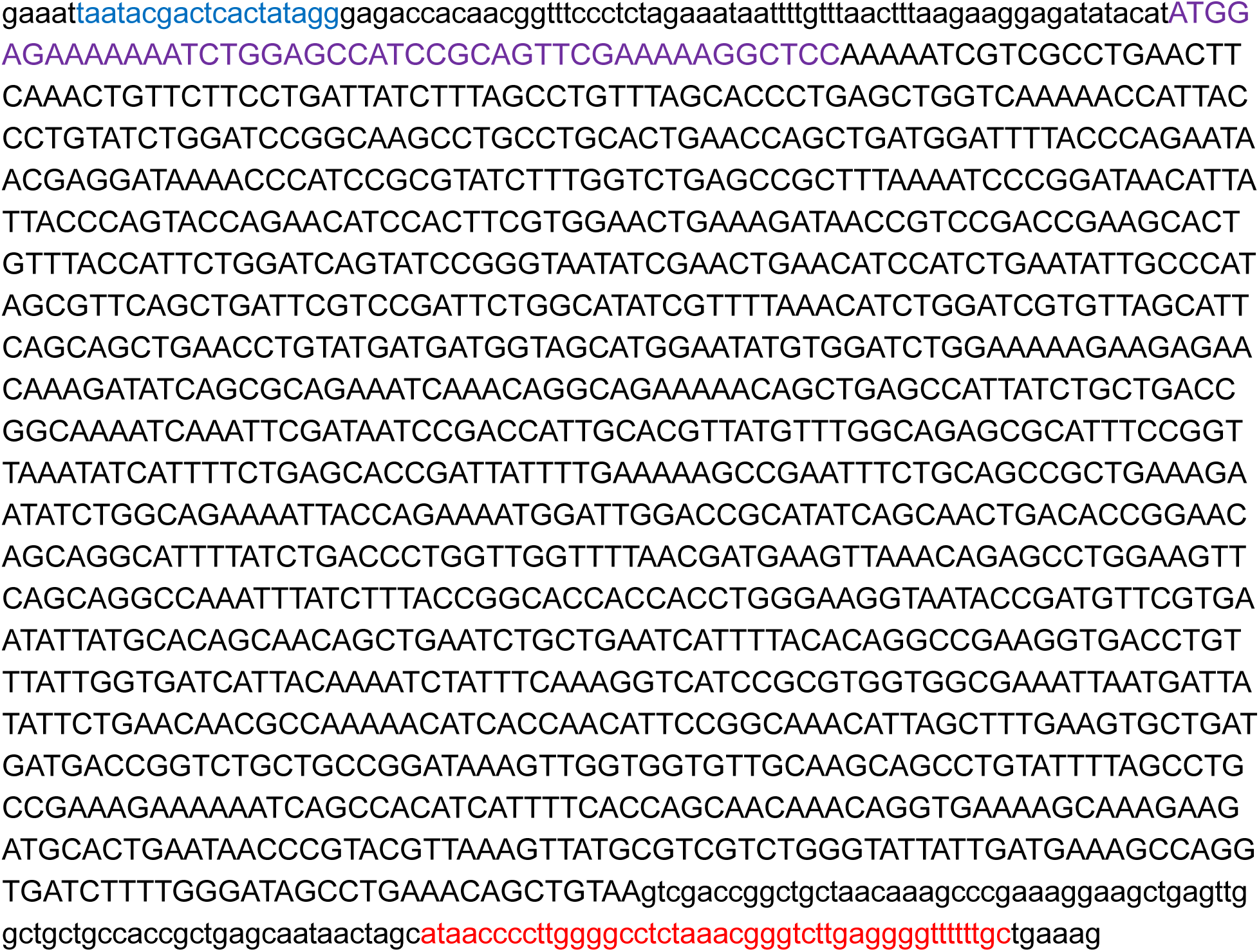

## DNA Sequence of CjCST-I in pJL1 Context

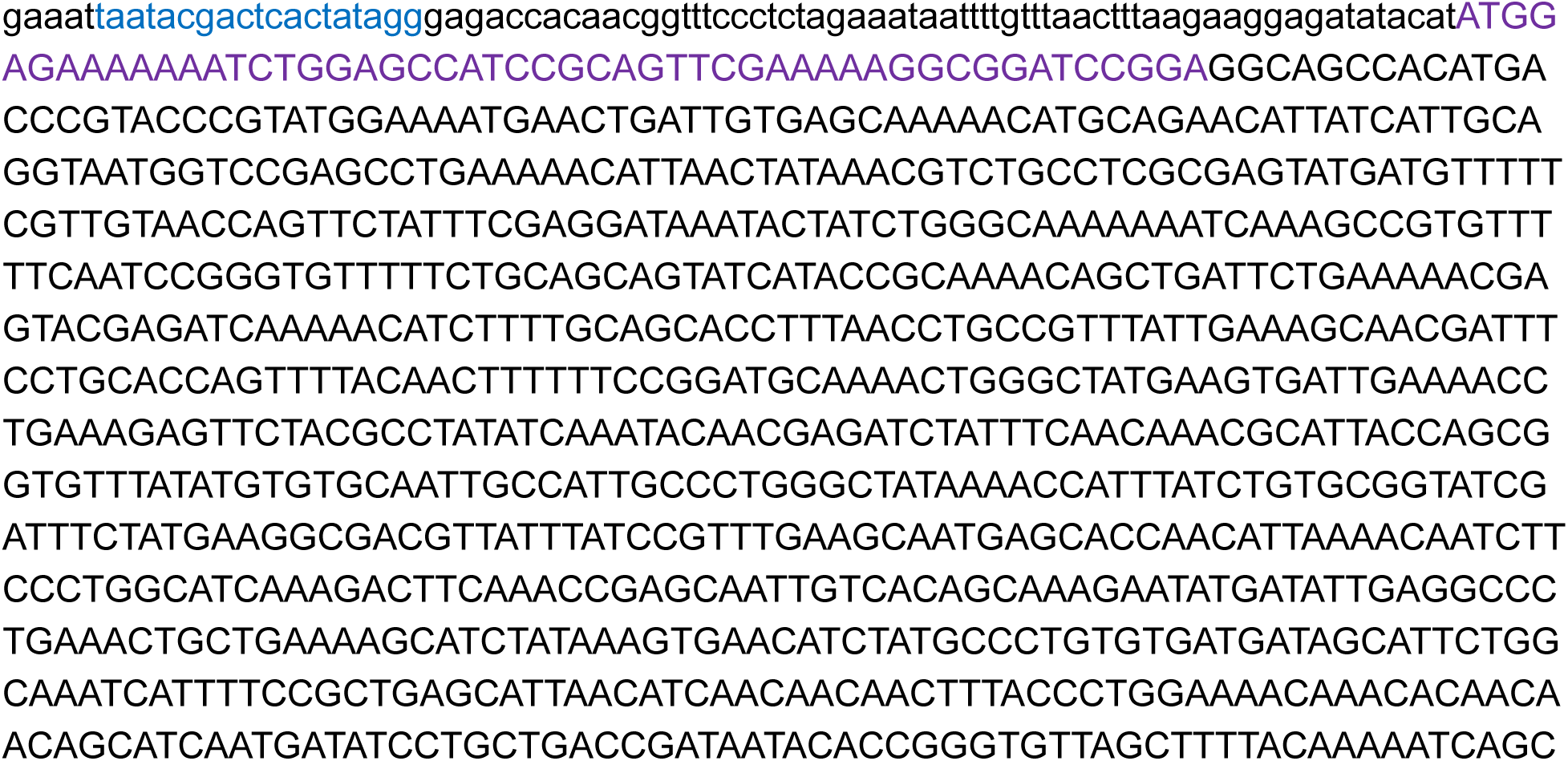

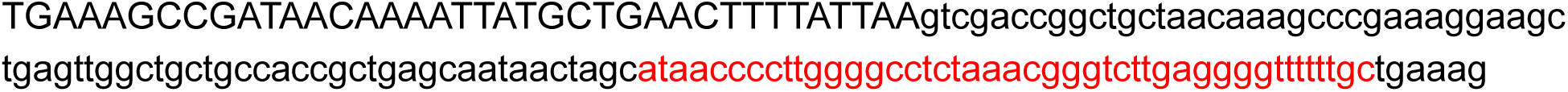

## DNA Sequence of CjCST-II in pJL1 Context

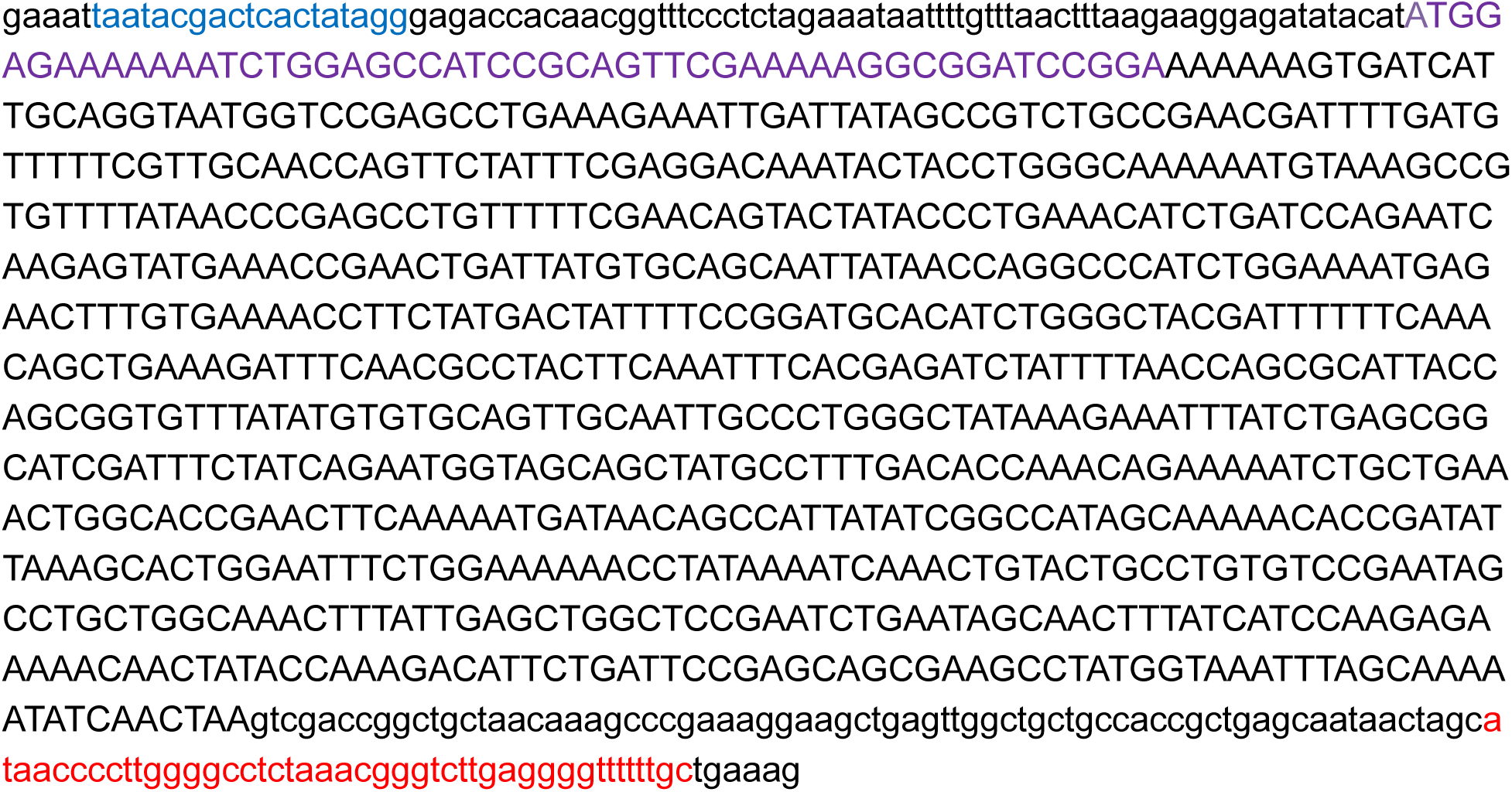

## DNA Sequence of SpPvg1 in pJL1 Context

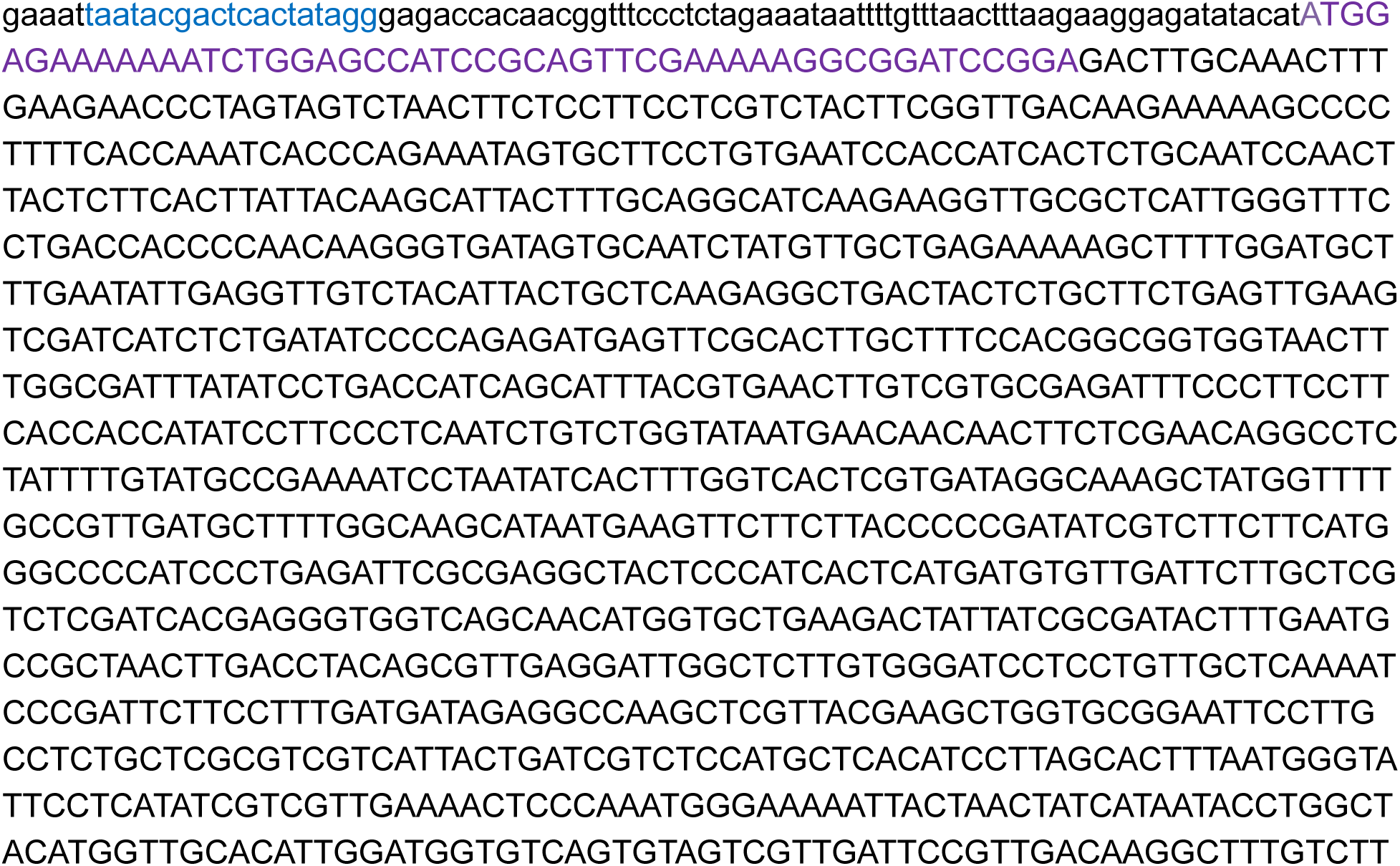

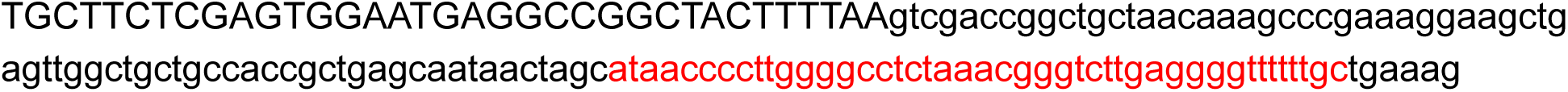

## DNA Sequence of VsST3 in pJL1 Context

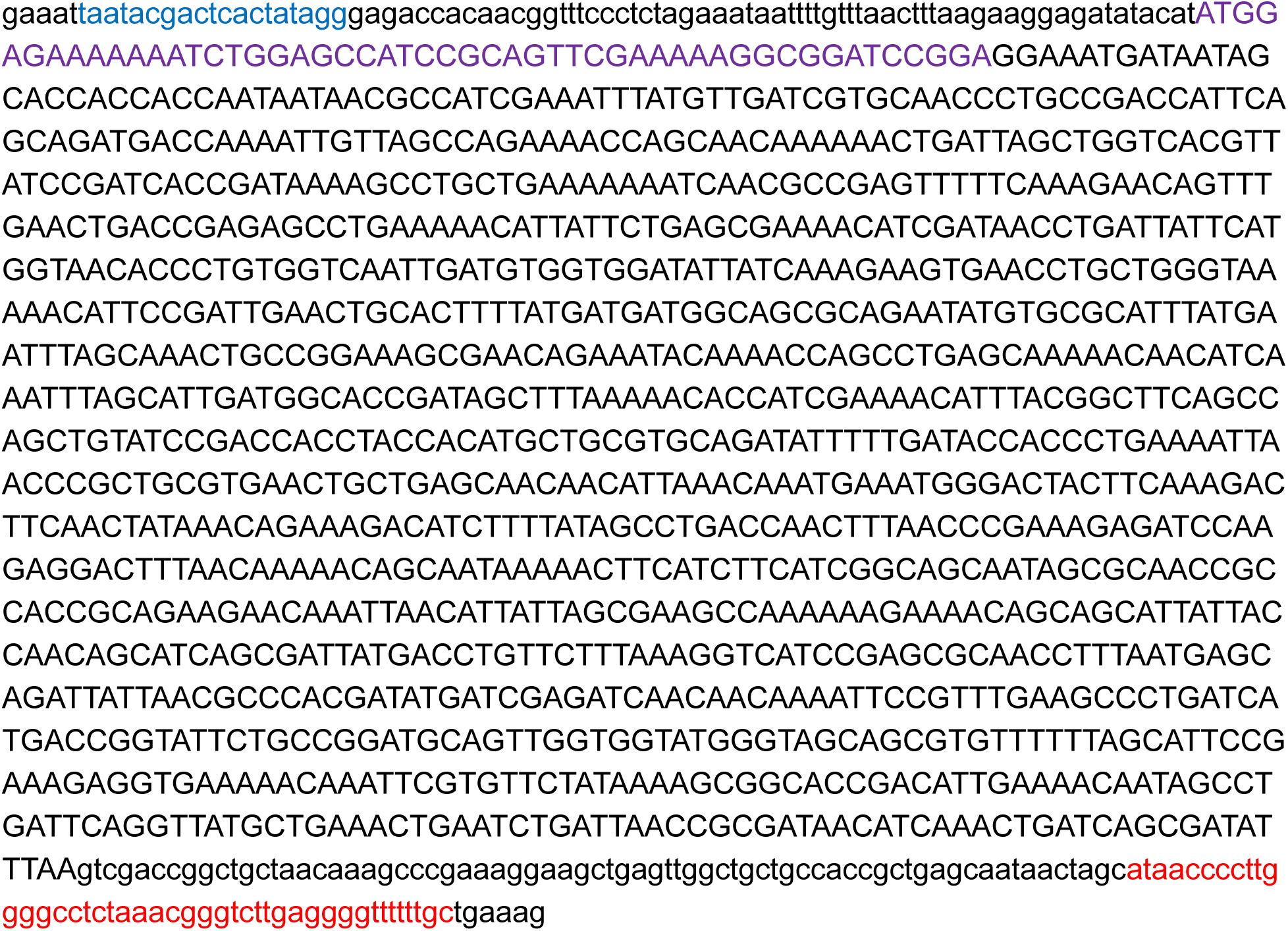

## DNA Sequence of HpFutA in pJL1 Context

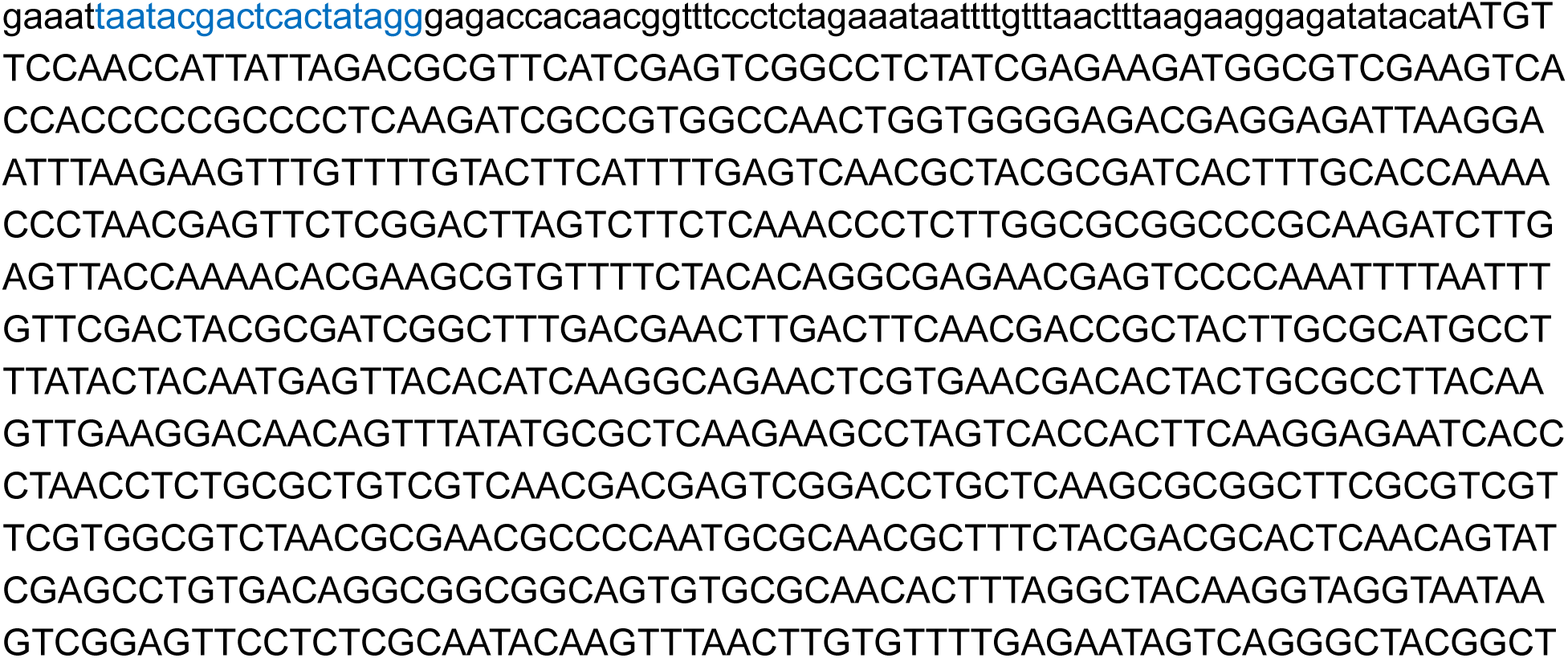

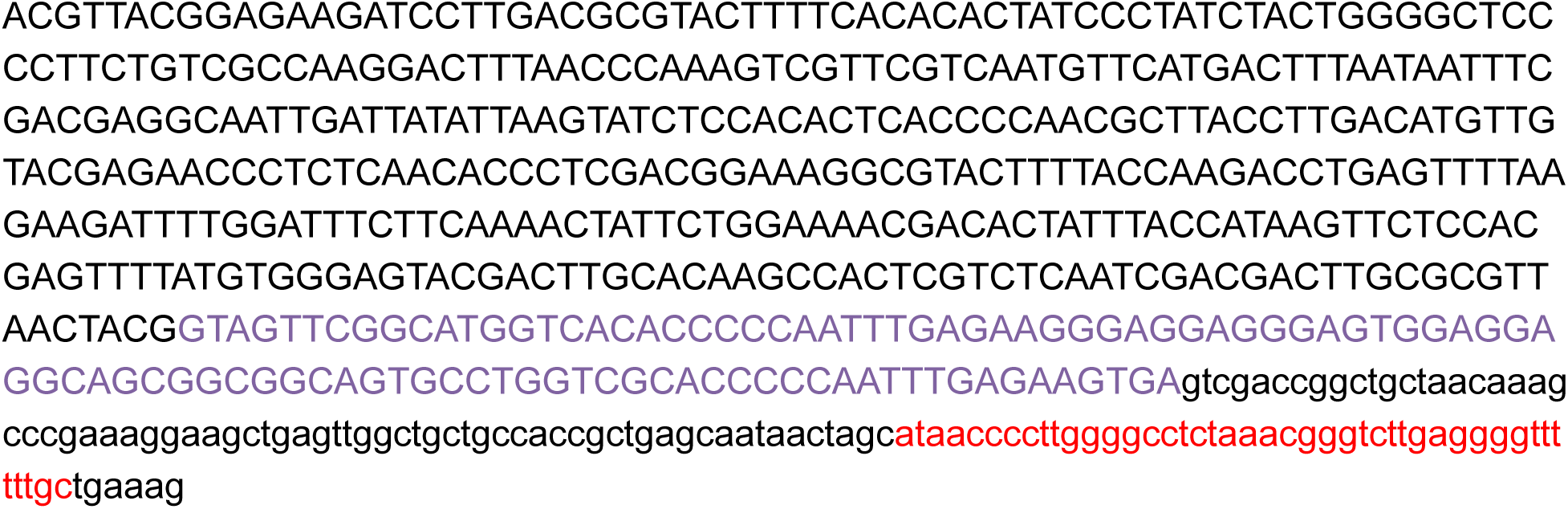

## DNA Sequence of HpFutC in pJL1 Context

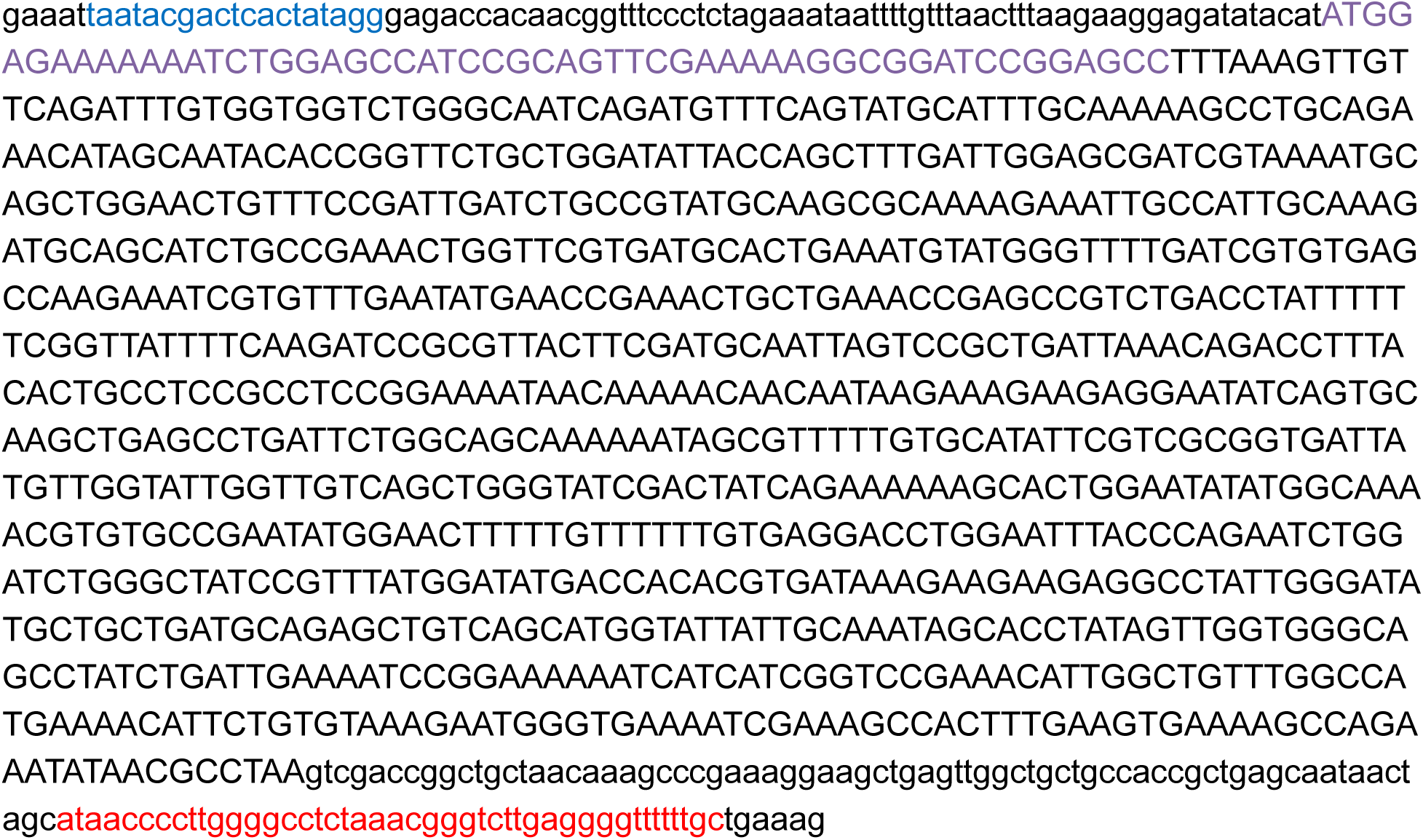

## DNA Sequence of BtGGTA in pJL1 Context

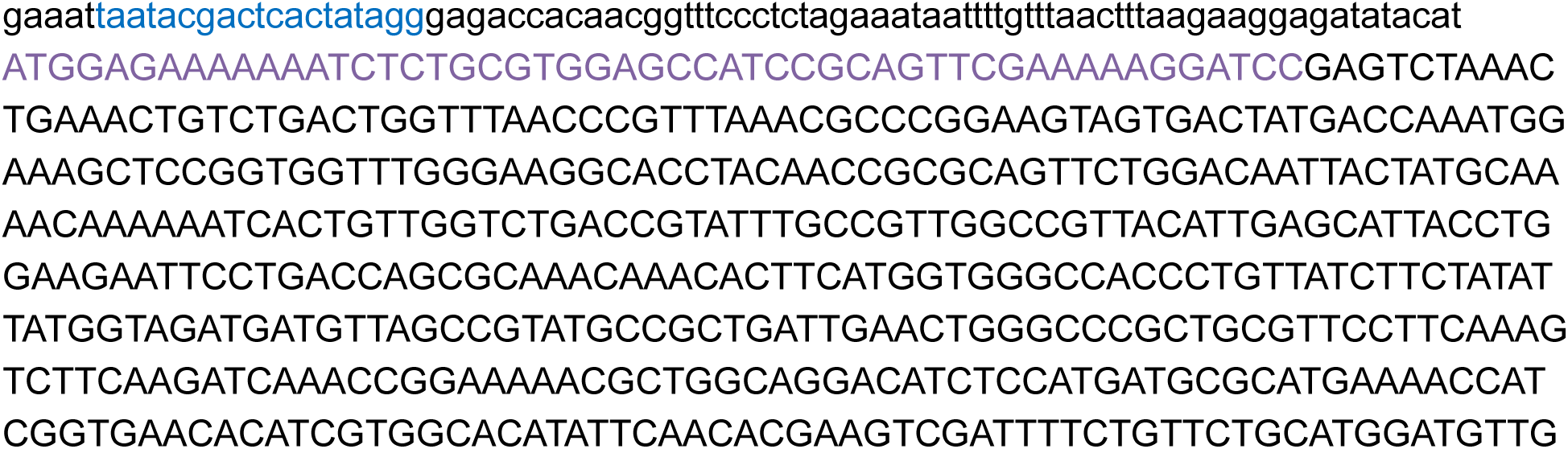

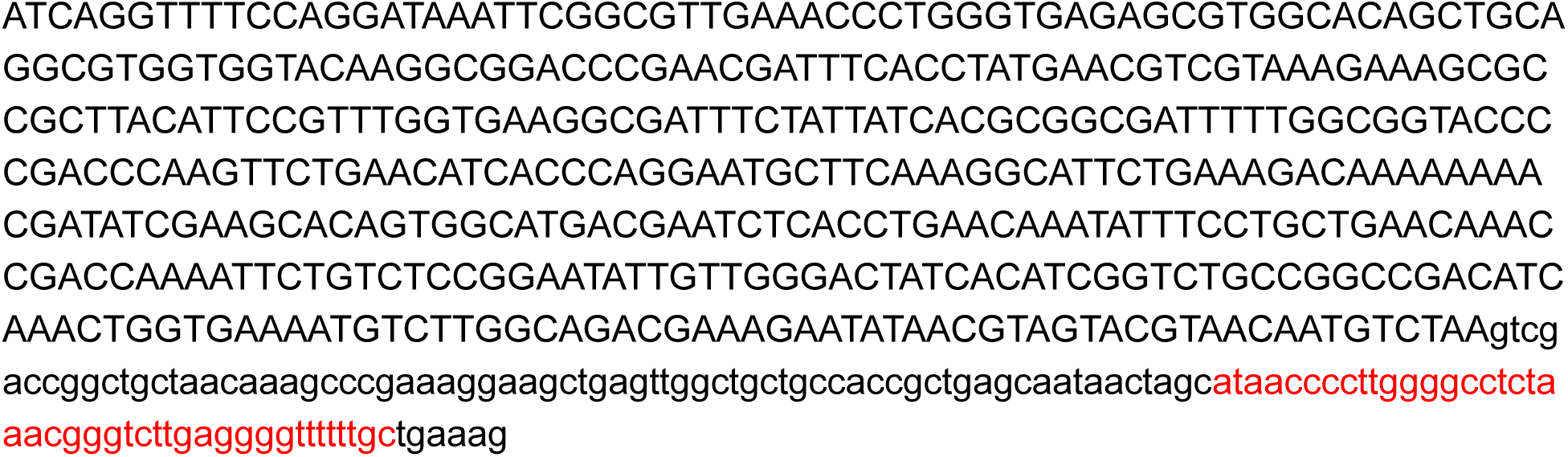

## DNA Sequence of HdGlcNAcT in pJL1 Context

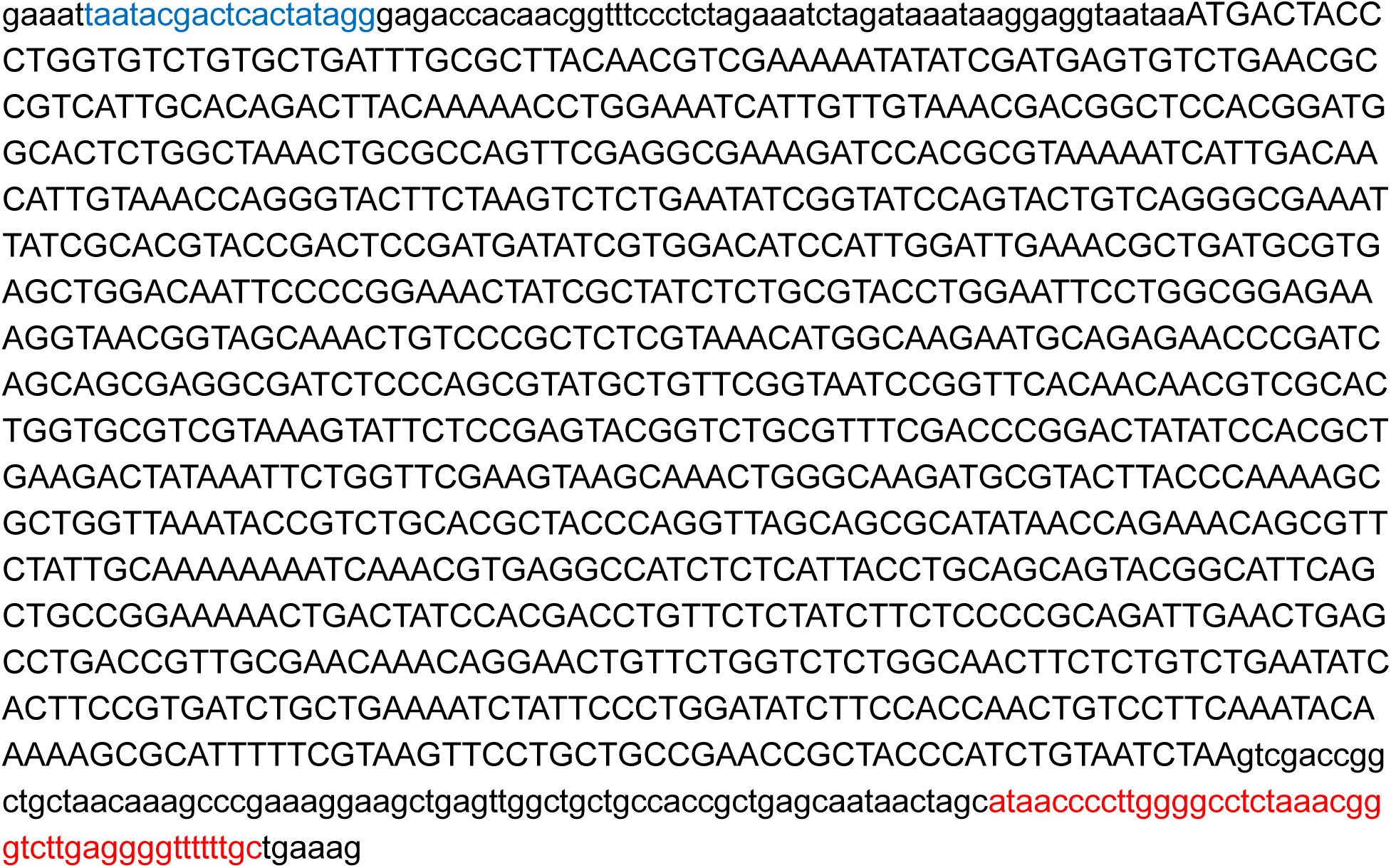

## DNA Sequence of NgLgtA in pJL1 Context

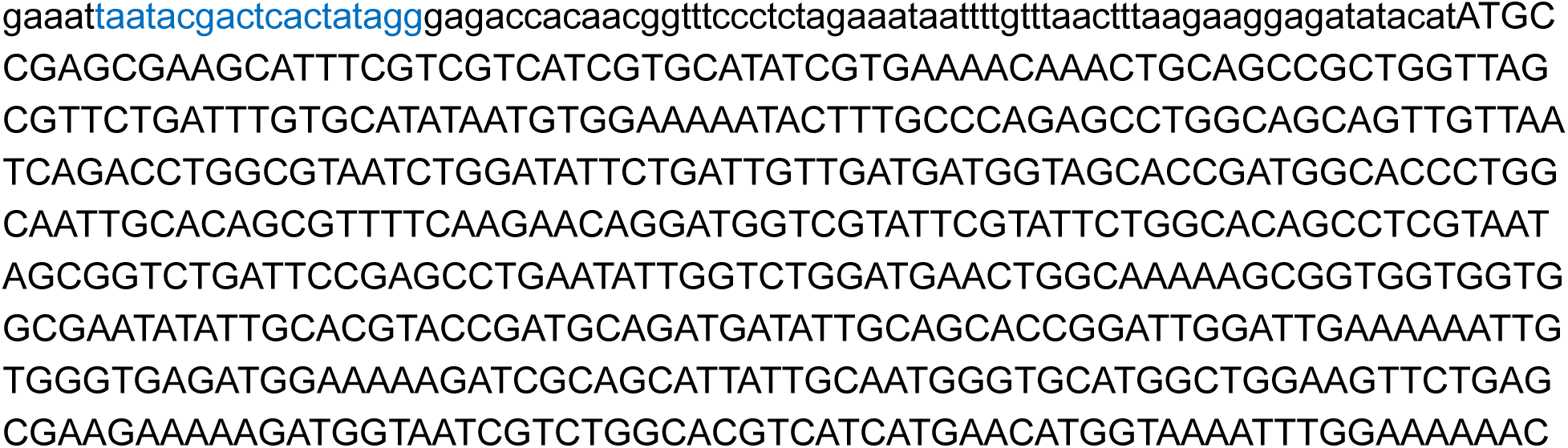

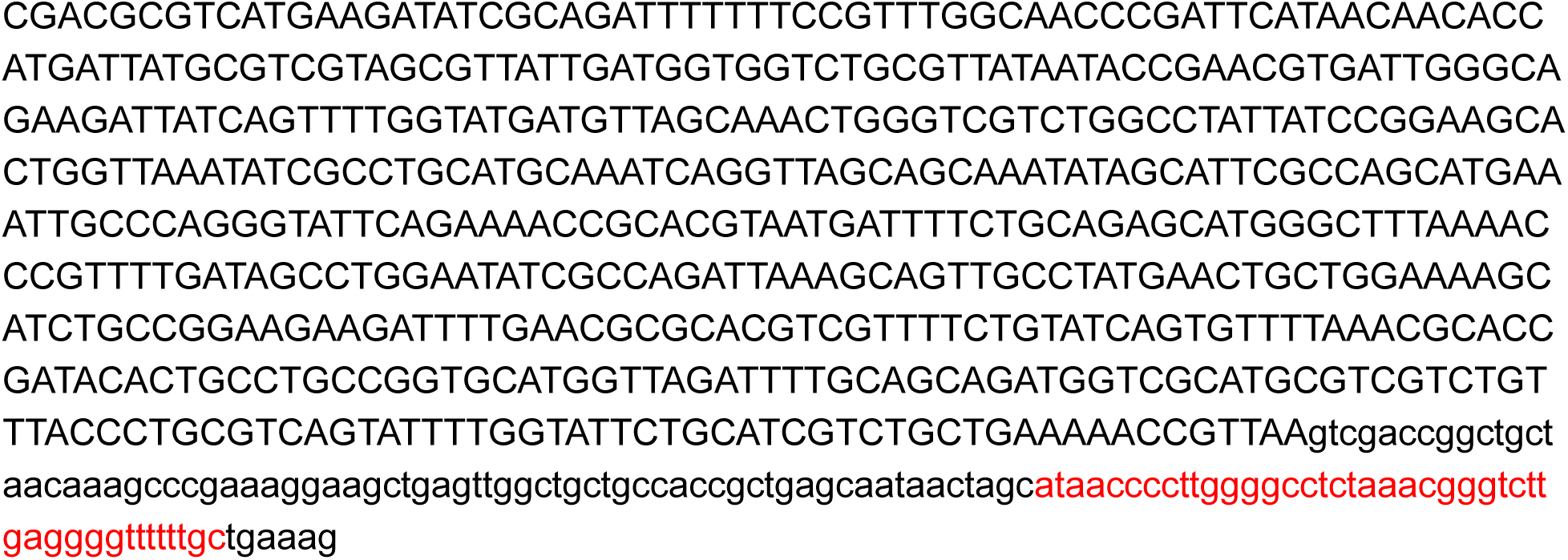

## DNA Sequence of PdST6 in pJL1 Context

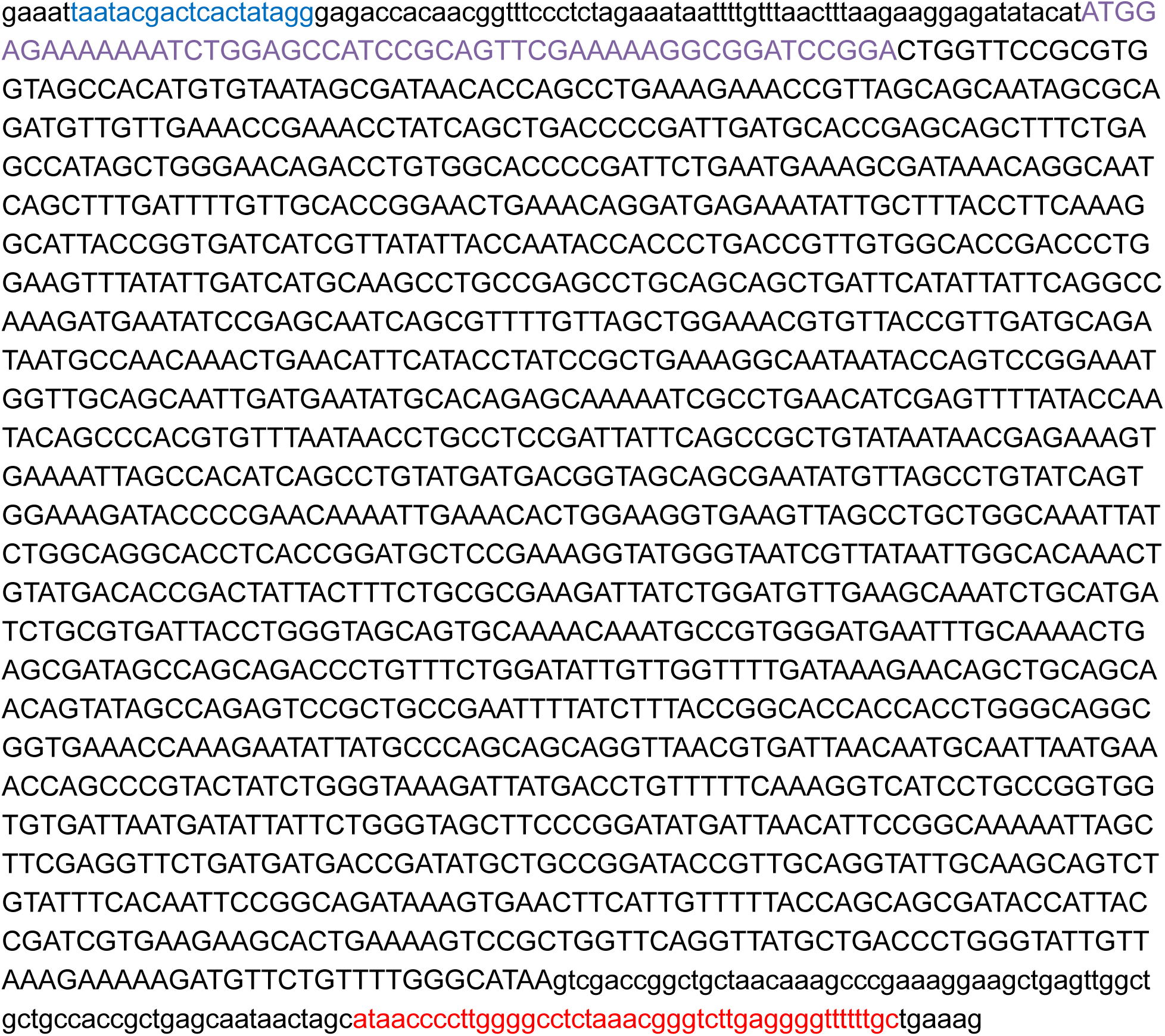

## DNA Sequence of PIST6 in pJL1 Context

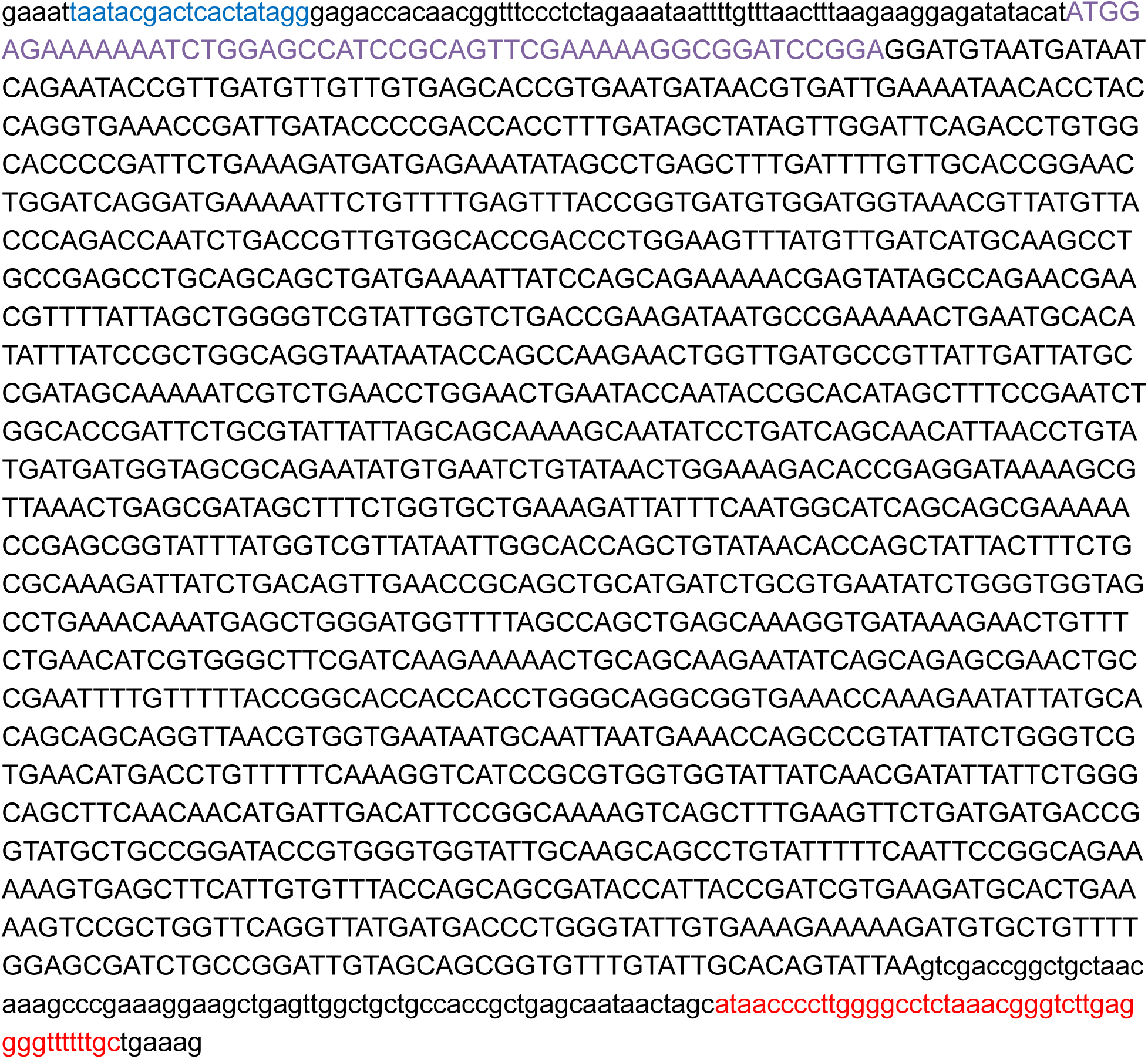

## DNA Sequence of PpST3 in pJL1 Context

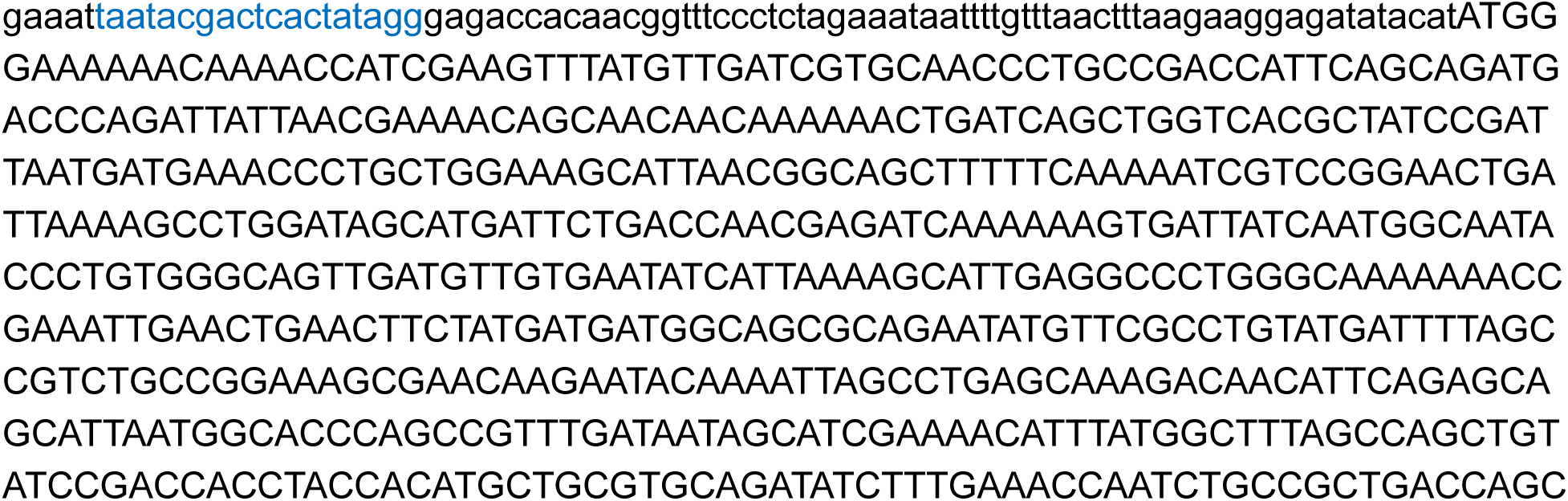

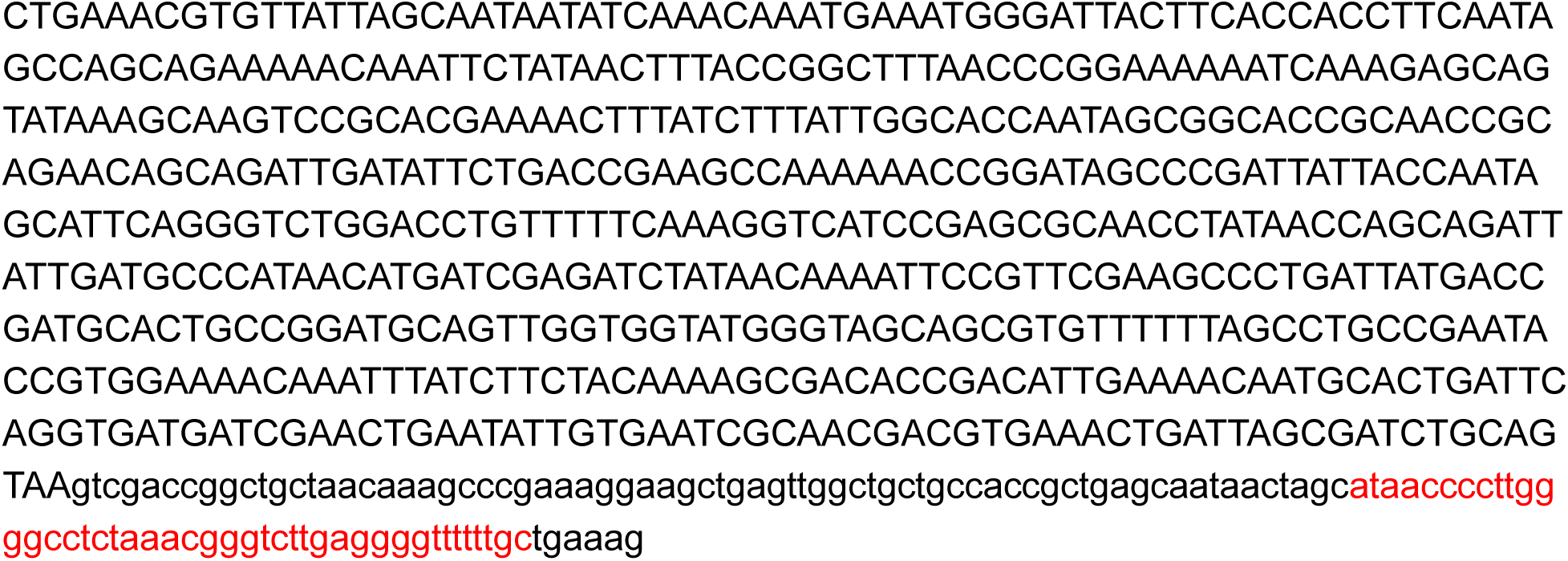

## DNA Sequence of H1HA10 in pJL1 Context

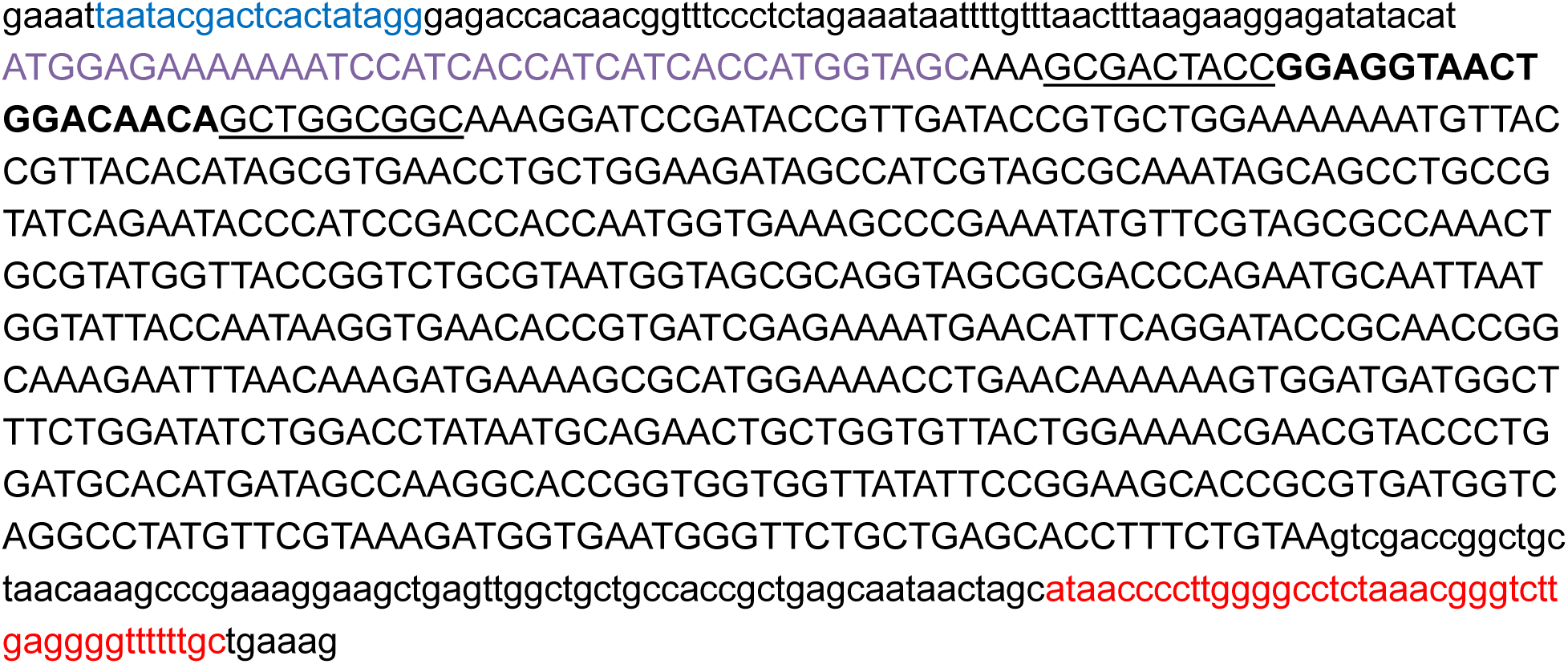

## DNA Sequence of ApNGT in pMAF10 Context

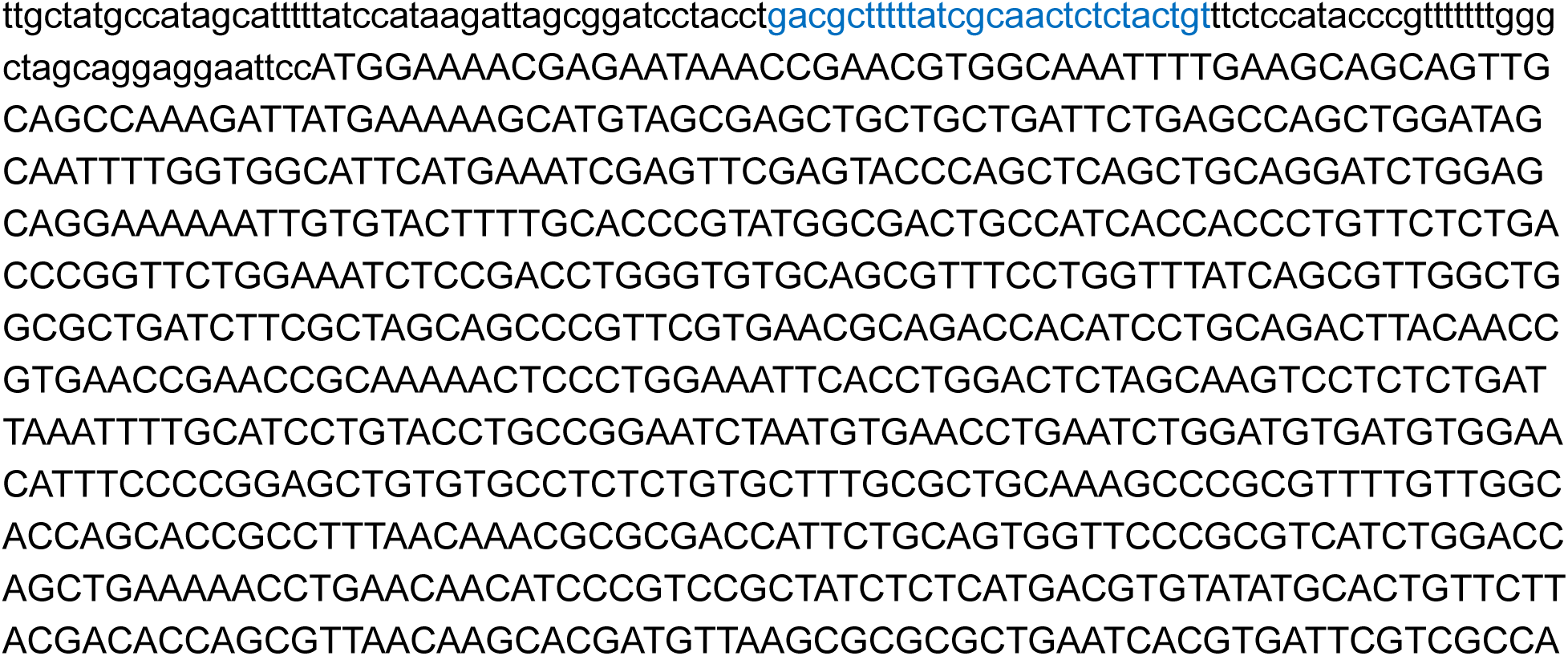

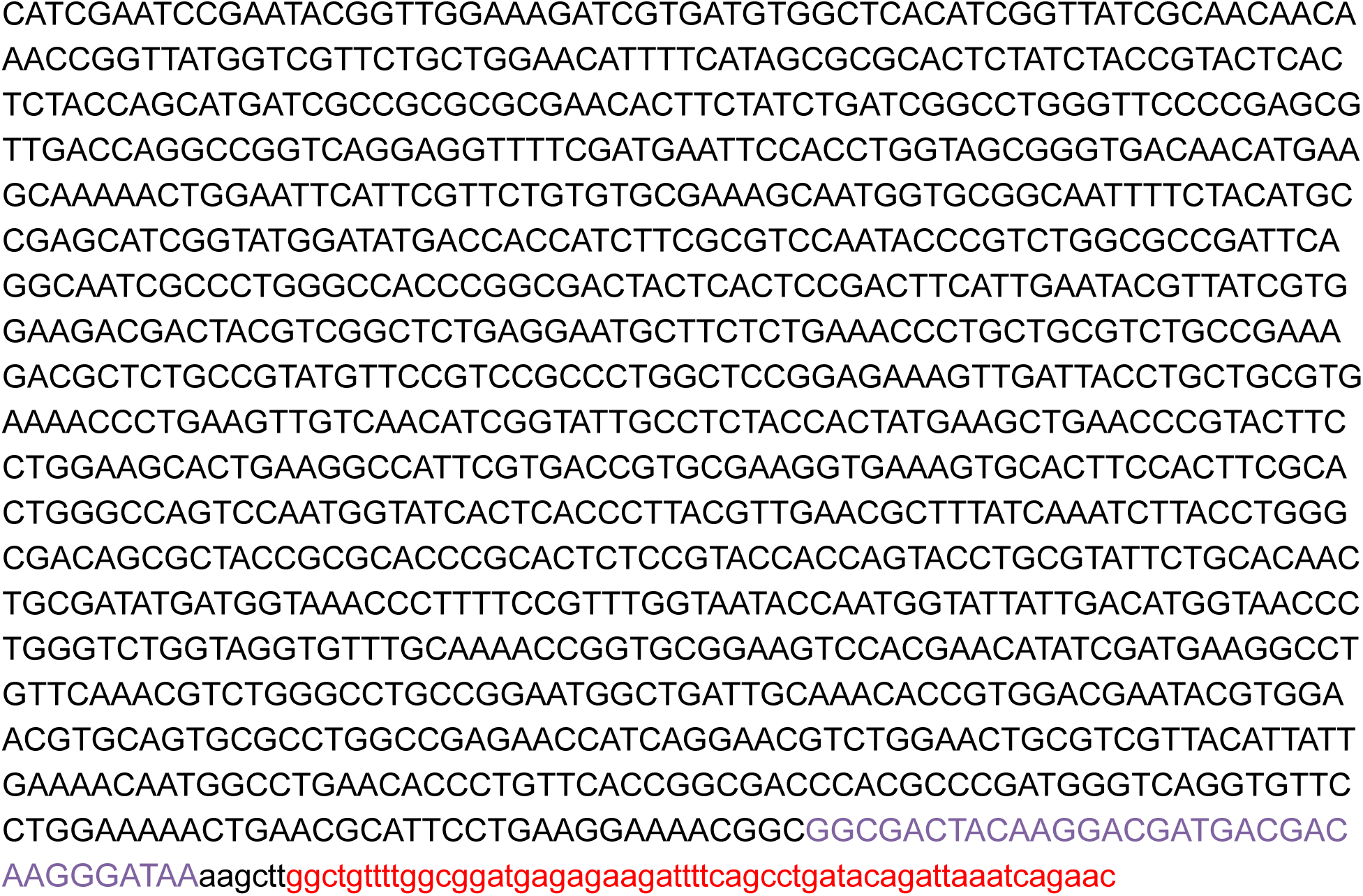

## DNA Sequence of NmLgtB.ApNGT in pMAF10 Context

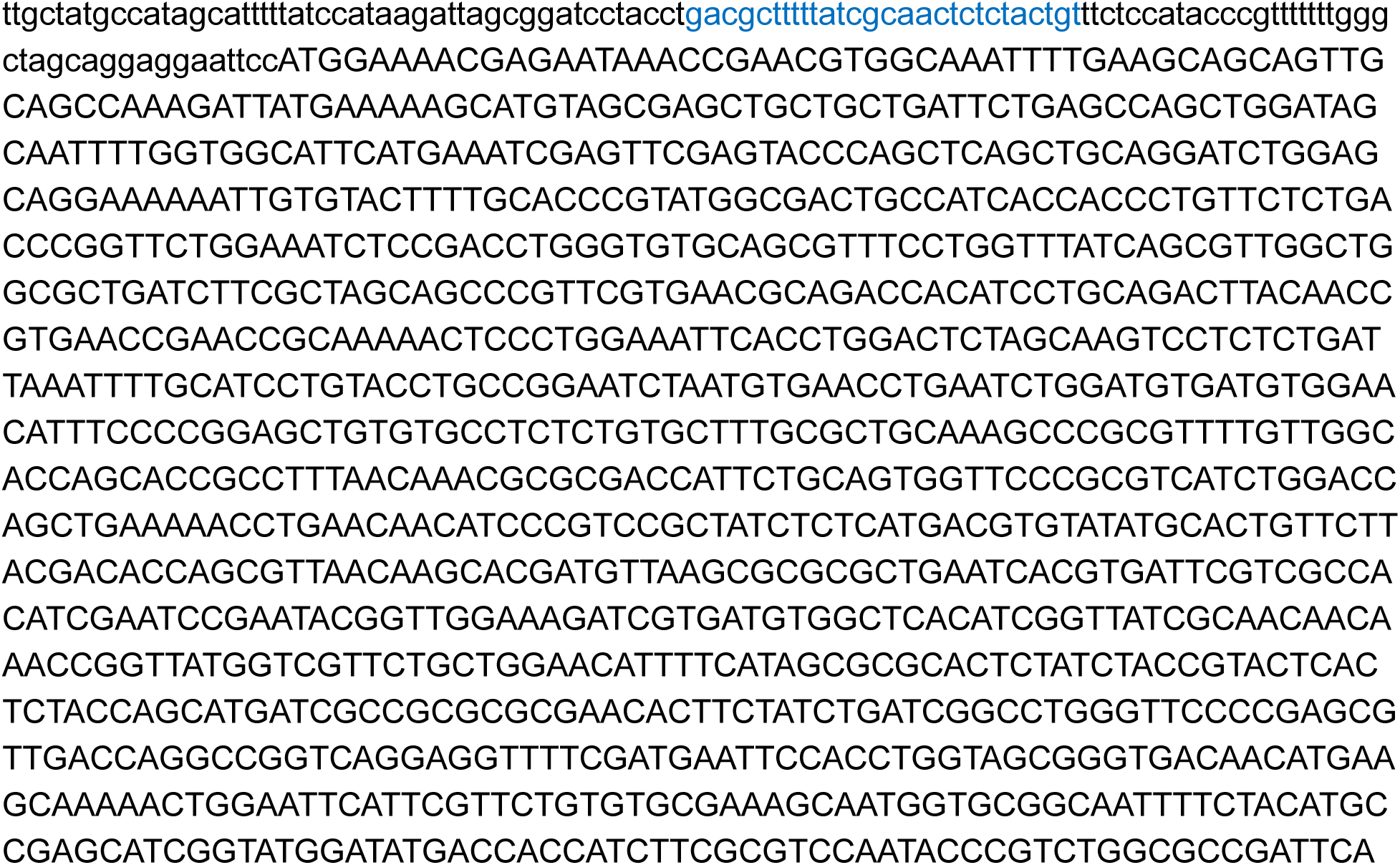

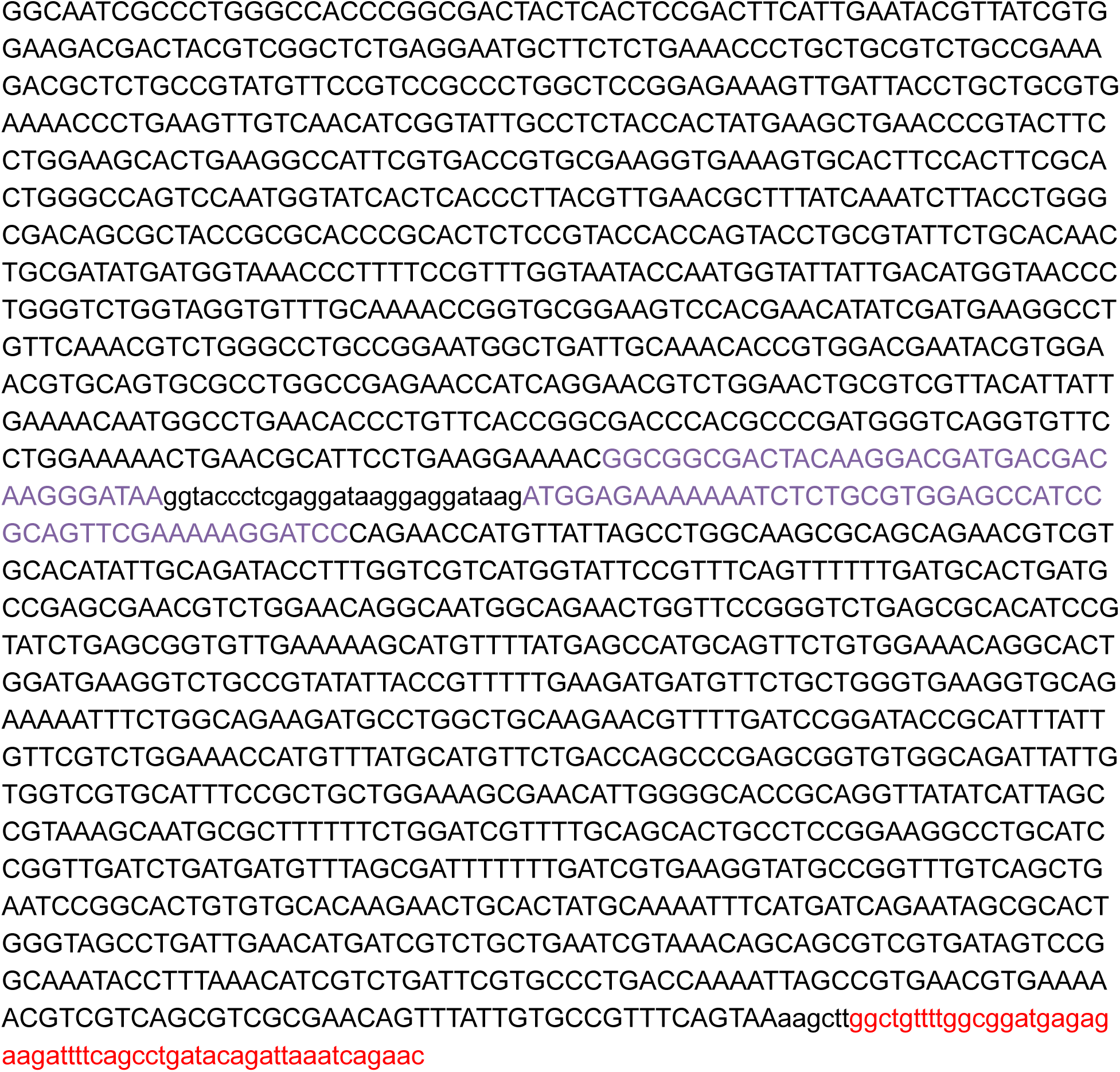

## DNA Sequence of CjCST-I.NmLgtB.ApNGT in pMAF10 Context

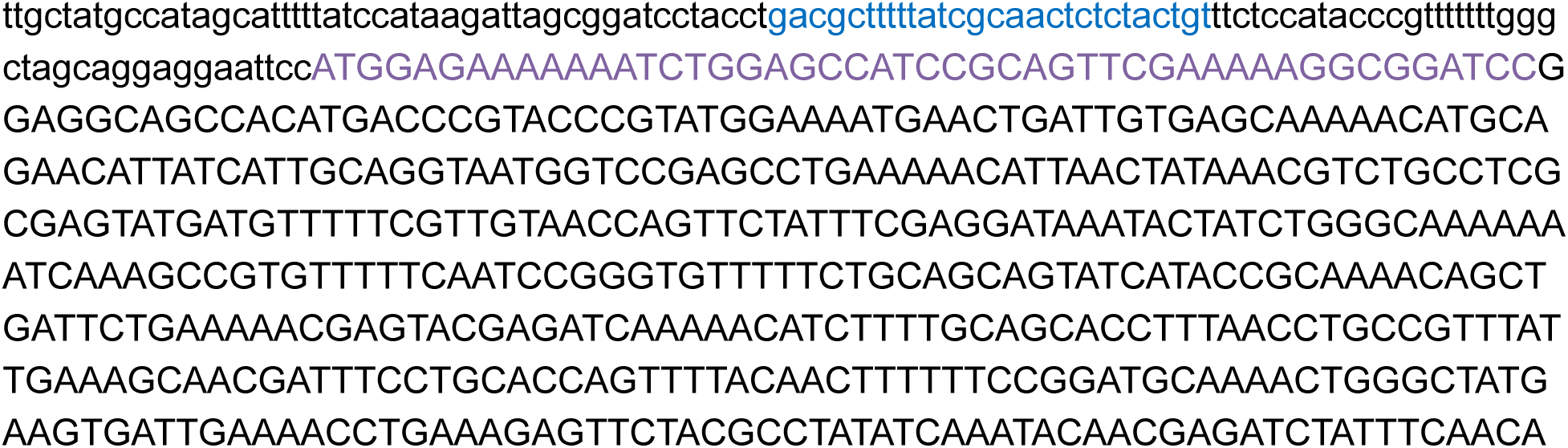

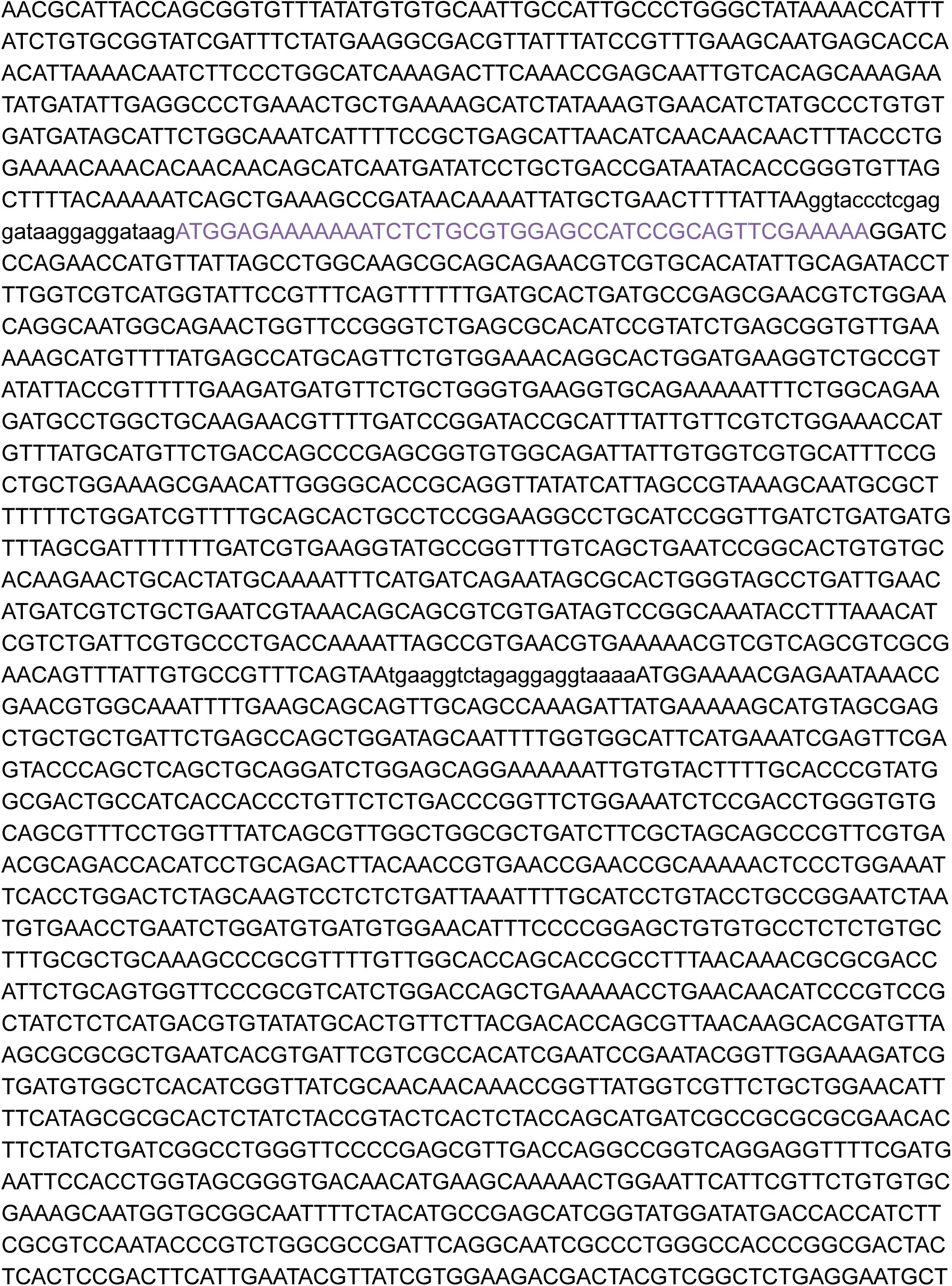

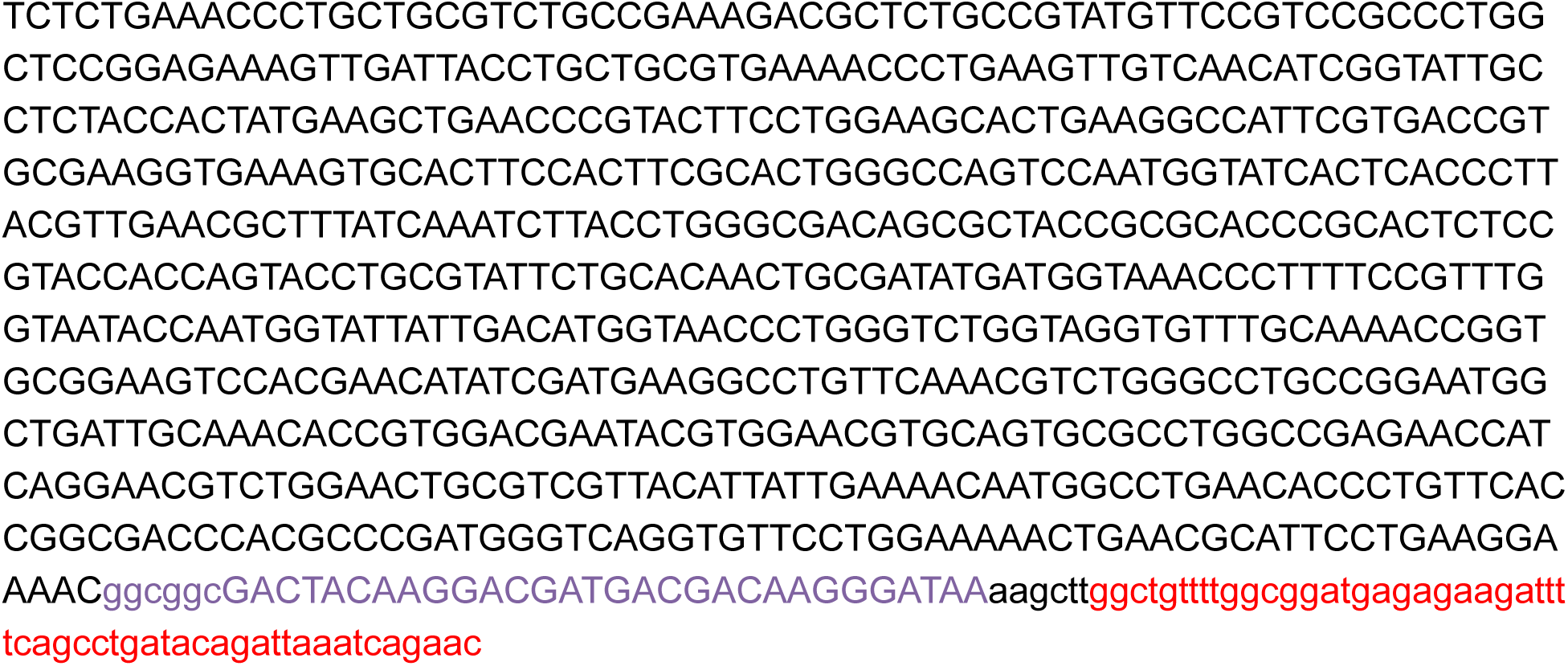

## DNA Sequence of PdST6.NmLgtB.ApNGT in pMAF10 Context

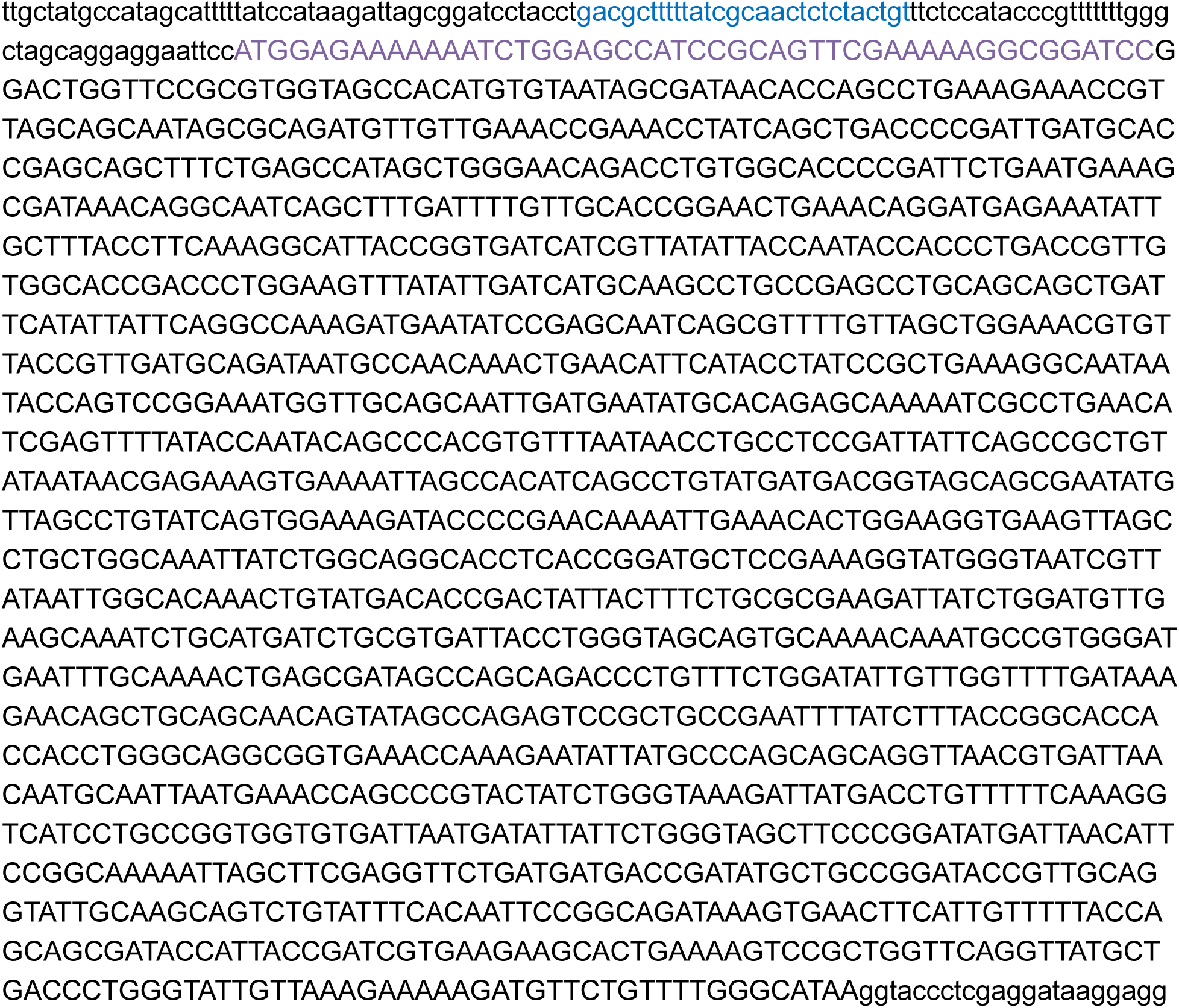

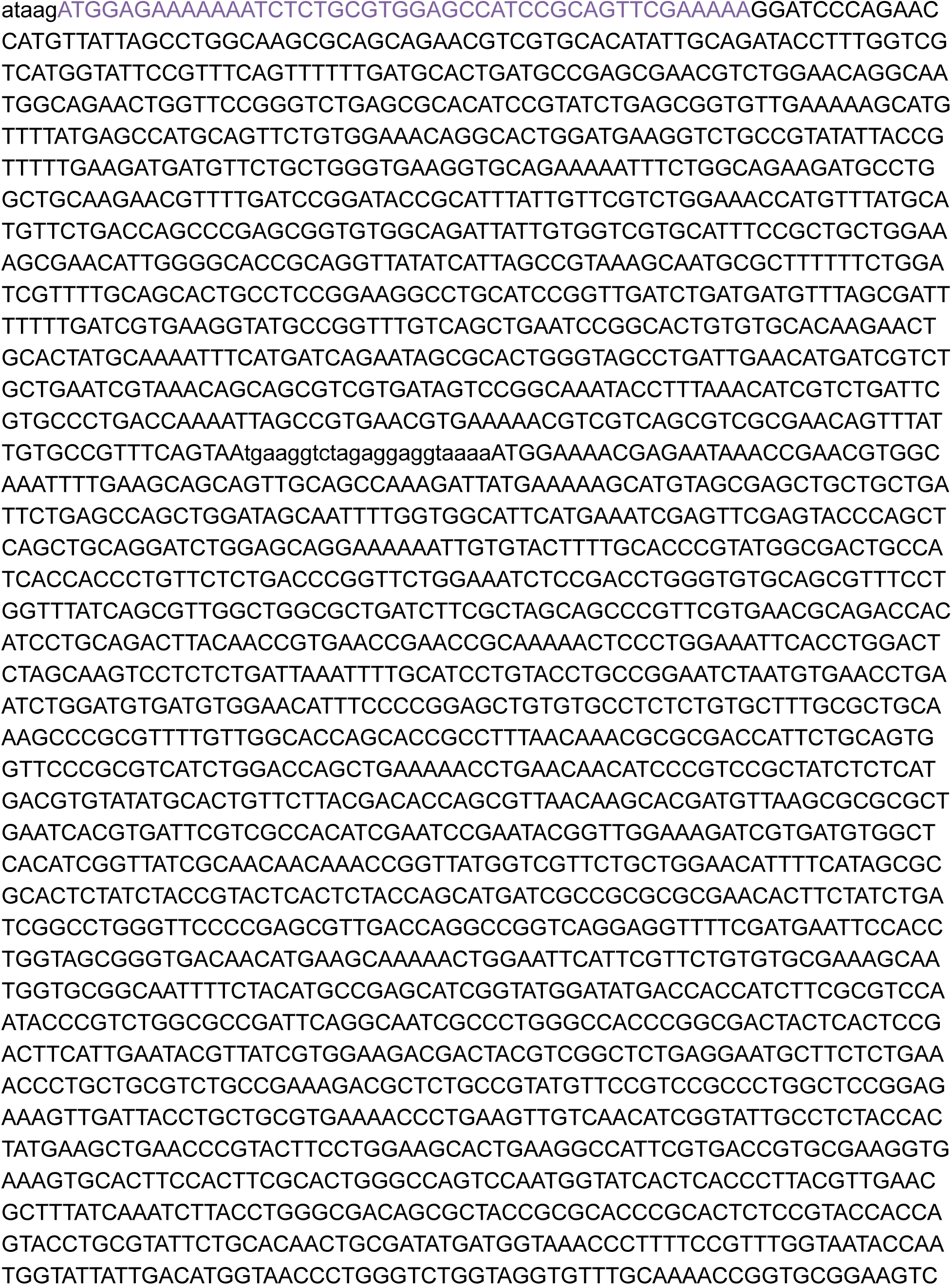

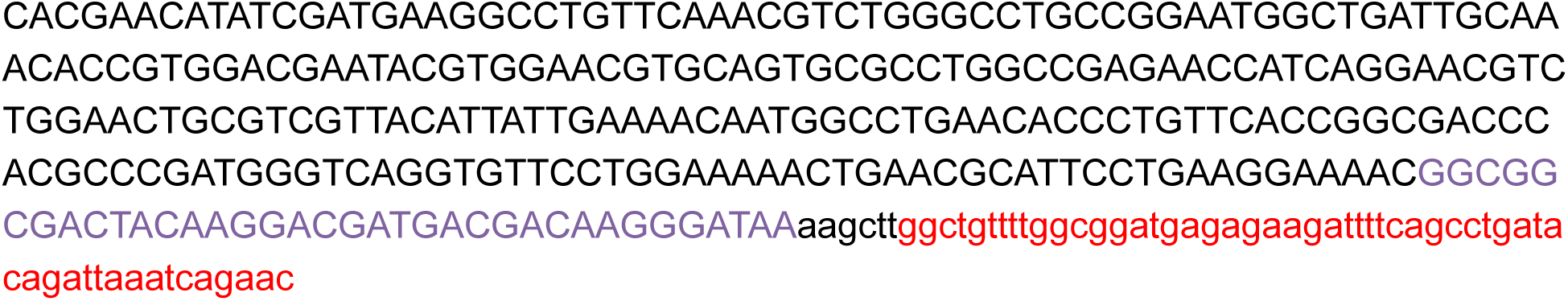

## DNA sequence of pCon.ConNeuA

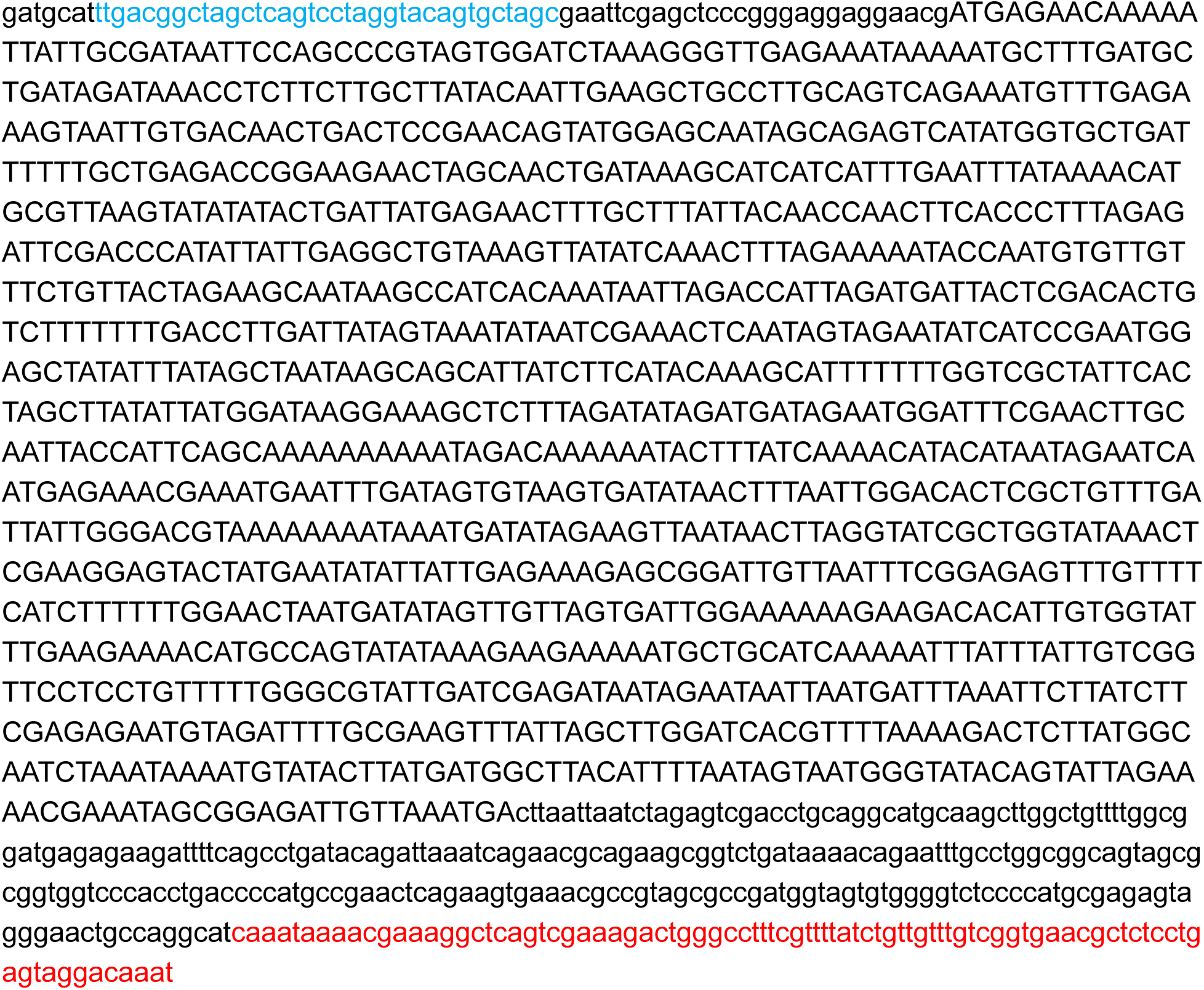

## Sequence of pBR322.Fc-6 in Context

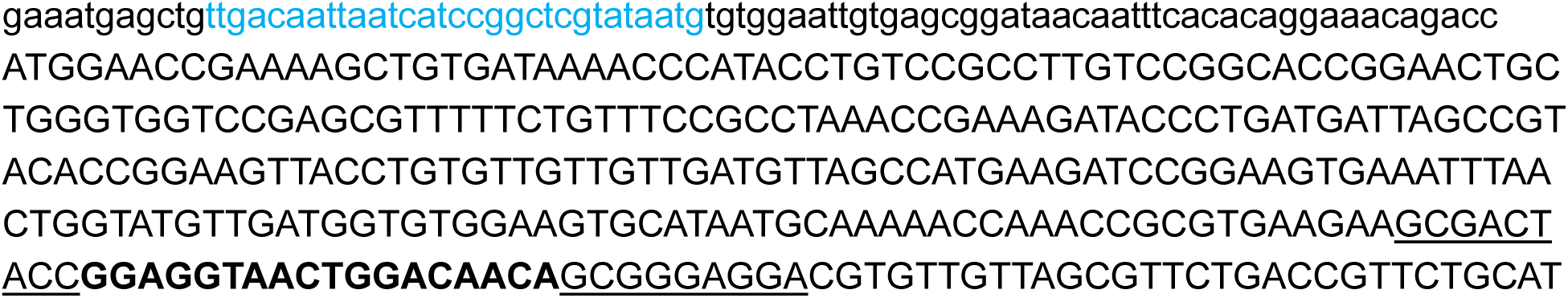

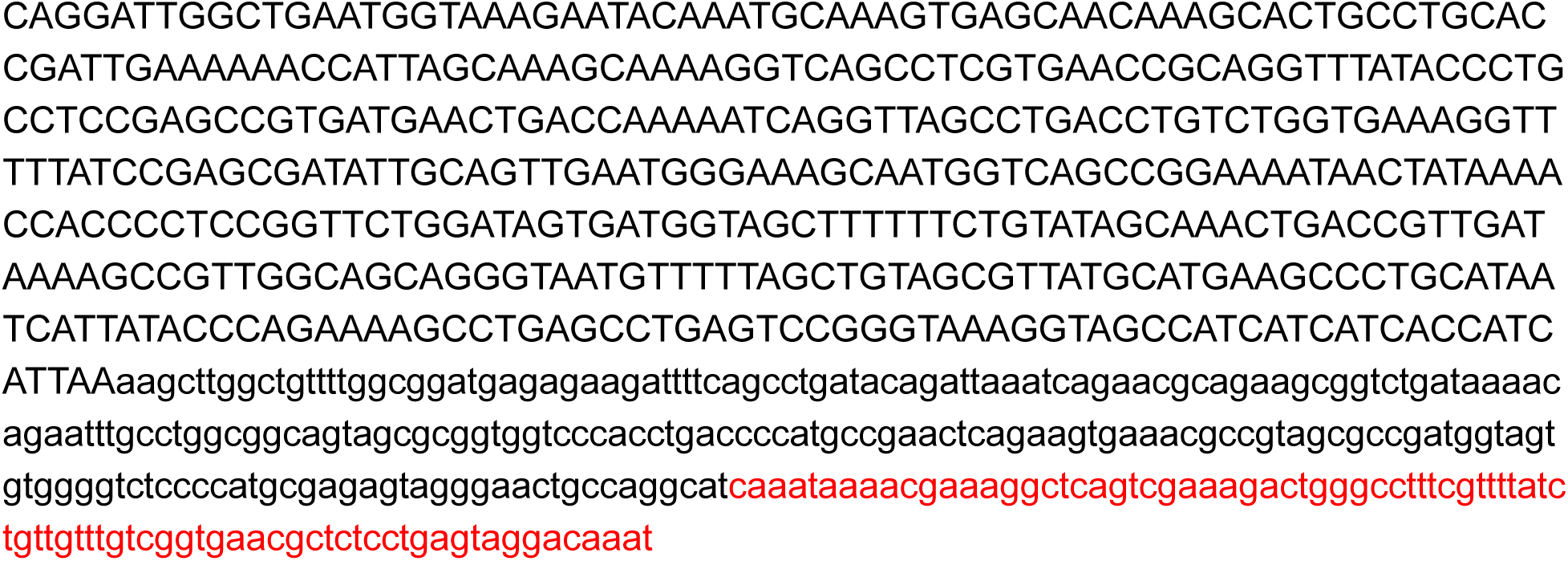

## Sequence of pBR322.Im7-6 in Context

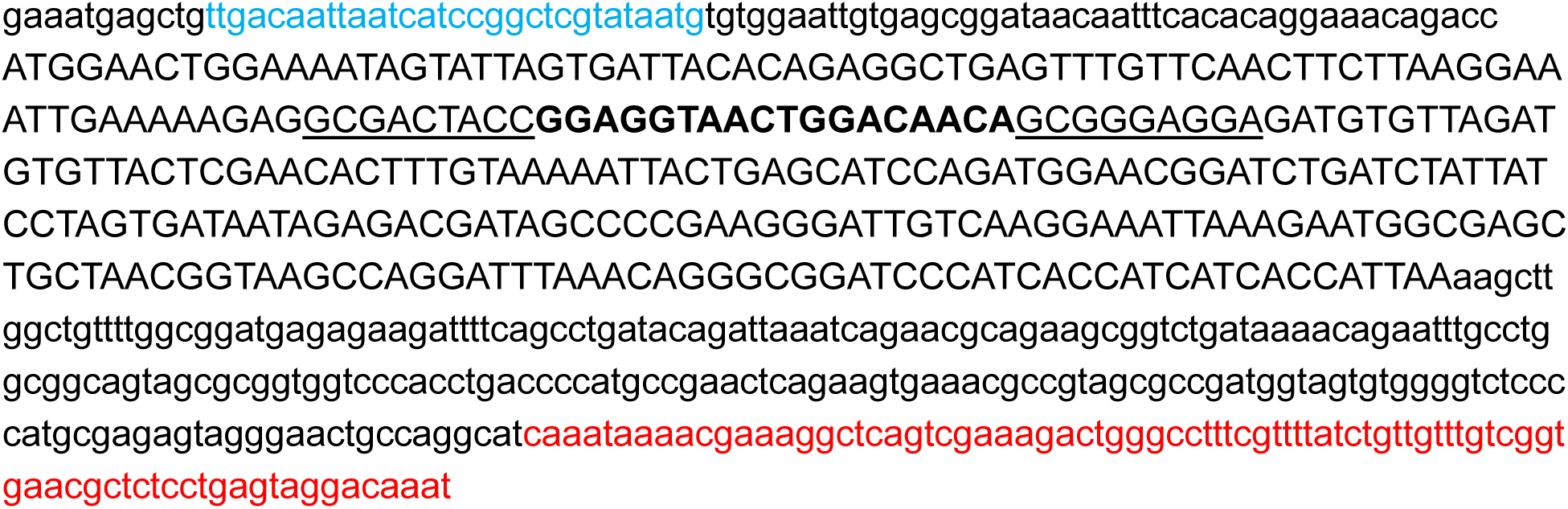

